# Keratin 17- and PKCα-dependent transient amplification of neutrophil influx after repeated stress to the skin

**DOI:** 10.1101/2023.10.11.561954

**Authors:** Yang Xu, Erez Cohen, Craig N. Johnson, Carole A. Parent, Pierre A. Coulombe

## Abstract

Neutrophils contribute to the pathogenesis of chronic inflammatory skin diseases. Little is known about the source and identity of the signals mediating their recruitment in inflamed skin. We used the phorbol ester TPA and UVB, alone or in combination, to induce sterile inflammation in mouse skin and assess whether keratinocyte-derived signals impact neutrophil recruitment. A single TPA treatment results in a neutrophil influx in the dermis that peaks at 12h and resolves within 24h. A second TPA treatment or a UVB challenge, when applied at 24h but not 48h later, accelerates, amplifies, and prolongs neutrophil infiltration. This transient amplification response (TAR) is mediated by local signals in inflamed skin, can be recapitulated in *ex vivo* culture, and involves the K17-dependent sustainment of protein kinase Cα (PKCα) activity and release of neutrophil chemoattractants by stressed keratinocytes. We show that K17 binds RACK1, a scaffold essential for PKCα activity. Finally, analyses of RNAseq data reveal the presence of a transcriptomic signature consistent with TAR and PKCα activation in chronic inflammatory skin diseases. These findings uncover a novel, transient, and keratin-dependent mechanism that amplifies neutrophil recruitment to the skin under stress, with direct implications for inflammatory skin disorders.

## Introduction

Inflammation and dysregulated immune responses are significant determinants of several skin-associated diseases, e.g., psoriasis (PSOR) (1, 2), atopic dermatitis (AD) (1), acne (3), and hidradenitis suppurativa (HS) (4, 5). Neutrophils normally act as essential effectors of innate immunity at sites of tissue injury, infection, and inflammation (6), but excessive neutrophil infiltration drives the progression and severity of skin diseases through various mechanisms (2, 7). Circulating neutrophils are recruited to sites of inflammation after sensing chemical cues in a process called chemotaxis (6). They usually are the first leukocytes to reach sites of tissue damage, infection or stress, and are tasked with defending the host by releasing proteases and reactive oxygen species, phagocytosing pathogens and cell debris, and releasing genomic DNA-histone complexes known as neutrophil extracellular traps (6). Neutrophils rarely occur in normal healthy skin, but readily infiltrate when skin is subjected to environmental stresses and in various inflammatory skin disorders, such as PSOR (1, 2), HS (4, 5), and cancer (7). Yet the kinetics of neutrophil infiltration and resolution and the mechanisms by which stressed skin tissue generates neutrophil-recruiting signals remain poorly understood.

Keratinocytes, the predominant cell type in the epidermis, are an important source of signals that regulate inflammation and immunity in skin (1, 8). These cells express abundant amounts of keratin proteins in a tightly regulated fashion (9). A distinct subset of keratins, including the type II keratins K6A-C paralogs and type I K16 and K17, are prominently involved in the response of epidermis to various stresses (10, 11). Expression of K6A-C, K16 and K17 occurs in ectoderm-derived epithelial appendages and in the thick epidermis of palms and soles in healthy human and mouse skin (12). Though absent from healthy interfollicular epidermis, K6A-C, K16 and K17 are rapidly induced after injury and many other stresses to the skin (11, 13), in chronic diseases such as PSOR (10), AD (14), acne (15) and in multiple types of carcinomas (16, 17). Besides, missense mutations in any one of the genes coding for K6A-C, K16 and K17 can cause pachyonychia congenita (PC) (18, 19), a rare disorder characterized by dystrophic nails, oral lesions, cutaneous cysts, and painful and debilitating palmoplantar keratoderma. The latter are lesions driven by dysregulated epithelial differentiation, redox balance, innate immunity and inflammatory pathways (20). As is the case for most intermediate filament (IF) proteins, the stress-responsive keratins fulfill classic roles for cytoskeletal proteins along with non-canonical roles in the cell. In a context-dependent fashion, for instance, the cytoplasmic pool of K17 regulates protein synthesis and keratinocyte growth (21, 22), TNFα-induced apoptosis (22) and inflammatory/immune gene expression (23) while a small nuclear pool of K17 regulates nuclear architecture (24), gene expression (25, 26), the cell cycle (24, 27), and the DNA damage and repair response (28). Though a role for K17 in altering inflammation and immune responses in skin experiencing stress has been established (25, 29), whether it regulates neutrophil infiltration, specifically, and the potential relevance for inflammatory skin diseases, are unknown.

Here we show that the skin transiently adapts to a recent stress exposure and exhibits a faster, stronger and prolonged neutrophil infiltration in response to additional stress exposures. This phenomenon, which we designate as TAR for transient amplification response, is short-lived and mediated by local, keratinocyte-derived signals. TAR entails a K17-dependent sustainment of the activity of protein kinase C alpha (PKCα), resulting in neutrophil chemokine release by stressed keratinocytes. Comparative analysis of transcriptomic datasets suggests that TAR, as observed in mouse skin under acute stress, is relevant to human inflammatory skin diseases.

## Results

### Transient amplification of neutrophil infiltration upon repeated stress to skin

Mouse ear skin was treated with 12-O-tetradecanoylphorbol-13-acetate (TPA), a phorbol ester widely used to induce sterile inflammation or as a tumor promoter (30). Skin tissue was harvested at different time points after single or dual topical TPA treatment (Tx) (**Fig. 1A**) and analyzed using immunostaining and CyTOF. After a single exposure, a neutrophil infiltrate is first observed in the dermis at 6h, peaks at 12h, and is cleared by 24h (**Fig. 1B**). When a second TPA Tx is given 24h later, the resulting neutrophil influx is significantly accelerated, amplified, and persists longer (**Fig. 1B**). CyTOF analysis of immune cells isolated from mouse ear tissue confirms the large increase in CD11b+Ly6g+ neutrophil infiltration 12h after a second TPA Tx, compared to 12h after a first Tx (**Fig. 1C**). We next tested whether the amplification of neutrophil infiltration represents a stably acquired property in TPA-Tx skin. When the second TPA Tx is given at 48h instead of 24h after the first Tx, the kinetics and amplitude of neutrophil recruitment replicate the response after a single Tx – i.e., there is no amplification (**Fig. 1D,E**). We refer to this phenomenon as a transient amplification response (TAR) whereby an earlier onset and more robust infiltration of neutrophils occurs upon repeated topical exposure to the chemical irritant TPA, in a manner that depends upon the length of interval between treatments.

**Figure 1.**
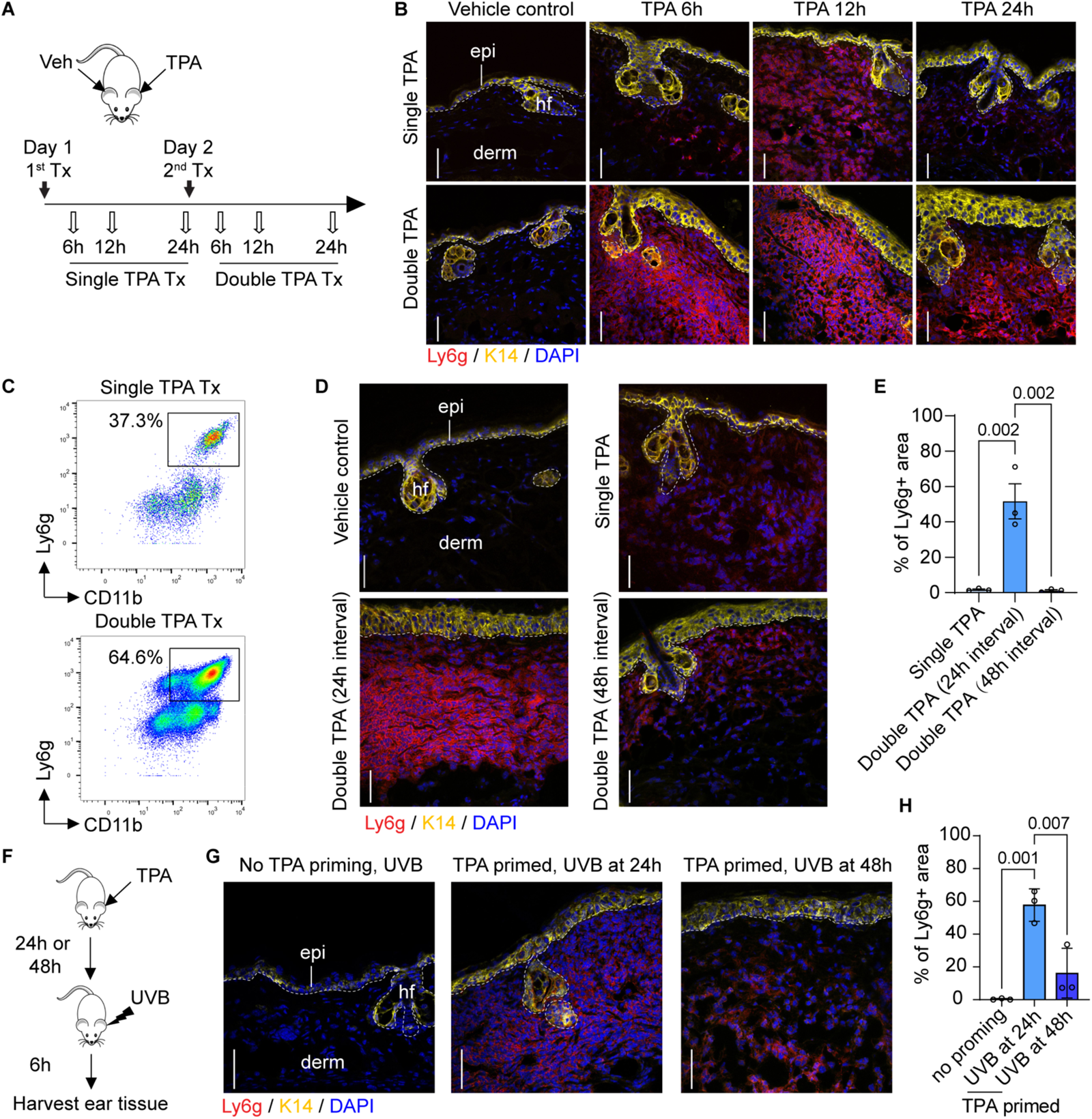
Transient amplified response (TAR) in skin subjected to repeated irritation. **A)** Strategy for single and double topical TPA Tx, 24h apart, and skin tissue sampling. **B)** Sections of WT mouse skin treated as shown in A were immunostained for K14, Ly6g, and nuclei (DAPI). epi, epidermis; derm, dermis; hf, hair follicle. Scale bars: 50μm. Dashed lines depict the dermo-epidermal interface. **C)** CyTOF analysis for CD11b (granulocytes, X-axis) and Ly6g (neutrophils; Y-axis) at 12h after a single and double TPA Tx of ear skin. Cells were pre-gated as single, live, non-beads, DNA- and CD45-positive. The Ly6g+CD11b+ neutrophil populations are boxed, and their frequencies indicated. n=3 mice. **D)** WT mouse skin was treated with acetone, single TPA, dual TPA (24h apart) or double TPA (48h apart). Skin sections were immunostained for K14, Ly6g, and nuclei (DAPI). Scale bars: 50μm. **E)** Quantitation of neutrophil fluorescence signal (percentage of Ly6G+ surface area in total dermis). n=3 mice. Data are shown as mean ± SEM. One-way ANOVA. **F)** Strategy for UVB and TPA combination Tx and skin tissue sampling. **G)** Sections of WT mouse skin treated as shown in F were immunostained for K14, Ly6g, and nuclei (DAPI). Scale bars: 50μm. **H)** Quantitation of neutrophil fluorescence signal (surface area measurements) of Ly6g. n=3 mice. Data are shown as mean ± SEM. One-way ANOVA.

TAR also takes place when the second external challenge to TPA-primed skin consists of an exposure to 400 mJ/cm^2^ UVB (**Fig. 1F**). Again, here, we found that the response to UVB as a second treatment entails an acceleration and increased amplitude of neutrophils in the dermis, and is manifested when UVB is administered at 24h, but not 48h, after the initial TPA Tx (**Fig. 1G,H**). Single or dual exposures of naïve skin to 400 mJ/cm^2^ UVB radiation does not cause a significant infiltration of neutrophils (**Fig. 1G**; data not shown). The latter is an indication that while TAR is not specific to TPA-induced irritation, it requires the presence of a component that is specifically induced by acute exposure to TPA, but not UVB, in mouse skin (see below).

### TAR is mediated by local signals and is not related to the barrier status in stressed skin

To assess whether TAR is related to the skin barrier status, we measured trans-epidermal water loss (TEWL) in dorsal ear skin treated with TPA. Significant and comparable increases in TEWL values occur at 24h and 48h after a single topical TPA Tx (**Suppl. Fig. 1A**), suggesting that differences in skin barrier status do not account for the difference seen in neutrophil infiltration when the repeat challenge is given at 48h vs. 24h after the initial insult. To examine whether TAR represents a systemic response after the initial stress, we performed studies in which sequential TPA Tx, 24h apart, are applied on the same versus contralateral ears (**Suppl**. **Fig. 1B**). We found that TAR only occurs when the same ear is exposed to dual TPA Tx (Suppl. Fig. 1C,D). These findings show that TAR is primarily driven by local signal(s) in treated skin and cannot be transferred by systemic circulating factors to naïve sites.

### Induced K17 in keratinocytes promotes neutrophil recruitment in stressed skin

Stress keratins are robustly induced in epidermal keratinocytes subjected to various challenges (11, 13). K17, in particular, is of interest for a potential role in TAR as it has been implicated in the immune response of TPA-Tx mouse skin and in several skin cancer models (23, 25, 28, 29). At baseline, K17 is expressed in skin epithelial appendages (e.g., hair follicles, sebaceous and sweat glands) but not in the interfollicular epidermis of mouse skin (**Fig. 2A**). K17 is induced within hours in the upper layers of epidermis after a single topical TPA Tx and is present in significant amounts when the skin is again exposed to TPA at later times (**Fig. 2A,B**). We next set out to assess whether TAR occurs in ear skin of *Krt17* null mouse (31). After a single TPA Tx, neutrophil infiltration parameters are indistinguishable in *Krt17* null and WT mice, suggesting that the constitutive pool of K17 in pilosebaceous units does not impact neutrophil infiltration in the dermis (**Suppl. Fig. 2A**, **Fig. 2C**). When a second TPA Tx (or UVB exposure) is given 24h later, however, neutrophil recruitment is significantly delayed in *Krt17* null skin (**Fig. 2B-E**). We also observed that intra-epidermal neutrophil infiltration occurs in *Krt17* null epidermis, but not in WT control, at 36h following dual TPA Tx spaced 24h apart (see white arrows in **Fig. 2B**). Of note, neutrophilic infiltration in the epidermis occurs in generalized pustular PSOR (32) and in HS (5). Though potentially interesting, this observation was not pursued further in the current study. We found no difference in the percentage of TUNEL-positive cells 6h after dual TPA-TPA or TPA-UVB Tx, spaced 24h apart, in *Krt17* null vs. WT mouse skin (**Suppl. Fig. 2B,C**), suggesting that the delay in TAR in *Krt17* null skin is not due to differences in apoptotic cell death. Together, these findings show that the induced pool of K17 in TPA-primed epidermal keratinocytes plays a significant role in the early amplification of the neutrophil influx induced by repeated stresses to skin.

**Figure 2.**
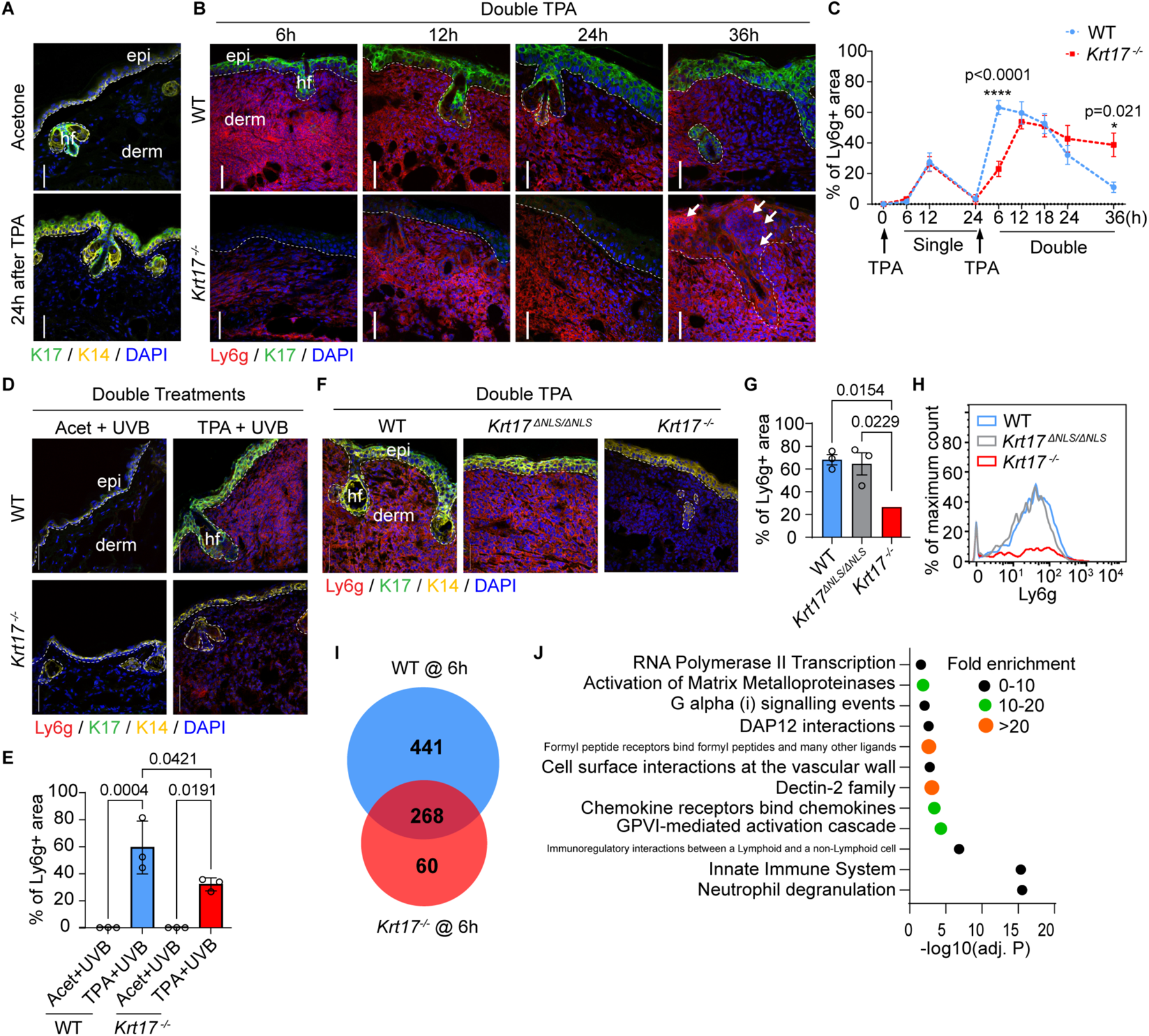
Role of induced K17 in epidermis of skin subjected to repeat topical challenges. **A)** Sections of WT skin treated with single acetone or TPA for 24h were immunostained for K14, K17, and nuclei (DAPI). Scale bars: 50μm. epi, epidermis; derm, dermis; hf, hair follicle. Dashed lines depict the dermo-epidermal interface. **B)** Sections of WT and *Krt17^-/-^* mouse skin treated with double TPA, 24h apart, immunostained stained for K14, Ly6g, and nuclei (DAPI). Time of sampling after treatment indicated above micrographs. Scale bars: 50μm. **C)** Quantitation of neutrophil fluorescence (surface area measurements), comparing WT (blue trace) and *Krt17^-/-^* (red trace) as a function of time after a single and double TPA Tx. n=3 mice. Data are shown as mean ± SEM. Two-way ANOVA. **D)** Sections of WT and *Krt17^-/-^* mouse skin treated with acetone or TPA, followed by UVB Tx 24h later, were immunostained for K14, K17, Ly6g, and nuclei (DAPI). Scale bars: 50μm. **E)** Quantitation of neutrophil fluorescence signal (surface area measurements) from D. n=3 mice. Data are shown as mean ± SEM. One-way ANOVA. **F)** Sections of WT, *Krt17^-/-^*, and *Krt17^ΔNLS/ΔNLS^* mouse skin treated with double TPA, 24h apart, were immunostained for K14, K17, Ly6g, and nuclei (DAPI). Scale bars: 50μm. **G)** Quantitation of neutrophil fluorescence (surface area measurements) from F. n=3 mice. Data are shown as mean ± SEM. One-way ANOVA. **H)** CyTOF analysis for Ly6g (neutrophils) at 6h after double TPA Tx (24h apart) of mouse ear skin. Cells were pre-gated as single, live, non-beads, DNA- and CD45-positive. The frequencies of Ly6g+ neutrophil populations are indicated. n=3 mice. **I)** Genes significantly upregulated after dual TPA Tx compared to dual acetone-Tx, 24h apart, in WT and *Krt17^-/-^* mouse skin (Cutoffs: FDR-adjusted P < 0.01, fold change > 8). **J)** Panther over-representation test using Reactome pathways (FDR-adjusted P < 0.05) for 441 genes significantly upregulated in double TPA-treated WT skin but not in Krt17^-/-^ skin. Acet, acetone.

A small pool of K17 occurs in the nucleus of tumor keratinocytes where it impacts nuclear architecture and chromatin organization (24), pro-inflammatory gene expression (25), and DNA damage and repair (28). To assess whether the nuclear-localized pool of K17 contributes to TAR, we subjected *Krt17^ΔNLS/ΔNLS^* mice, which harbor a mutated nuclear localization signal in K17 (24), to dual TPA Tx, 24h apart. Immunofluorescence staining of tissue sections (**Fig. 2F,G**) and CyTOF analysis of immune cell populations (**Fig. 2H**) show that the ear skin of *Krt17^ΔNLS/ΔNLS^* mice display a TAR that is indistinguishable from WT mice. Therefore, TAR is primarily regulated by the cytoplasmic pool of K17 in stressed epidermal keratinocytes.

We next performed bulk RNAseq analysis of WT and *Krt17* null mouse ear tissue obtained at 6h after a second TPA Tx to investigate TAR-related changes in gene expression. Volcano plots highlighting genes that are up- or down-regulated by 8-fold or more (adj. P<0.01) as a function of genotype (WT vs. *Krt17* null) and treatment (control vs. TPA) are shown in **Suppl. Fig. 2D,E**. We focused subsequent analyses on upregulated genes (**Fig. 2I**; see full listing in **Suppl. Table 4**; downregulated genes (see **Suppl. Fig. 2F**) are listed in **Suppl. Table 5**). A total of 441 genes were selectively upregulated in WT, 60 were selectively upregulated in *Krt17* null, and 268 were upregulated in both genotypes (**Fig. 2I**). To investigate which signaling pathways are affected by *Krt17* expression in TPA-induced skin inflammation *in vivo*, the 441 genes upregulated in WT only and 268 genes upregulated in both WT and *Krt17* null skin were separately analyzed using the Panther over-representation test and Reactome pathway tool (33, 34) (note: too few genes (n=60) were upregulated in *Krt17* null skin, specifically, to yield significant pathway findings). Several pathways are significantly over-represented (FDR-adjusted p value <0.05) in WT skin and in both WT and *Krt17* null skin (**Fig. 2J; Suppl. Fig. 2G**). The “innate immune system” and “neutrophil degranulation” were the pathways most significantly enriched in a K17-dependent fashion, and virtually all other pathways showing significant enrichment reflect neutrophil or innate immune processes (**Fig. 2J** and **Suppl. Fig. 2G**). Though there was overlap in pathway significance, there is a significant difference in individual genes showing upregulation in the K17-dependent vs. K17-independent groupings (**Suppl. Table 4**). Taken together, this unbiased transcriptomic screen provides independent support for a significant role for K17 in regulating the skin’s response to stress, the innate immune/neutrophil response in particular. Further insight gained from this RNAseq dataset is reported below.

### K17 regulates neutrophil recruitment by promoting release of CXCR2 and CXCR3 ligands

We next conducted studies using A431 human tumor keratinocytes in culture, which express K17 at baseline, to explore whether keratinocyte-derived signals can directly mediate neutrophil chemotaxis in a TPA-dependent manner. A *KRT17* null A431 variant (25) is available (**Suppl. Fig. 3A**). A neutrophil chemotactic response to conditioned medium (CM) collected from either parental or *KRT17* null cells was assessed in transwell migration assays (**Fig. 3A**). CM from TPA-treated parental cells markedly stimulated the chemotaxis of human primary neutrophils relative to CM from vehicle-treated cells (**Fig. 3B**). By contrast, the chemotactic response to CM obtained from TPA-treated *KRT17* null cells was significantly reduced (**Fig. 3B**). Serial dilution of CM from both cell types showed a dose-dependent neutrophil response, suggesting that the latter entails a receptor-ligand interaction (Suppl. Fig. 3B). Restoration of K17 expression by CMV-driven expression of a GFP-WTK17 fusion in TPA-Tx *KRT17* null A431 cells rescued neutrophil chemotaxis (Suppl. Fig. 3C). Together, these findings show that CM derived from TPA-stressed A431 keratinocytes induces neutrophil directional migration owing to the presence of specific chemotactic factor(s) produced and/or secreted in a K17-dependent manner.

**Figure 3.**
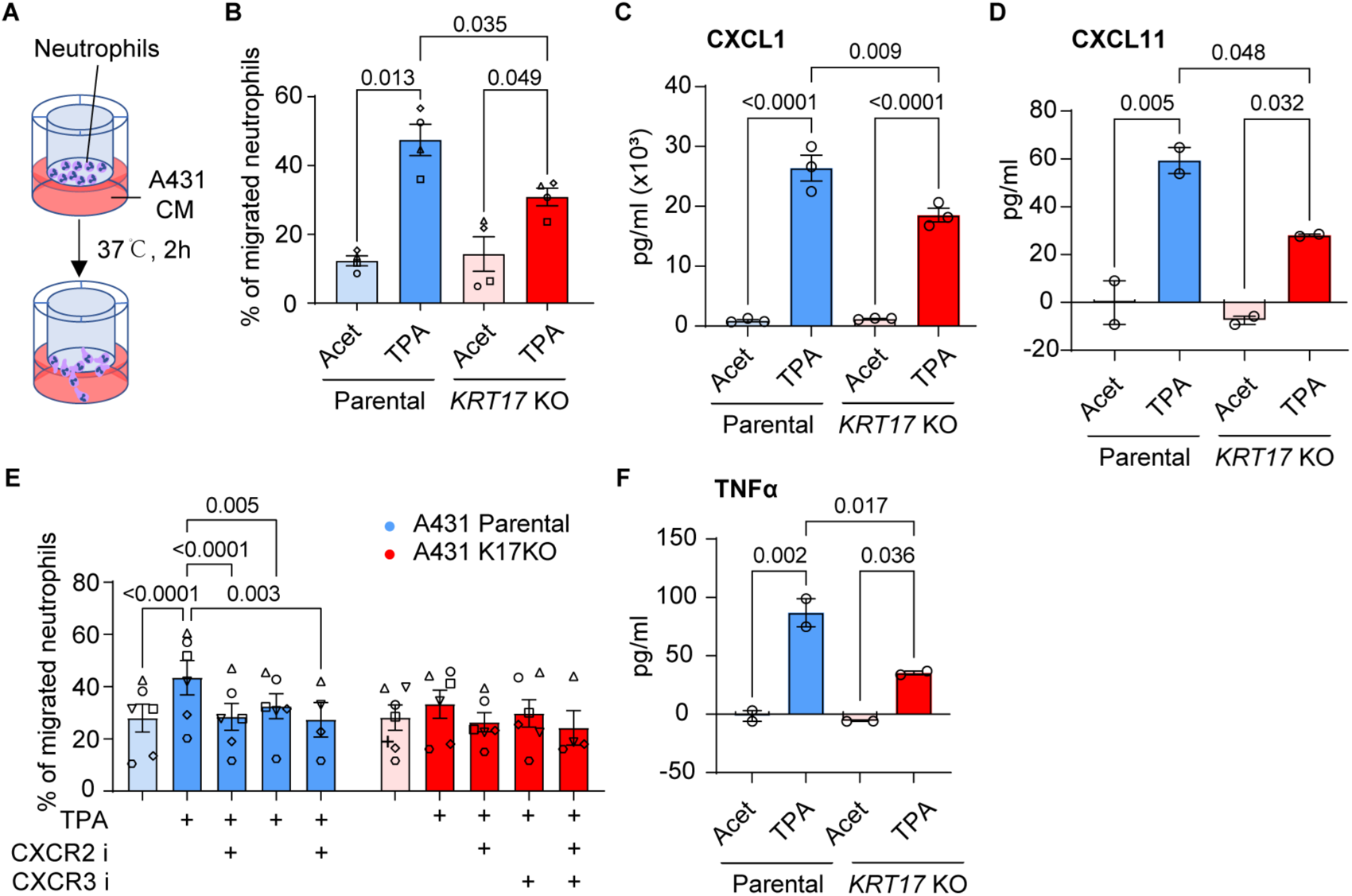
K17 promotes the secretion of neutrophil CXCR2 and CXCR3 ligands. **A)** Schematic of neutrophil transwell migration assay. **B)** Migration of human primary neutrophils towards conditioned medium (CM) from A431 keratinocyte cultures. Individual symbols depict data using neutrophils from different donors (n=4). Data are shown as mean ± SEM. One-way ANOVA. **C-D)** ELISA measurements of selected chemokine levels in A431 CM (pg/ml). N=3 measurements for CXCL1 (2 technical replicates each), n=2 measurements for CXCL11 (4 technical replicates each). Data are shown as mean ± SEM. One-way ANOVA. **E)** Migration of human primary neutrophils towards A431 CM in the presence of CXCR2 antagonist and/or CXCR3 antagonist. Individual symbols depict data using neutrophils from different donors (n=4-6). Data are shown as mean ± SEM. Two-way ANOVA. **F)** ELISA measurements of TNFα levels in A431 CM (pg/ml). n=2 measurements (4 technical replicates each). Data are shown as mean ± SEM. One-way ANOVA. Acet, Acetone.

ELISA assays were performed to compare the composition of CM collected from TPA-Tx parental and *KRT17* null A431 cells (Suppl. Table. 3). We prioritized analysis of secreted signals known to impact neutrophil migration, e.g., CXCL1 and CXCL2 (35), which act through CXCR2, and CXCL9 and CXCL11 (36, 37), which act through CXCR3. In addition to CXCR2, neutrophils under stress also express CXCR3 (see Suppl. Fig. 3D and refs (36, 37)) while CXCL9 and CXCL11 expression is dependent upon K17 in stressed human and mouse keratinocytes (25, 29). We found that CXCL1, CXCL2, CXCL9, and CXCL11 chemokines occur at higher levels in CM isolated from TPA-Tx parental relative to *KRT17* null A431 cells (these differences reached statistical significance for CXCL1 and CXCL11; see **Fig. 3C,D** and **Suppl. Fig. 3E,F**). We also found that significantly fewer neutrophils migrated towards CM derived from parental A431 cells pre-treated with antagonists for CXCR2 (AZD5069) (38) or CXCR3 (AMG487) (39), or both, prior to TPA Tx (**Fig. 3E**; antagonist specificity was confirmed using fMLF (Suppl. Fig. 3G)). These results indicate a role for CXCR2 and CXCR3 ligands and their receptors in mediating the keratinocyte- and K17-dependent recruitment of neutrophils.

### PKCα mediates the K17-dependent amplification of neutrophil influx in stressed skin

PKCα, the direct target of TPA, fulfills key roles in skin including the regulation of inflammatory responses and of desmosome cell-cell adhesion (40–42). Previous studies showed that massive overexpression of PKCα, driven by a strong keratin promoter in epidermis, leads to a severe neutrophilic inflammation in TPA-Tx skin (40, 42). We next assessed endogenous PKCα activity in WT mouse skin treated with TPA using an antibody against phospho-PKCα (Thr638), which reports on the active form of PKCα (43). The signal for phospho-PKCα (Thr638) is indeed increased at 1h after a single topical TPA Tx and has returned to baseline by 24h (**Figs. 4A,B**). Application of a second TPA Tx at that time (24h after the first Tx) results in a markedly higher signal for phospho-PKCα (**Figs. 4A,B**). By contrast, the signal intensity when using an antibody to total PKCα is unaffected by TPA Tx in WT mouse skin (**Suppl. Fig. 4A**). Next, we treated mouse ear skin with Gö6976, a pharmacological inhibitor of PKCα (44), prior to the second TPA Tx to test whether this kinase participates in TAR. Gö6976 Tx significantly curtailed TAR in WT skin but had no impact on the low-level infiltration of neutrophils in *Krt17* null skin (**Fig. 4C,D**). These findings suggest that PKCα and/or its downstream effectors play a key role in mediating TAR in skin tissue *in vivo*. We extended these *in vivo* findings by showing that pre-treatment of A431 keratinocytes with Gö6976 prior to TPA stimulation significantly reduced the ability of their CM to stimulate neutrophil chemotaxis (**Fig. 4E**). These findings show that endogenous PKCα activity mediates neutrophil infiltration in a K17-dependent manner in mouse skin and in human keratinocytes treated with TPA.

**Figure 4.**
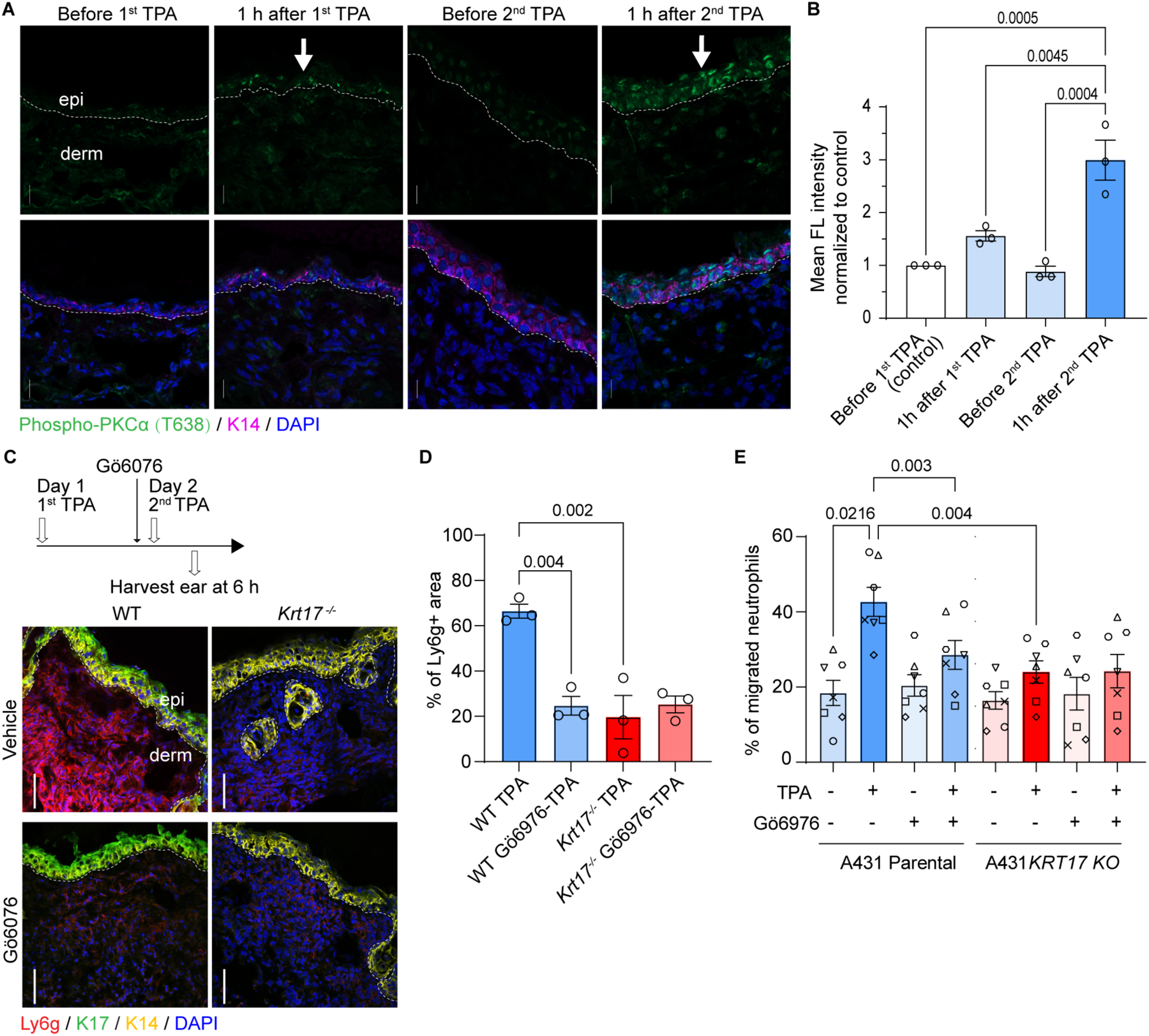
PKCα activity is crucial for K17-dependent neutrophil recruitment. **A)** Sections of WT skin before Tx, at 1 h after 1^st^ TPA Tx, before 2^nd^ TPA Tx (24h after the 1^st^ TPA), and at 1h after 2^nd^ TPA were immunostained for phospho-PKCα (T638), K14, and nuclei (DAPI). Scale bars: 50μm. epi, epidermis; derm, dermis. Dashed lines depict the dermo-epidermal interface. **B)** Quantitation of the signal for phospho-PKCα levels in epidermis from A. n=3 mice. Data are shown as mean ± SEM. One-way ANOVA. **C)** Impact of pre-treatment with Gö6976 at 30min prior to a 2^nd^ TPA Tx in WT and *Krt17^-/-^* mouse skin. Time interval between two TPA Tx was 24h. Sections were immuno-stained for K14, K17, Ly6g, and nuclei (DAPI). Scale bars: 50μm. **D)** Quantitation of neutrophil fluorescence signal (surface area measurements) from C. n=3 mice. Data are shown as mean ± SEM. One-way ANOVA. **E)** Migration of human primary neutrophils towards CM from A431 cells treated with TPA and Gö6976. Individual symbols depict data using neutrophils from different donors (n=7). Data are shown as mean ± SEM. One-way ANOVA.

We next wondered whether PKCα signaling is involved in eliciting TAR after sequential TPA and UVB Tx (see **Fig. 1**). PKCα activity levels was also significantly elevated in the epidermis of ear skin treated with TPA and UVB, 24h apart (**Suppl. Fig. 4B,C**). Pre-treatment with Gö6976 prior to UVB Tx in TPA-primed skin resulted in a significant reduction of neutrophil dermal infiltration (**Suppl. Fig. 4D**). This was confirmed by measurements of surface area of neutrophil infiltration in the dermis (**Suppl. Fig. 4E**) as well as of the dermal depth of the neutrophil infiltrate from the dermal-epidermal interface (**Suppl. Fig. 4F**). The latter index revealed that the epidermis-proximal pool of neutrophils was preferentially impacted by pre-treatment with Gö6976 in the setting of the TPA-UVB Tx (see white asterisks in **Suppl. Fig. 4D**). Accordingly, TAR does not require the sequential use of a direct PKCα activator such as TPA and may represent a general property of keratinocytes in acutely stressed skin.

### K17 regulates the subcellular localization and activity of PKCα in stressed keratinocytes

To investigate whether induced K17 participates in regulating PKCα in stressed keratinocytes, we compared the response of epidermis from WT and *Krt17* null mice to dual TPA Tx, 24h apart. Immunostaining of tissue sections for phospho-PKCα evidenced a weak signal in WT and *Krt17* null epidermis at baseline (data not shown; see **Fig. 4A,B**). After dual TPA Tx, the signal for phospho-PKCα is primarily concentrated in the epidermis and is significantly stronger in WT compared to *Krt17* null skin (**Fig. 5A,B**). By contrast, PKCα protein itself occurs at similar levels in WT and *Krt17* null epidermis after dual TPA Tx, based on immunostaining (**Suppl. Fig. 4A**). These data suggest that K17 may act as an amplifier of PKCα activity in stressed skin keratinocytes *in vivo*.

**Figure 5.**
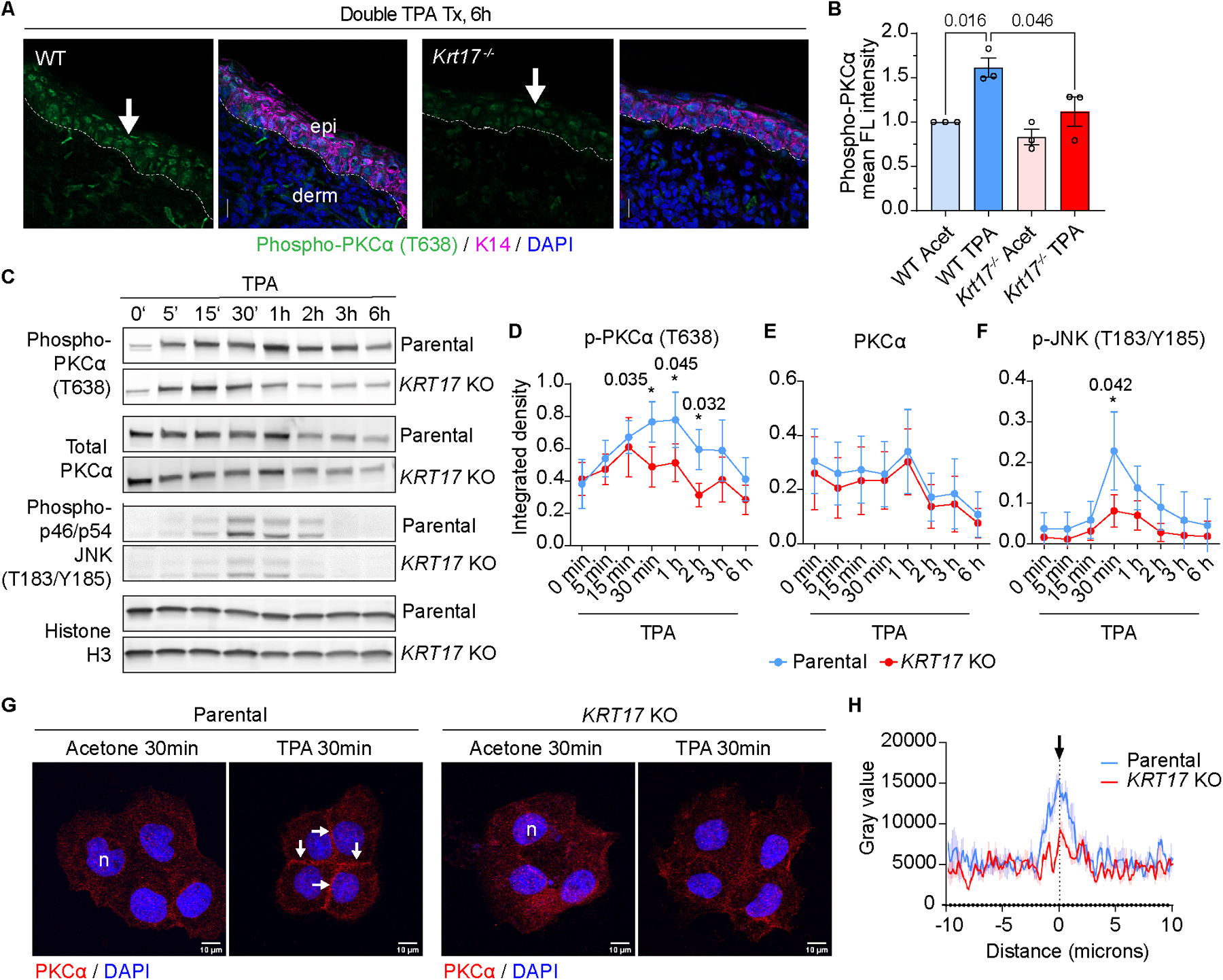
Subcellular localization and activity of PKCα are K17-dependent in A431 cells. **A)** Sections of WT and *Krt17^-/-^* mouse skin at 6h after double TPA Tx, 24h apart, were immuno-stained for phospho-PKCα (T638), K14, and nuclei (DAPI). Scale bars: 50μm. epi, epidermis; derm, dermis. Dashed lines depict the dermo-epidermal interface. **B)** Quantitation of the phospho-PKCα signal from micrographs as shown in A. n=3 mice. Data reported as mean ± SEM. One-way ANOVA. **C)** Western blot analysis of whole cell protein lysates at various times after TPA Tx (0-6h; see top of blots). **D-F**) Western blot quantitation of phospho-PKCα (T638), PKCα, and phospho-JNK (T183/Y185) normalized to histone H3, n=3. Data reported as mean ± SEM. Two-way ANOVA. **G)** Immunostaining for PKCα and nuclei (DAPI) in parental and *KRT17* KO A431 cells at 30min after vehicle or TPA Tx. Scale bars: 10μm. Arrows point to PKCα staining at cell-cell interface after TPA Tx. **H)** Quantitation of PKCα fluorescence intensity at cell-cell interface. n=16. Data reported as mean ± SEM. Acet, acetone; n, nucleus.

We took advantage of A431 keratinocytes, in culture, to continue probing the role of K17 in regulating PKCα activity. Time-course analyses of the activation of PKCα (via phosphorylation), using western immunoblotting, suggest similar kinetics in parental and *KRT17* null A431 cells during the first 15min after TPA addition (**Fig. 5C,D**). In parental A431 keratinocytes, phospho-PKCα levels continue to increase until 1h after TPA, then gradually decrease and have returned to baseline by 6h. By comparison, such a sustained increase in phospho-PKCα does not occur in *KRT17* null A431 cells, which show lower levels of phospho-PKCα at 30min, 1h and 2h after TPA addition. As was the case in mouse skin, total PKCα protein levels are indistinguishable across the same time frame in parental vs. *KRT17* null A431 cells (**Fig. 5C,E**). In further support of a difference between genotypes, phosphorylation of JNK, a direct target of classic PKC (45), also occurs at reduced levels in TPA-Tx *KRT17* null relative to parental A431 keratinocytes (**Fig. 5C,F**). Finally, using immunostaining, we find that translocation of PKCα to the cell membrane, a hallmark of its activation (46), occurs in TPA-treated parental A431 cells but significantly less so in *KRT17* null cells (**Fig. 5G,H**). Taken together, these findings demonstrate a role for K17 in sustaining PKCα activity after exposure of keratinocytes to stress.

PKCα regulates cell-cell adhesion in epidermal keratinocytes by phosphorylating desmoplakin, leading to detachment of keratin IFs and destabilization of desmosomes (41). Accordingly, we examined the status of desmosomes in TPA-Tx A431 cells in culture. Parental and *KRT17* null cells subjected to acetone vehicle Tx show indistinguishable desmoplakin and E-cadherin staining which, as expected, is concentrated at cell-cell junctions (**Suppl. Fig. 5A**). A partial loss of desmoplakin staining at cell-cell junctions occurs at 1h after TPA Tx in parental A431 cells (**Suppl. Fig. 5A**). By comparison, desmoplakin staining is slightly reduced at cell-cell interface at 30 min and definitely so at 1h after TPA Tx of *KRT17* null A431 cells (**Suppl. Fig. 5A**). The latter occurs despite attenuated PKCα activity in *KRT17* null cells (see **Fig. 5**), suggesting that desmosomes are impacted by PKCα-independent mechanisms in the absence of K17 (47).

### K17 interacts with the essential PKCα scaffolding protein RACK1

The scaffolding protein RACK1 binds classic PKC isozymes and is essential for their activation and enhanced partitioning to the outer cell membrane (48, 49). Kroger *et al.* showed that RACK1 physically interacts with K5, a type II keratin, in mouse keratinocytes (41) and Yang *et al.* reported the presence of RACK1 in K17 immunoprecipitates from human HaCaT cells (50). We confirmed that RACK1 co-immunoprecipates with K17 in protein lysates from A431 keratinocytes, whether treated with TPA or not (**Fig. 6A**), and that steady state levels for RACK1 protein are similar in parental and *KRT17* null A431 cells (**Fig. 6B**). PLA assays show that the proximity of RACK1 and K17 is robust in parental A431 cells at baseline (acetone control), and is enhanced ∼2.5 fold (P<0.02) at 30 min after TPA Tx (**Fig. 6C,D**). PLA assays also show enhanced proximity between RACK1 and phospho-PKCα (P<0.0001; **Fig. 6E,F**), and between K17 and PKCα (P<0.0001; **Fig. 6G,H**) at 30 min after TPA Tx in parental A431 cells, but both are significantly reduced in *KRT17* null A431 cells. We conclude that K17 is bound to RACK1 in cultures of A431 keratinocytes at baseline but their interaction markedly changes early after TPA Tx, coinciding with enhanced proximity between RACK1 and active PKCα.

**Figure 6.**
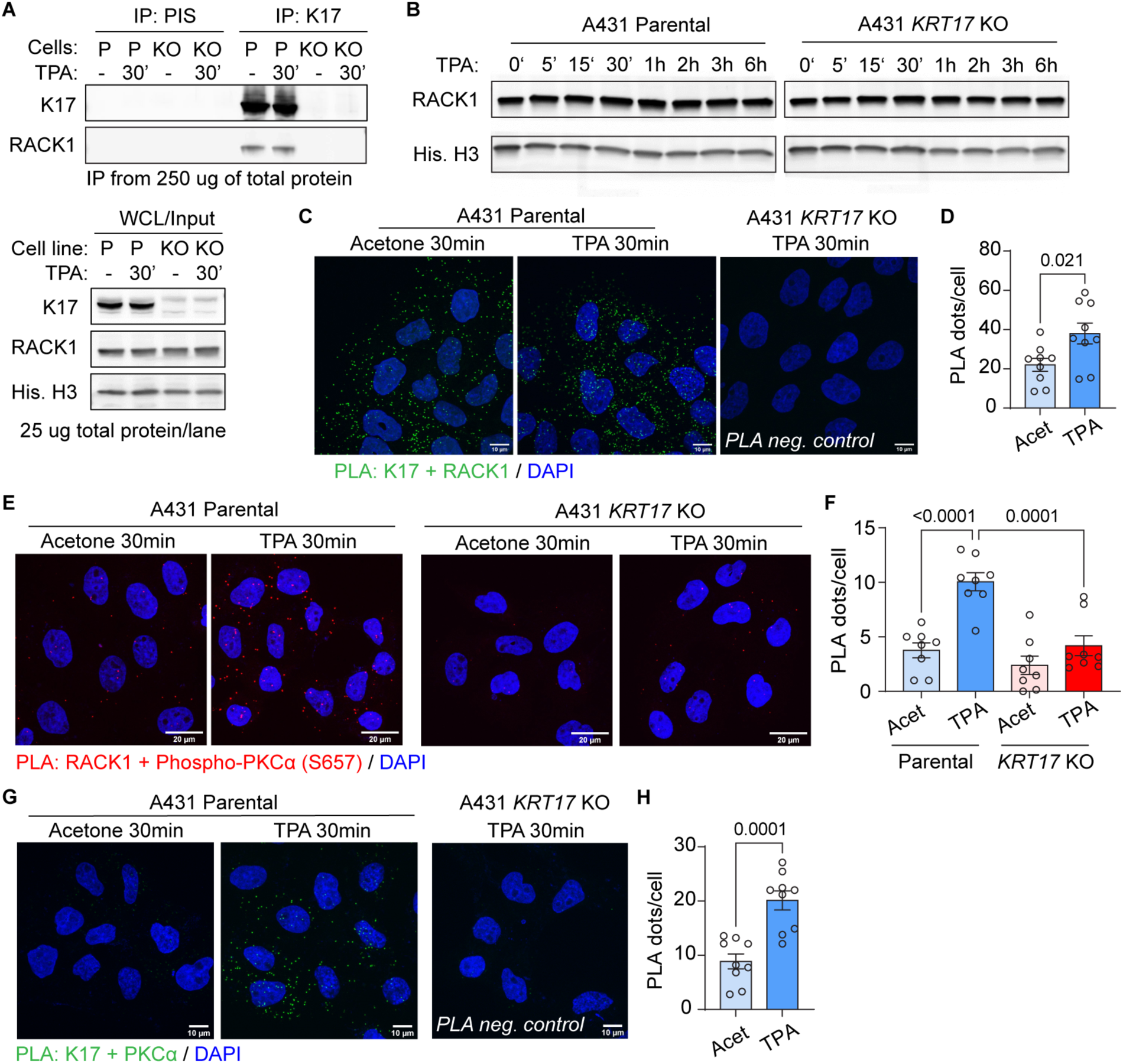
K17 interacts with the PKCα scaffolding protein RACK1. **A)** RACK1 occurs in K17 immunoprecipitates (IPs) from detergent-soluble protein extracts prepared from parental and *KRT17* null A431 cells. Rabbit preimmune serum (PIS) was used as IgG control for IP, and input protein content is reported in bottom gel. **B)** Western blot analysis of RACK1 level in parental and *KRT17* null A431 cells before and after TPA Tx. **C-H)** PLA assays were performed for (C) K17 and RACK1, (E) RACK1 and phosphor-PKCα (S657), and (G) K17 and PKCα in A431 keratinocytes treated with vehicle or TPA for 30min. Maximum intensity projection (MIP) images are shown. Scale bars: 10μm in C and G, or 20μm in E. In **D**, **F**, and **H**, PLA signals were quantified as the total number of PLA dots in a microscopic field divided by the number of nuclei in that same (field). n=8 or 9 fields. Data are shown as mean ± SEM. Unpaired t tests were performed in D and H and one-way ANOVA was done in F. Acet, acetone.

Keratins behave as insoluble proteins in aqueous salt solutions but are readily solubilized in buffers containing a high content of strong denaturing agents such as urea (51). Besides, the solubility of keratin proteins is naturally affected by post-translational modifications, the most studied of which is phosphorylation (52). We next applied a Triton X-100-based fractionation protocol (53) to monitor the partitioning of K17, RACK1 and PKCα proteins in parental A431 keratinocytes after Tx with TPA. K17 is most abundant in the detergent-insoluble fraction, as expected, but also occurs in low salt buffer and detergent-soluble fractions (**Suppl. Fig. 5B**). At 1h after TPA Tx the Triton-soluble pool of K17 is slightly increased (**Suppl. Fig. 5B**), likely reflecting a remodeling of K17-containing filaments (54). Significant amounts of RACK1 and PKCα are extracted by the low salt buffer, as expected, while RACK1 but not PKCα also occurs in the Triton-soluble pool (**Suppl. Fig. 5B**). Interestingly, we observed that a small amount of RACK1 and PKCα co-partitions with K17 to the keratin-rich detergent-insoluble pool (**Suppl. Fig. 5B**). These findings are consistent with the PLA data but do not conclusively inform on the pool of K17 that interacts with RACK1 and regulates PKCα.

### The N-terminal head domain of K17 regulates PKCα activity

In an effort to map cis-acting determinants in K17 that mediate regulation PKCα activity, we exploited the observation that expression of a GFP-WTK17 fusion (25) in *KRT17* null A431 cells rescues the TPA-induced recruitment of PKCα to the plasma membrane (PM) in TPA-treated transfected cells (**Fig. 7A,B**, note that proximal A431 null cells not expressing GFP-WTK17 failed to show PKCα recruitment to the PM in this setting). Expression of GFP-WTK14, which is highly related to K17 in primary structure (12) and constitutively expressed in epidermis, led to filament formation but did not rescue the PM localization of PKCα in transfected and TPA-Tx *KRT17* null A431 cells (**Fig. 7A,B**). Next, we constructed K14-K17 chimeras and tested them in this rescue assay to map the domain(s) within K17 that confers PKCα activation. We found that expression of a GFP-K17head-K14rod-K14tail chimera, but not a GFP-K14head-K17rod-K17tail chimera, succeeded in rescuing PM localization in transfected and TPA-Tx *KRT17* null A431 cells (**Figs. 7C,D**). Thus, the N-terminal head domain is key to K17’s ability to regulate PKCα.

**Figure 7.**
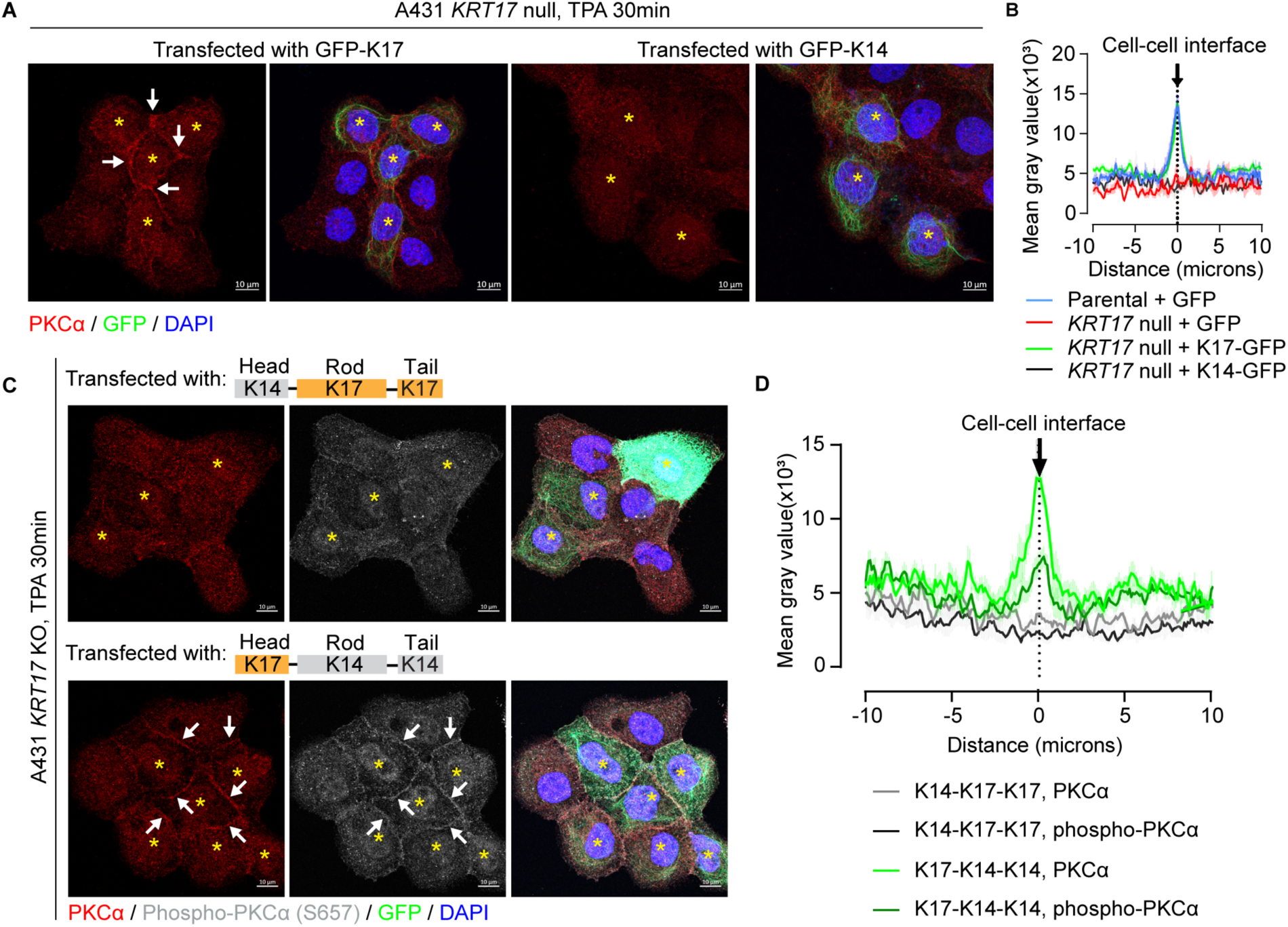
The N-terminal head domain of K17 is crucial for PKCα regulation. **A)** Rescue assays involving expression of GFP-WTK17 and GFP-WTK14 fusion proteins in *KRT17* KO A431 cells. Cells were treated with TPA for 30min and were stained for PKCα and nuclei (DAPI). Examples of transfected cells are identified by asterisks. Scale bars: 10μm. **B)** Quantitation of PKCα fluorescence intensity across cell-cell interfaces of adjoining transfected cells shown in A. n=12 cell for parental + GFP (control); n=16 for *KRT17* KO + GFP (control); n=22 for *KRT17* KO + K17-GFP; n=24 for *KRT17* KO + K14-GFP. Data are shown as mean ± SEM. **C)** Expression of GFP-K17head-K14rod-K14tail and GFP-K14head-K17rod-K17tail chimeric fusion cDNAs in *KRT17* KO A431 cells. Cells were treated with TPA for 30min and immunostained for PKCα and phospho-PKCα (S657), and for nuclei (DAPI). Scale bars: 10μm. **D)** Quantitation of PKCα and phospho-PKCα (S657) fluorescence intensity across cell-cell interfaces in adjoining transfected cells as shown in C. n=20 for K17-K14-K14 group, and n=16 for K17-K14-K14 group. Data reported as mean ± SEM. In **A and C**, arrows depict staining for PKCα and phosphor-PKCα at the plasma membrane in adjoining cells expressing GFP-WTK17- or GFP-K17-K14-K14.

### Relevance to chronic human inflammatory skin diseases

We next explored whether one can uncover a transcriptional signature consistent with TAR, as seen in mice, in human skin disorders with a known neutrophil involvement (see below). We focused on upregulated genes (>8-fold; adj. P<0.01) in WT but not *Krt17* null skin at 6h after dual TPA Tx, 24h apart. Of the 441 genes in this group (see **Fig. 2I**), 289 have human homologs and 209 occur in a well-curated PSOR single-cell (sc) RNAseq dataset (55) (**Suppl. Fig. 6A** and **Suppl. Table 6**). Analysis of these 209 genes using the Panther statistical over-representation test (33, 34) indicated an enrichment for genes involved in neutrophil function and regulation, e.g., Dectin-2 family (56), GPVI proteins (57), and cytokine and chemokine production (**Suppl. Table 7**). Most of these 209 genes show preferred expression in clusters typified by myeloid lineage markers (LYZ), CD16, and additional neutrophil markers (‘Myeloid-lineage’ cluster) (**Suppl. Fig. 6B**). This said, a small subset of 12 genes shows higher expression in skin keratinocytes over the myeloid-lineage cells (*ANGPTL4, IFITM1, IFI16, PHLDA2, KLK1, PDPN, AKR1B10, ACAT2, TUBB6, OAS1, C19orf48, HRH2*) (**Suppl. Fig. 6C**). Six of these genes, *AKR1B10* (58), *ANGPTL4* (59), *OAS1* (60), *HRH2* (61), *KLK1* (62) and *PDPN* (63), remarkably, have known roles in regulating PKCα or other classic PKCs, or in steps downstream from activated PKCα. Using these 12 genes we devised a composite expression score (TAR12 score) and used it to query PSOR (55), AD (64) and HS (4) scRNAseq datasets (see Methods and ref. (10)). The distribution of TAR12 scores in single cells of lesional PSOR (**Fig. 8A**) and HS skin (**Fig. 8D**) is quasi-normal. Cells with the highest TAR12 scores (cut-off set at either the 95^th^ and 75^th^ percentiles) also show the highest levels of reads for inflammatory, immune response and stress response genes (data not shown), including the highest levels of *KRT6A* and *KRT17* mRNAs (**Fig. 8B,C** and **Fig. 8E,F**) A similar outcomes is observed in a scRNAseq dataset from AD patients (**Suppl. Figs. 6D-F**). These findings are significant in three ways. They show that: (i) the transcriptional signature of WT mouse ear skin when TAR is first manifested (at 6h after two TPA Tx, 24h apart) shows a strong neutrophil character; (ii) several of the genes upregulated in TAR-exhibiting mouse skin are expressed in human inflammatory skin disorders (PSOR, AD, HS) and involved in regulating PKC activity; and (iii) a transcriptional signature inferred from TAR in mouse skin is shared with keratinocytes exhibiting high stress keratin expression, including *KRT17*, in skin disorders.

**Figure 8.**
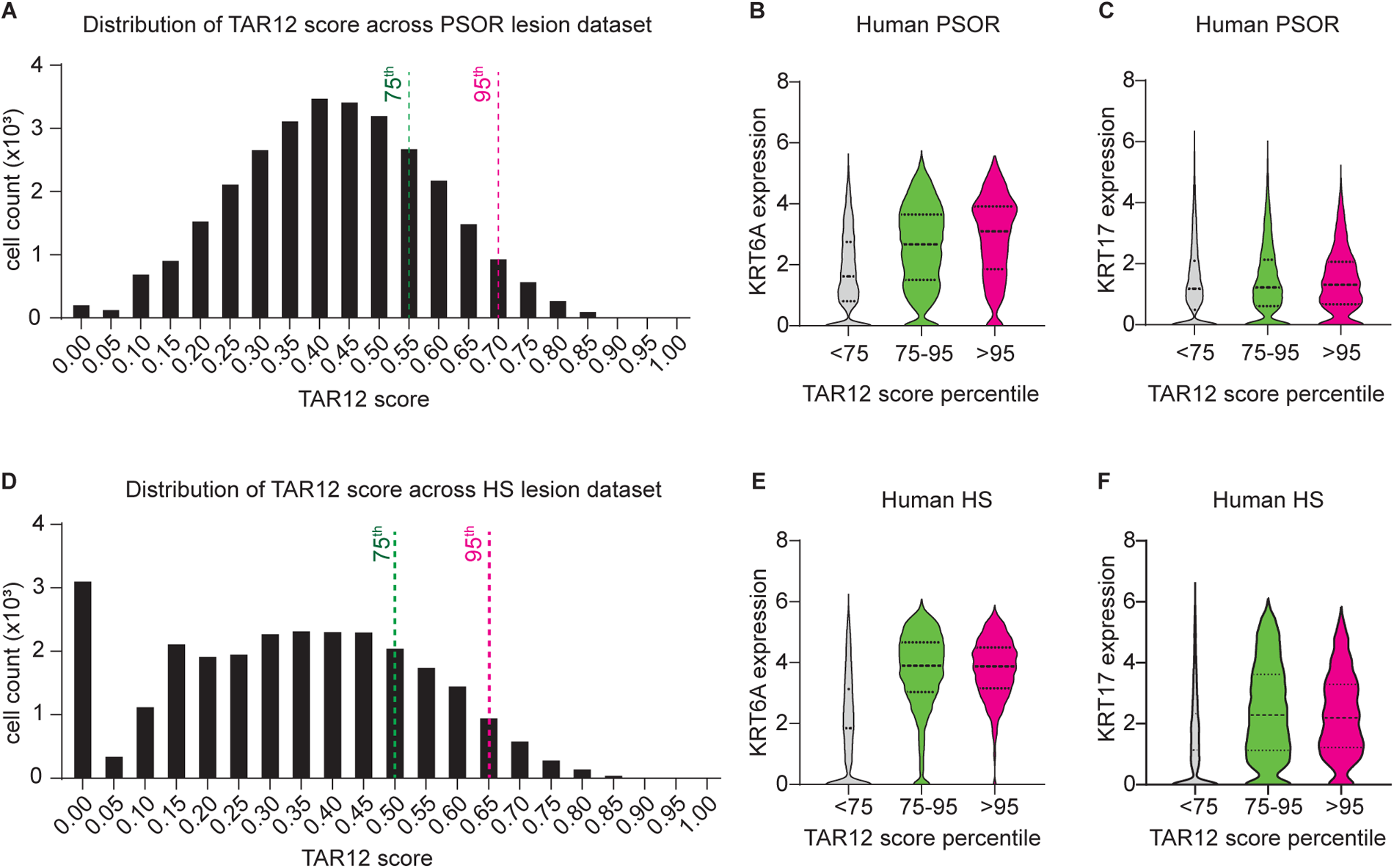
A TAR transcriptional signature occurs in stressed keratinocytes from human inflammatory skin diseases. A composite score, TAR12, was devised based on integrated analyses of transcriptomics datasets from TPA-Tx WT mouse skin and from human psoriasis (see main text) and used to query single cell RNA sequencing datasets from (A-C) psoriasis (PSOR) and (D-F) hidradenitis suppurativa (HS) in the human. **A, D)** Distribution of TAR12 scores in all individual cells from (A) PSOR and (D) HS. **B, C, E, F)** Violin plots reporting on expression levels for (B,E) *KRT6A* and (C,F) *KRT17* in cells showing high, medium, and low TAR12 scores. Cut-offs were arbitrarily set at the 95th percentile for a high TAR score and at the 75th percentile for a medium TAR score based on the distributions shown in A and D.

## Discussion

We show here that the cytoplasmic pool of K17 induced by stress in epidermal keratinocytes amplifies neutrophil recruitment in the superficial and deeper dermis upon additional stress exposures. This property represents a short-lived, transient adaptation that entails the K17-dependent regulation of PKCα activation and release of classic neutrophil chemokines. Analysis of transcriptomic data suggests that the observations made in acutely stressed mouse skin may be relevant to human chronic skin inflammatory diseases such as PSOR, AD and HS. These findings and their implications significantly advance our understanding of the relevance and roles of stress keratins, PKCα, and their partners in complex epithelia such as skin (10).

Classic examples of adaptation to stress include the photoprotective pigmentation that follows UV radiation in skin (65) and adaptive and innate immune cell responses (66). Recently, a new mechanism has been described whereby skin epithelial stem cells develop a stable epigenetic memory of past exposures to stress (67–69). This memory persists for >6 months after an initial insult, entails a Stat3- and Fos/Jun-dependent open chromatin state in epithelial stem cells, and enhances the skin’s sensitivity to subsequent stimuli (67, 68). The TAR described here is distinct as it is driven by epidermal keratinocytes, is short-lived (<48h), and regulated in part by K17-dependent PKCα signaling at the cell periphery. We envision that TAR may serve the purpose of transiently protecting skin against additional stress exposures during a period of enhanced keratinocyte vulnerability. Of interest, others have shown that AP-1 (70) and Stat3 (71) regulate *KRT17* transcription, and that the nuclear pool of K17 protein modulates Stat3 function in keratinocytes of inflamed skin (26). Conceivably, therefore, stresses that induce short-term TAR in differentiating keratinocytes of the epidermis may trigger a longer-lived epigenetic memory in epithelial stem cells of the tissue.

In the course of this study, we explored the ability of various types of acute stresses (e.g., TPA, UVB, UVA, Imiquimod) to elicit a TAR-like response in mouse ear skin. These efforts showed that TAR manifests only when TPA is applied as the initial insult. The latter correlates with the observation that, of the stressors tested, only TPA results in a robust and prolonged (up to 48h) induction of K17 in treated epidermis within hours after a single exposure. TAR does not occur when the second stress (TPA or UVB) is administered at 48h, instead of 24h, after the initial TPA priming event. While necessary, therefore, K17 is not sufficient to mediate the early phase of TAR. Transient factors present 24h but not 48h after a priming event may be required for TAR to develop in full. A post-translational modification(s) (PTMs) on K17, for instance, could be such a factor. PTMs can act *in cis* or *trans* to affect protein structure and/or interaction with other proteins (52). The N-terminal head domain of K17, which as we showed here is required for TPA-induced PKCα activation, undergoes phosphorylation on several Ser/Thr residues in response to TPA and other stimuli (e.g., Thr9, Ser10, Ser13, Ser32, Ser44 (21, 22, 72)). Ser44 phosphorylation on K17 fosters its interaction with 14-3-3σ and enables the resulting complex to promote protein synthesis and cell growth in keratinocytes challenged to grow (21). Moreover, the K17 Ser44 modification persists for only a few hours after a single TPA exposure (22). We did test a K17 Ser44Ala mutant available to us (21) and found that, like WT K17, it rescues the PKCα phenotypes of *KRT17* null A431 keratinocytes (data not shown). K17 is also subject to SUMOylation (73) and ubiquitination (50, 72), so other PTMs (and other protein partners) could have a positive influence on TAR. Alternatively, it could be that a negative regulator of TAR is present or effective only at 48h beyond the initial, K17-inducing priming event in skin. Solving this enigma requires additional studies but is poised to yield novel biomarkers and/or clues as to how to mitigate TAR in the setting of human chronic inflammatory skin diseases.

We found that K17 promotes the membrane recruitment and activity of PKCα in epidermal keratinocytes subjected to inflammation-inducing stress, both in cell culture and *in situ*. This newly defined property involves specific *cis*-acting determinant(s) located in the N-terminal head domain of K17 (12) and RACK1, a scaffolding protein essential for PKCα activity (48, 49). PKC regulates several signaling events associated with the pathogenesis of inflammation in various cell types, including nitric oxide biosynthesis (74), inflammatory cytokine (75) and superoxide production (76), and phospholipase A2 activation (77). Increased levels of diacylglycerol (78) and of PKCα (79) occur in psoriatic skin, consistent with a role for this kinase in regulating skin inflammation. Kroger *et al.* (41) reported that K5-K14 filaments promote the formation of calcium-induced, stable adhesions between keratinocytes by physically sequestering PKCα away from desmosomes, and showcased K5-K14 as negative regulator of PKCα activity. Our data shows that K17 is a positive regulator of both the membrane association and activity of this kinase. These differences highlight the need for deeper studies of the mechanism and regulators of the keratin-RACK1-PKCα signaling axis in epidermal keratinocytes.

Our findings add to evidence linking all subclasses of IFs to the regulation of cellular kinases, with direct significance for various cellular processes. Thus K6, a type II keratin that is partially co-regulated with *KRT17*/K17 in stressed and diseased skin (10), directly binds Src, inhibiting its activity and promoting the coordination of migration and adhesion within keratinocyte sheets during wound re-epithelialization (80, 81). K8/K18, a keratin pair prominent in simple epithelia, binds to Akt1 stimulates its phosphorylation-dependent activation, and promotes hepatocyte survival after liver injury (82). K10, a type I keratin expressed at an early stage in differentiating keratinocytes of epidermis, also binds Akt, leading to sequestration of the kinase and promoting cell cycle arrest (83). Vimentin, a type III IF, binds directly to CaMKII in smooth muscle cells and sequesters it to regulate contractility. Nestin, a type IV IF, binds directly to and sequesters Cdk5 to promote survival of neuronal progenitor cells (84) and also binds Plk1 to positively regulate its phosphorylation and smooth muscle contraction (85). The K17-PKCα and, more generally, the IF-kinases partnership provides compelling examples of the growing role of IFs in regulating key signaling pathways in a biological context-dependent manner (10, 17, 20, 82).

The uncovering of a transient amplification of neutrophil infiltration in skin experiencing repeated acute stresses originated in studies involving mouse skin in situ along with human keratinocytes and neutrophils in culture, suggesting that this novel mechanism is conserved in human and mouse and may in fact be relevant to chronic inflammatory skin diseases. To test for the latter, we developed and applied a strategy that relies on comparative analysis of transcriptomics data and successfully related key findings obtained in mouse skin under stress to chronic diseases affecting humans, including PSOR, AD, and HS. In addition to test and refine the “TAR12” signature that emanated from our study, future efforts should assess whether enhanced PKCα activity, in particular, and other aspects of TAR indeed play an important role in the genesis and/or maintenance of skin lesions in disorders such as PSOR, HS, AD and others. Additionally, further studies focused on other cell types are needed as repeated stress to the skin engages additional key cellular effectors of inflammation and/or innate immunity such as macrophages, Langerhans cells and dendritic epidermal T-cells (86). Meanwhile the findings we report here deepen our appreciation of the paracrine and cell-autonomous pathways that shape the skin’s response to repeated acute stresses, and may inform research to devise new biomarkers or therapeutic targets for neutrophil-driven inflammatory skin diseases.

## Supporting information

Xu et al. Suppl Materials ALL

## Acknowledgments

The authors are grateful to current and past members of the Coulombe and Parent laboratories for advice and support, to Tenzin Ngodup, Behirda Karaj and Dajia Wang for their contributions to specific phases of the project and to the Platelet Pharmacology and Physiology Core, CyTOF Core, and Rogel Cancer Center Immunology Core at the Univ. of Michigan, and RayBiotech (Peachtree Corner, GA) for their services. EC received fellowship support from NIH grant T32 AR007197 and from the National Psoriasis Foundation. These studies were supported by grants R01 AI152517 (to CAP), R01 AR079418 (to PAC) and P30 AR075043.

## Contributions

YX, CAP and PAC together designed the study and analyzed all findings. YX carried out all experiments involving mice and cultured cells. CJ carried out the analysis of the RNAseq data from mouse skin and the single-cell RNAseq datasets from human skin diseases, and EC led the effort of integrating them in a shared conceptual framework. YX, CAP and PAC led the effort of writing the manuscript, with input from EC and CJ.

## Materials and Methods

### Mouse models and treatments

All mouse experiments involved 2-to 3-month-old male mice in the FVB background and were approved the Institutional Animal Use and Care Committee of University of Michigan Medical School. The *Krt17*^−/−^ (31) and *Krt17^ΔNLS/ΔNLS^* (24) mouse strains were previously described. Mice were treated with 20μl of 0.2mg/ml of TPA (Cell Signaling Technology #4174S), topically on the dorsal side of ear skin, with the other ear treated with acetone vehicle control. For acute UVB treatment, mice were anesthetized with isoflurane and put in a dark cabinet equipped with 4 UVB lamps (305-312nm) (Daavlin Distributing Company). The dorsal side of ear skin was exposed to 400 mJ/cm^2^ of UVB radiation (87). For topical treatment of Gö6976, 100μM Go6976 dissolved in 10μl acetone was applied to mouse ear 30min before TPA treatment.

### Tissue collection and cryosectioning

Mice receiving treatments were euthanized and ear skin tissue was collected at specific time points as indicated. Ear samples were embedded in -40°C optimal cutting temperature compound (O.C.T, Sakura® Finetek USA, #4583). Cryosectioning was performed at -20°C using a CRYOSTAR^TM^ NX50 Cryostat (Thermo Scientific^TM^) and MX35 ultramicrotome blade (Epredia #3053835). 5μm thick cross-sections were cut and placed on positively charged microscope slides (VWR #48311-703), and stored at -40°C until further use.

### Cell lines and treatments

A431 cells were obtained from the American Type Culture Collection (ATCC). A *KRT17* null A431 variant was described (25). Cells were cultured in DMEM medium (Gibco #11995-065) supplemented with 10% FBS and 0.01% Penicillin-Streptomycin (Gibco #15140-122). Cells were treated with 20nM TPA in acetone for the time indicated. For studies using Gö6976, 1μM Gö6976 dissolved in acetone was added 30min before TPA treatment.

### Indirect immunofluorescence and TUNEL staining

For frozen tissue sections, slides stored at – 40℃ were taken to room temperature, dried for 10min, fixed with 4% paraformaldehyde (PFA) (Electron Microscopy Sciences #15710) for 10min at room temperature, followed by 3 washes of 5min with 1x PBS. A circle was drawn around each tissue section with a hydrophobic barrier pen (CALIBIOCHEM #402176). Tissue sections were blocked with blocking buffer (2% normal goat or donkey serum, 1% bovine serum albumin in 1x PBS) for 30min at room temperature.

For cultured cell samples, cells seeded on glass coverslips in 12-well or 24-well plates and cultured overnight at 37°C and 5% CO2 to let cells attached. After treatment, cells were fixed with 4% PFA for 10min, permeabilized with 0.1% Triton X-100 for 5min, then blocked with blocking buffer (5% normal goat or donkey serum in 1x PBS) for 1h at room temperature.

Unconjugated primary antibodies were diluted in blocking buffer and applied overnight at 4°C. On the second day, samples were washed and incubated with fluorophore-conjugated secondary antibodies for 1h at room temperature in dark. Samples were then stained with 1μg/ml of DAPI (Milipore Sigma #268298), washed, mounted with coverslips via FluoroSave reagent (EMD Millipore #345789) and dried overnight. The Click-iT Plus TUNEL Assay for *in situ* apoptosis detection with Alexa Fluor^TM^ 488 (Invitrogen #C10617) was used as recommended by the manufacturer. Both tissue sections and cultured cell samples were imaged using a Zeiss LSM800 confocal microscope (Zeiss, Germany). The antibodies used are listed in **Suppl. Table 1**.

### Transient transfection of GFP-tagged keratin constructs

The GFP-K17WT and GFP-K14WT constructs (pGFP-C3 vector backbone) have been described (24). The K14-K17 chimeras were constructed by PCR-driven overlap extension (88) using the oligonucleotide primers listed in **Suppl. Table 2** with wild-type K17 and wild-type K14 cDNAs as templates. Constructs were transiently transfected into *KRT17* null A431 cells using the SF Cell Line 4D-Nucleofector™ X Kit (Lonza #V4XC-2032) and Lonza 4D-nucleofector X unit “A431 cell” program. 0.5μg of plasmid was used to transfect every 0.4 million cells. Transfected cells were plated on coverslips for immunofluorescence assays or in 6-well plates for conditioned media collection. The cells were allowed to rest for 48 h and then subjected to the desired outcome.

### Proximity ligation assays (PLA)

Cultured cells were fixed with 4% PFA solution for 15min, permeabilized with 0.1% Triton X-100 solution for 10min at room temperature, washed, blocked with 2.5% normal goat serum for 30min at 37°C. Primary antibodies (**Suppl. Table 1**) diluted in blocking buffer were added and the preparations were incubated at 37°C for 1h. The Duolink PLA Fluorescence kit (Millipore Sigma) was used according to the manufacturor’s instructions. Briefly, cells were incubated with PLA probes for 1h at 37°C, with the ligation mixture for 30min at 37°C, and with the amplification mixture for 100min at 37°C. Cells were then stained with DAPI (1μg/ml) for 10min at room temperature and mounted on microscope slides. Images were captured using a Zeiss LSM800 confocal microscope.

### Western blotting and immunoprecipitation

Whole cell lysates were prepared in NP-40 lysis buffer [0.5% NP-40, 150mM NaCl, 20 mM Tris (PH 7.5), EDTA 1mM, 1x cOmplete protease inhibitor cocktail solution (Millipore #11836170001), 1x Pierce™ phosphatase inhibitor cocktail solution (Thermo Scientific #A32957), milliQ water]. Briefly, cultured cells in plates were washed once with cold 1x PBS and transferred to ice. Lysis buffer was added, and cells were scraped off from the plate and rotated at 4℃ for 1h. Supernatants were collected after centrifugation (12,000 x g, 20 min) and total protein levels were measured using Pierce™ BCA Protein Assay Kits (Thermo Scientific #23227). Proteins were denatured by boiling with Laemmli buffer (Bio-Rad #1610747) supplemented with 10% β-mercaptoethanol (β-ME) at 95℃ for 10min. SDS-PAGE electrophoresis was performed on 4-15% gradient gels (Bio-Rad #4561084) or 4-20% gradient gels (GenScript #M00656). Proteins on gels were transferred to nitrocellulose membranes (Bio-Rad #1620115) using a transblot turbo transfer system (Bio-Rad). Blots were blocked in 5% BSA in tris-buffered saline containing 0.1% Tween20 (TBST-T) for 1 h at room temperature and incubated overnight at 4 °C with primary antibodies (cf. **Suppl. Table 1**) diluted in the blocking buffer. Secondary antibodies (cf. **Suppl. Table 1**) diluted in blocking buffer were applied for 1 h at room temperature. Blots were developed using SuperSignal West Pico PLUS chemiluminescent substrate (Thermo Scientific #34580) or ECL^™^ Select Western Blotting Detection Reagent (Cytiva #RPN2235), and imaged using a FluorChem Q system (ProteinSimple). For K17 immunoprecipitation (IP), Dynabeads™ Protein A Immunoprecipitation Kit (Invitrogen #10006D) was used. 50μl of beads was incubated with 2μl of rabbit polyclonal K17 antibody at 4℃ overnight. The same volume of rabbit preimmune serum was used as control. A431 cells (*KRT17* WT and *KRT17* KO) were lysed using NP-40 lysis buffer as described before. Whole cell lysates (250μg of total protein) were incubated with antibody-beads complex at 4℃ overnight. The eluted IP samples were denatured and subjected to western blotting.

### Harvesting conditioned medium (CM)

3 x 10^5^ A431 cells were seeded in each well of 6-well plates and cultured overnight in 2ml of medium supplemented with full serum (10% FBS). On day 2, cells were treated with 20nM TPA in full serum medium for 6h. After rinsing (1x PBS), 2ml serum-reduced medium (0.1% FBS) was added cells incubated for another 24h. Conditioned medium (CM) was collected using a 3ml syringe (BD Syringe) and filtered through a 0.22μm PVDF syringe filter (Millipore #SLGVM33RS), divided into aliquots and stored at – 20°C until use. Measurements of selected cytokines and chemokines in CM were performed by RayBiotech and Immunology Core at University of Michigan.

### Isolation of human peripheral blood neutrophil

Blood from anonymous healthy human donors who had not taken aspirin for 7 days and NSAIDS for 48 hours was obtained by venipuncture from the Platelet Pharmacology and Physiology Core at the University of Michigan. Neutrophils were isolated as described (89). Briefly, whole blood was incubated with an equal volume of 3% dextran (Sigma #D1037) in 0.9% NaCl for 30min at 37°C to facilitate red blood cell sedimentation. The upper layer (plasma) was collected and centrifuged at 400xg for 5min, and the cell pellet containing platelets, monocytes, lymphocytes, and neutrophils was collected and resuspended in 1x mHBSS (150mM NaCl, 4mM KCl, 1.2mM MgCl2, 10mg/ml glucose, and 20mM HEPES). Histopaque-1077 solution (Sigma #10771) was carefully underlaid below the suspension and centrifuged at 400xg for 20min (ACC/DEC = 2/0) to separate peripheral blood mononuclear cells from neutrophils. Residual erythrocytes in the pellet were removed using ACK lysing buffer (Thermo Fisher #A1049201). Purified neutrophils were resuspended in 1x mHBSS and kept at 37℃ until used. In our hands this protocol yields >99% live neutrophils with >95% purity.

### Neutrophil transwell migration assays

were performed in a transwell system consisting of a 24-well tissue culture plate and 3.0-μm pore size PET membrane inserts (Sterlitech Corporation #9323012). Plates and inserts were coated with 2% bovine serum albumin (BSA) for 1h at 37℃ to prevent neutrophil adhesion. Freshly isolated neutrophils (0.4 million cells in 100μl of 1x mHBSS) were added into the upper chamber (insert) and 600μl of CM or control medium (0.1% FBS) into the lower chamber (well of the plate). After a 2-hour incubation at 37℃, the insert together with neutrophils remaining in the upper chamber were discarded. Cells in the bottom chamber were counted using a hemocytometer. The percentage of migrated cells was calculated by relating the cell number in the bottom chamber to total cells seeded in the upper chamber (0.4 million). Data was normalized by subtracting the percentage of migrated cells to control medium (90). When testing the effect of inhibitors, freshly isolated neutrophils were pretreated with 1μM of AZD5069 (CXCR2 inhibitor; Cayman Chemical Company #28297), 1μM of AMG487 (CXCR3 inhibitor; Tocris #4487) for 30min before the transwell migration experiments. A431-derived CM were supplemented with same concentration of inhibitors before used in migration assays. These inhibitors together with 1μg/ml of TNFα neutralizing antibody Infliximab (Novus Biologicals #NBP2-52655) do not alter neutrophil migration in response to fMLF, a potent neutrophil chemoattractant (90), suggesting that neutrophils can respond normally to other chemotactic cues in the presence of the inhibitors (**Suppl. Fig. 3G**). The neutralizing antibody of TNFα was added due to its K17-dependent release (**Suppl. Table 3**) and its suggested role in promoting neutrophil influx induced by antigen challenges *in vivo* (91).

### Quantitative real-time PCR analysis (qRT-PCR)

Neutrophils (10 million cells) were collected prior to treatment (t0) or after 2h of rotation at 37℃ for 2h in mHBSS, reduced serum DMEM (0.1% FBS), CM of acetone treated parental A431 cells, or CM of TPA-treated parental A431 cells. RNA isolation was then performed using Qiagen RNeasy mini kit (Qiagen #74104) following the manufacturer’s protocol. Then, total RNA was converted to complementary DNA (cDNA) using iScript™ cDNA Synthesis Kit (Bio-Rad #1708891). The cDNA obtained was subjected to qRT-PCR using the itaq™ Universal SYBR green kit (Bio-Rad, #1725122) and the CFX 96 Real-Time System (Bio-Rad). The PCR parameters for qRT-PCR screen were 95°C for 5min, followed by 40 cycles of 95°C for 10 sec and 60°C for 30s. No cDNA template controls and a melt curve were included in every PCR run. The normalized expression value of the target gene was determined by first averaging the relative expression of the target gene for each cDNA sample (ΔCq=average Cq_target gene_ – average Cq_reference gene_), and then normalizing the relative expression value of the experimental condition to the control condition (2^-(ΔCqExperimental-ΔCqControl)^). Primers used in qRT-PCR assays are listed in **Suppl. Table 2**.

### Mouse skin immune cell profiling (CyTOF)

The dorsal ears of mice were treated with TPA or acetone. Ear tissues were collected for isolation of immune cells. The tissues were rinsed in 10ml of skin digestion medium (450ml HBSS, 50 mlFBS, 1.1915g HEPES, 0.9356g EDTA, filtered through a 0.22μm PVDF filter) for 30 minutes at 37°C. Then the tissues were transferred to fresh 9 ml of skin digestion medium, cut into small pieces, and 1ml of triple enzyme solution (1mg/ml type I deoxyribonuclease, 1mg/ml type V hyaluronidase, and 5mg/ml collagenase, filtered through a 0.22μm PVDF filter) was added and incubated for 30 minutes at 37°C. The solution was filtered through a 70um filter and centrifuged at 1600 rpm for 5min to form cell pellets. The cell pellet was resuspended with 1 ml of pre-warmed PBS and the cell number was counted. Cells were then centrifuged at 300 x g for 5min and resuspended in 200μl of PBS. Isolated cells were stained following Maxpar Cell Surface Staining with Fresh Fix protocol (Standard BioTools). Briefly, cell viability staining was performed by adding 50μl of Cell-ID Cisplatin (Standard BioTools #201195) to the cell suspension. Cells were washed with Maxpar Cell Staining Buffer (Standard BioTools #201068), centrifuged at 300 × g for 5min, and resuspended in 25μl of staining buffer containing Fc block to incubate for 10min at room temperature. Then 50μL of antibody mixture was added and incubated on ice for 1h. Maxpar Cell Staining Buffer was added for washing and cells were collected after centrifugation. Cells were fixed with 1.6% PFA solution for 10min at room temperature, centrifuged at 800 x g for 5 min. The cell pellet was resuspended in 1ml of 125 nM Cell-ID Intercalator-Ir (Standard BioTools #201192A) made using Maxpar Fix and Perm Buffer (Standard BioTools #201067) and incubated overnight at 4°C. Conventional CyTOF was performed by the UMICH CyTOF Core. Final data were gated and analyzed using FlowJo.

### Bulk RNA sequencing

TPA or acetone was applied to the dorsal ears of *Krt17^-/-^*and WT mice twice, 24h apart (n = 3 mice for each group), and ears were collected at 6h after the second treatment. RNA was isolated and purified using the Qiagen RNeasy mini kit (Qiagen #74104) and Qiagen RNase-Free DNase kit (Qiagen #79254) using the manufacturer’s instructions. RNA samples were transferred to the University of Michigan Advanced Genomics Core (AGC) for RNA quality control, RNA ribo-depletion library preparation, and next-generation sequencing (NovaSeq S4 300 cycle). Core supplied FASTQ files were mapped to the GRCm39.vm30 reference using STAR version 2.7.10a and the ENCODE standard options from the version documentation (92). The reference matching GTF file was used with featureCounts from the Bioconductor R package Rsubread to create the counts matrix. The edgeR Bioconductor package was used to filter low expressing genes using the ‘filterByExpr’ function, calculate log counts per million, and normalization factors using the weighted trimmed mean of M-values (TMM) method. The Limma Biocoductor package, with the precision weights, ‘voom’ approach was used to perform linear models (93). Differential gene expression was determined using custom thresholds as described for each analysis in the main text and in figure legends. ***Note: Deposition of this RNAseq data set in the Gene Expression Omnibus (GEO) repository is pending*.**

### Analysis of human scRNAseq datasets

PSOR, HS, and AD datasets were previously published (4, 55, 64) and analyzed through standard Seurat SCTransform v2 pipeline (94). In short, raw data was filtered to retain cells with [500<unique features<7,500] for PSOR dataset or [500<unique features and <50,000 total counts] for HS and AD dataset as well as a mitochondrial content <15%. Following the SCTransform “v2” normalization and variance stabilization, the top principal components (30 for PSOR, 15 for HS, 20 for AD) were used to construct a shared nearest neighbor (SNN) graph, and subsequently the Louvain algorithm was used to generate clusters (resolution parameter of 0.7 for PSOR, 0.6 for HS and AD). For composite scores, raw counts were library size normalized and log transformed. To identify gene homology (mouse to human), genes upregulated in a K17-dependent manner post-TPA Tx (>8-fold; Adj. P<0.01) were analyzed against the MGI Mouse-Vertebrate homology database. 234 of the 441 genes were identified to have human homologs, totaling 259 unique human genes. Of the 259 human genes homologs, 209 genes showed expression in the PSOR dataset. The genes were then filtered for average expression level across each cell type (as determined by Seurat clustering) in PSOR dataset. Clusters were labeled as the following cell types: Keratinocytes (KCs, containing *KRT10*, *KRT15*, *KRT2*), Melanocytes (Mel, containing *PMEL*), T cells (TC, containing *CD3D*+), myeloid lineage (containing *LYZ*+) and an unknown cluster KRT23-MMP7 (*KRT23*+, *KRT18*+, *MMP7*+). 12 genes were found to have higher expression in keratinocytes (KC) than in myeloid-lineage population. The “TAR12” composite score of cells, representing the average expression level of these 12 genes in each cell, was then calculated. A comparative analysis was conducted between the expression level of TAR12 and level of *KRT6A* and *KRT17* in each cell.

### Image quantification and statistical analyses

Quantitation of immunofluorescence images, PLA images, and western blots were performed using ImageJ. *In vivo* neutrophil infiltration levels were calculated as the area fraction of Ly6g-positive staining in the TPA-treated skin (minimum threshold = 5000) subtracted by background (area fraction of Ly6g-positive staining in acetone-treated skin at the same time point). Quantification of PKCα membrane localization was based on the intensity of the PKCα signal on a line profile (line width 20μm) across adjacent cells.

Fluorescence intensities were calculated as the signal intensity of regions of interest (ROIs) for several fields within tissue sections, and several tissue sections, across biological replicates (e.g., mice). For PLA images, the minimum threshold was set to 3000 to eliminate background noise, and ImageJ Analysis Particles was used to automatically count number of particles >0.3μm^2^ in size. For western blot images, normalized protein levels were calculated by dividing the integrated density of target protein band by the density of the housekeeping protein band. To quantify TUNEL-positive cells, the number of positively stained cells was counted as well as the total number of cells based on DAPI staining. The average percentage of positive cells was then calculated across multiple ROIs for each biological replicate. Error bars on histograms represent standard error of the mean (SEM) across biological replicates. Statistical significance was determined by one-way ANOVA followed by Tukey’s multiple comparisons test, two-way ANOVA followed by Sidak’s multiple comparisons test, and paired or unpaired t test. Differences with p<0.05 were considered statistically significant. All statistical analyses were performed using GraphPad Prism.

## References

1. Schabitz A, Eyerich K, and Garzorz-Stark N. So close, and yet so far away: The dichotomy of the specific immune response and inflammation in psoriasis and atopic dermatitis. J Intern Med. 2021;290(1):27–39.

2. Chiang CC, Cheng WJ, Korinek M, Lin CY, and Hwang TL. Neutrophils in Psoriasis. Front Immunol. 2019;10:2376.

3. Farrar MD, and Ingham E. Acne: inflammation. Clin Dermatol. 2004;22(5):380–4.

4. Gudjonsson JE, Tsoi LC, Ma F, Billi AC, van Straalen KR, Vossen A, et al. Contribution of plasma cells and B cells to hidradenitis suppurativa pathogenesis. JCI Insight. 2020;5(19).

5. Navrazhina K, Garcet S, Zheng X, Hur HB, Frew JW, and Krueger JG. High inflammation in hidradenitis suppurativa extends to perilesional skin and can be subdivided by lipocalin-2 expression. J Allergy Clin Immunol. 2022;149(1):135–44 e12.

6. Burn GL, Foti A, Marsman G, Patel DF, and Zychlinsky A. The Neutrophil. Immunity. 2021;54(7):1377–91.

7. Khou S, Popa A, Luci C, Bihl F, Meghraoui-Kheddar A, Bourdely P, et al. Tumor-Associated Neutrophils Dampen Adaptive Immunity and Promote Cutaneous Squamous Cell Carcinoma Development. Cancers (Basel). 2020;12(7).

8. Zhou X, Chen Y, Cui L, Shi Y, and Guo C. Advances in the pathogenesis of psoriasis: from keratinocyte perspective. Cell Death Dis. 2022;13(1):81.

9. Cohen E, Johnson C, Redmond CJ, Nair RR, and Coulombe PA. Revisiting the significance of keratin expression in complex epithelia. J Cell Sci. 2022;135(20).

10. Cohen E, Johnson, C., Wasikowski, R., Billi, A.C., Tsoi, L.C., Kahlenberg, J.M., Gudjonsson, J.E., Coulombe, P.A. Significance of stress keratin expression in normal and diseased epithelia. Cell Reports. 2023;Submitted.

11. McGowan K, and Coulombe PA. The wound repair-associated keratins 6, 16, and 17. Insights into the role of intermediate filaments in specifying keratinocyte cytoarchitecture. Subcell Biochem. 1998;31:173–204.

12. McGowan KM, and Coulombe PA. Onset of keratin 17 expression coincides with the definition of major epithelial lineages during skin development. J Cell Biol. 1998;143(2):469–86.

13. Proby CM, Churchill, L., Purkis, P. E., Glover, M. T., Sexton, C. J., Leigh, I. M. Keratin 17 expression as a marker for epithelial transformation in viral warts. Am J Pathol. 1993;143(6):1667–78.

14. Ungar B, Garcet S, Gonzalez J, Dhingra N, Correa da Rosa J, Shemer A, et al. An Integrated Model of Atopic Dermatitis Biomarkers Highlights the Systemic Nature of the Disease. J Invest Dermatol. 2017;137(3):603–13.

15. Hughes BR, Morris C, Cunliffe WJ, and Leigh IM. Keratin expression in pilosebaceous epithelia in truncal skin of acne patients. Br J Dermatol. 1996;134(2):247–56.

16. Moll R, Krepler, R., Franke, W.W. Complex cytokeratin polypeptide patterns observed in certain human carcinomas. Differentiation. 1983;23(3):256–69.

17. Baraks G, Tseng R, Pan CH, Kasliwal S, Leiton CV, Shroyer KR, et al. Dissecting the Oncogenic Roles of Keratin 17 in the Hallmarks of Cancer. Cancer Res. 2022;82(7):1159–66.

18. McLean WHI, Rugg, E.L., Lunny, D.P., Morley, S.M., Lane, E.B., Swensson, O., Dopping-Hepenstal, P.J.C., Griffiths, W.A.D., Eady, R.A.J., Higgins, C., Navsaria, H.A., Leigh, I.M., Stachan, T., Kunkeler, L., Munro, C.S. Keratin 16 and keratin 17 mutations cause pachyonychia congenita. Nature Genetics. 1995;9:273–8.

19. Samuelov L, Smith FJD, Hansen CD, and Sprecher E. Revisiting pachyonychia congenita: a case-cohort study of 815 patients. Br J Dermatol. 2020;182(3):738–46.

20. Zieman AG, and Coulombe PA. Pathophysiology of pachyonychia congenita-associated palmoplantar keratoderma: new insights into skin epithelial homeostasis and avenues for treatment. Br J Dermatol. 2020;182(3):564–73.

21. Kim S, Wong P, and Coulombe PA. A keratin cytoskeletal protein regulates protein synthesis and epithelial cell growth. Nature. 2006;441(7091):362-5.

22. Pan X, Kane LA, Van Eyk JE, and Coulombe PA. Type I keratin 17 protein is phosphorylated on serine 44 by p90 ribosomal protein S6 kinase 1 (RSK1) in a growth- and stress-dependent fashion. J Biol Chem. 2011;286(49):42403–13.

23. Chung BM, Arutyunov, A., Ilagan, E., Yao, N., Wills-Karp, M., Coulombe, P.A. Regulation of C-X-C chemokine gene expression by keratin 17 and hnRNP K in skin tumor keratinocytes. J Cell Biol. 2015;208(5):613–27.

24. Jacob JT, Nair RR, Poll BG, Pineda CM, Hobbs RP, Matunis MJ, et al. Keratin 17 regulates nuclear morphology and chromatin organization. J Cell Sci. 2020;133(20).

25. Hobbs RP, DePianto DJ, Jacob JT, Han MC, Chung BM, Batazzi AS, et al. Keratin-dependent regulation of Aire and gene expression in skin tumor keratinocytes. Nat Genet. 2015;47(8):933–8.

26. Luo Y, Zhu Z, Li B, Bai X, Fang H, Qiao P, et al. Keratin 17 Promotes T Cell Response in Allergic Contact Dermatitis by Upregulating C-C Motif Chemokine Ligand 20. Front Immunol. 2022;13:764793.

27. Escobar-Hoyos LF, Shah R, Roa-Pena L, Vanner EA, Najafian N, Banach A, et al. Keratin-17 promotes p27KIP1 nuclear export and degradation and offers potential prognostic utility. Cancer Res. 2015.

28. Nair RR, Hsu J, Jacob JT, Pineda CM, Hobbs RP, and Coulombe PA. A role for keratin 17 during DNA damage response and tumor initiation. Proc Natl Acad Sci U S A. 2021;118(13).

29. Depianto D, Kerns ML, Dlugosz AA, and Coulombe PA. Keratin 17 promotes epithelial proliferation and tumor growth by polarizing the immune response in skin. Nat Genet. 2010;42(10):910–4.

30. Abel EL, Angel JM, Kiguchi K, and DiGiovanni J. Multi-stage chemical carcinogenesis in mouse skin: fundamentals and applications. Nat Protoc. 2009;4(9):1350–62.

31. McGowan KM, Tong X, Colucci-Guyon E, Langa F, Babinet C, and Coulombe PA. Keratin 17 null mice exhibit age- and strain-dependent alopecia. Genes Dev. 2002;16(11):1412–22.

32. Marrakchi S, and Puig L. Pathophysiology of Generalized Pustular Psoriasis. Am J Clin Dermatol. 2022;23(Suppl 1):13–9.

33. Thomas PD, Ebert D, Muruganujan A, Mushayahama T, Albou LP, and Mi H. PANTHER: Making genome-scale phylogenetics accessible to all. Protein Sci. 2022;31(1):8–22.

34. Mi H, Muruganujan A, Huang X, Ebert D, Mills C, Guo X, et al. Protocol Update for large-scale genome and gene function analysis with the PANTHER classification system (v.14.0). Nat Protoc. 2019;14(3):703–21.

35. Kobayashi Y. The role of chemokines in neutrophil biology. Front Biosci. 2008;13:2400–7.

36. Hartl D, Krauss-Etschmann S, Koller B, Hordijk PL, Kuijpers TW, Hoffmann F, et al. Infiltrated neutrophils acquire novel chemokine receptor expression and chemokine responsiveness in chronic inflammatory lung diseases. J Immunol. 2008;181(11):8053–67.

37. Eruslanov EB, Bhojnagarwala PS, Quatromoni JG, Stephen TL, Ranganathan A, Deshpande C, et al. Tumor-associated neutrophils stimulate T cell responses in early-stage human lung cancer. J Clin Invest. 2014;124(12):5466–80.

38. Nicholls DJ, Wiley K, Dainty I, MacIntosh F, Phillips C, Gaw A, et al. Pharmacological characterization of AZD5069, a slowly reversible CXC chemokine receptor 2 antagonist. J Pharmacol Exp Ther. 2015;353(2):340–50.

39. Johnson M, Li AR, Liu J, Fu Z, Zhu L, Miao S, et al. Discovery and optimization of a series of quinazolinone-derived antagonists of CXCR3. Bioorg Med Chem Lett. 2007;17(12):3339–43.

40. Cataisson C, Joseloff E, Murillas R, Wang A, Atwell C, Torgerson S, et al. Activation of cutaneous protein kinase C alpha induces keratinocyte apoptosis and intraepidermal inflammation by independent signaling pathways. J Immunol. 2003;171(5):2703–13.

41. Kroger C, Loschke F, Schwarz N, Windoffer R, Leube RE, and Magin TM. Keratins control intercellular adhesion involving PKC-alpha-mediated desmoplakin phosphorylation. J Cell Biol. 2013;201(5):681–92.

42. Cataisson C, Pearson AJ, Torgerson S, Nedospasov SA, and Yuspa SH. Protein kinase C alpha-mediated chemotaxis of neutrophils requires NF-kappa B activity but is independent of TNF alpha signaling in mouse skin in vivo. J Immunol. 2005;174(3):1686–92.

43. Bornancin F, and Parker PJ. Phosphorylation of threonine 638 critically controls the dephosphorylation and inactivation of protein kinase Calpha. Curr Biol. 1996;6(9):1114–23.

44. Martiny-Baron G, Kazanietz MG, Mischak H, Blumberg PM, Kochs G, Hug H, et al. Selective inhibition of protein kinase C isozymes by the indolocarbazole Go 6976. J Biol Chem. 1993;268(13):9194–7.

45. Lopez-Bergami P, Habelhah H, Bhoumik A, Zhang W, Wang LH, and Ronai Z. RACK1 mediates activation of JNK by protein kinase C [corrected]. Mol Cell. 2005;19(3):309–20.

46. Ringvold HC, and Khalil RA. Protein Kinase C as Regulator of Vascular Smooth Muscle Function and Potential Target in Vascular Disorders. Adv Pharmacol. 2017;78:203–301.

47. Buchau F, Vielmuth F, Waschke J, and Magin TM. Bidirectional regulation of desmosome hyperadhesion by keratin isotypes and desmosomal components. Cell Mol Life Sci. 2022;79(5):223.

48. Adams DR, Ron D, and Kiely PA. RACK1, A multifaceted scaffolding protein: Structure and function. Cell Commun Signal. 2011;9:22.

49. Lei J, Li Q, Gao Y, Zhao L, and Liu Y. Increased PKCalpha activity by Rack1 overexpression is responsible for chemotherapy resistance in T-cell acute lymphoblastic leukemia-derived cell line. Sci Rep. 2016;6:33717.

50. Yang L, Jin L, Ke Y, Fan X, Zhang T, Zhang C, et al. E3 Ligase Trim21 Ubiquitylates and Stabilizes Keratin 17 to Induce STAT3 Activation in Psoriasis. J Invest Dermatol. 2018;138(12):2568–77.

51. Coulombe PA, and Fuchs E. Elucidating the early stages of keratin filament assembly. J Cell Biol. 1990;111(1):153–69

52. Snider NT, and Omary MB. Post-translational modifications of intermediate filament proteins: mechanisms and functions. Nat Rev Mol Cell Biol. 2014;15(3):163–77.

53. Sawant M, Schwarz N, Windoffer R, Magin TM, Krieger J, Mucke N, et al. Threonine 150 Phosphorylation of Keratin 5 Is Linked to Epidermolysis Bullosa Simplex and Regulates Filament Assembly and Cell Viability. J Invest Dermatol. 2018;138(3):627–36.

54. Chung BM, Murray CI, Van Eyk JE, and Coulombe PA. Identification of novel interaction between annexin A2 and keratin 17: evidence for reciprocal regulation. J Biol Chem. 2012;287(10):7573–81.

55. Cheng JB, Sedgewick AJ, Finnegan AI, Harirchian P, Lee J, Kwon S, et al. Transcriptional Programming of Normal and Inflamed Human Epidermis at Single-Cell Resolution. Cell reports. 2018;25(4):871–83.

56. Kerscher B, Willment JA, and Brown GD. The Dectin-2 family of C-type lectin-like receptors: an update. Int Immunol. 2013;25(5):271–7.

57. Burkard P, Schonhart C, Vogtle T, Kohler D, Tang L, Johnson D, et al. A key role for platelet GPVI in neutrophil recruitment, migration and NETosis in the early stages of acute lung injury. Blood. 2023.

58. Huang C, Cao Z, Ma J, Shen Y, Bu Y, Khoshaba R, et al. AKR1B10 activates diacylglycerol (DAG) second messenger in breast cancer cells. Mol Carcinog. 2018;57(10):1300–10.

59. Frank D, Moussi CJ, Ulferts S, Lorenzen L, Schwan C, and Grosse R. Vesicle-Associated Actin Assembly by Formins Promotes TGFbeta-Induced ANGPTL4 Trafficking, Secretion and Cell Invasion. Adv Sci (Weinh). 2023;10(9):e2204896.

60. Yu F, and Floyd-Smith G. Protein kinase C is required for induction of 2’,5’-oligoadenylate synthetases. Exp Cell Res. 1997;234(2):240–8.

61. Wang LD, Hoeltzel M, Butler K, Hare B, Todisco A, Wang M, et al. Activation of the human histamine H2 receptor is linked to cell proliferation and c-fos gene transcription. Am J Physiol. 1997;273(6):C2037–45.

62. Gkika D, Topala CN, Chang Q, Picard N, Thebault S, Houillier P, et al. Tissue kallikrein stimulates Ca(2+) reabsorption via PKC-dependent plasma membrane accumulation of TRPV5. EMBO J. 2006;25(20):4707–16.

63. Pula B, Witkiewicz W, Dziegiel P, and Podhorska-Okolow M. Significance of podoplanin expression in cancer-associated fibroblasts: a comprehensive review. Int J Oncol. 2013;42(6):1849–57.

64. Ma F, Gharaee-Kermani M, Tsoi LC, Plazyo O, Chaskar P, Harms P, et al. Single-cell profiling of prurigo nodularis demonstrates immune-stromal crosstalk driving profibrotic responses and reversal with nemolizumab. J Allergy Clin Immunol. 2023.

65. Yardman-Frank JM, and Fisher DE. Skin pigmentation and its control: From ultraviolet radiation to stem cells. Exp Dermatol. 2021;30(4):560–71.

66. Natoli G, and Ostuni R. Adaptation and memory in immune responses. Nat Immunol. 2019;20(7):783–92.

67. Naik S, Larsen SB, Gomez NC, Alaverdyan K, Sendoel A, Yuan S, et al. Inflammatory memory sensitizes skin epithelial stem cells to tissue damage. Nature. 2017;550(7677):475-80.

68. Larsen SB, Cowley CJ, Sajjath SM, Barrows D, Yang Y, Carroll TS, et al. Establishment, maintenance, and recall of inflammatory memory. Cell Stem Cell. 2021;28(10):1758–74 e8.

69. Naik S, and Fuchs E. Inflammatory memory and tissue adaptation in sickness and in health. Nature. 2022;607(7918):249-55.

70. Ma S, Rao L, Freedberg IM, and Blumenberg M. Transcriptional control of K5, K6, K14, and K17 keratin genes by AP-1 and NF-kappaB family members. Gene Expr. 1997;6(6):361–70.

71. Shi X, Jin L, Dang E, Chang T, Feng Z, Liu Y, et al. IL-17A upregulates keratin 17 expression in keratinocytes through STAT1- and STAT3-dependent mechanisms. J Invest Dermatol. 2011;131(12):2401–8.

72. Kawalerski RR, Gonçalves MT, Pan C-H, Tseng R, Roa-Peña L, Leiton CV, et al. Disassembly of embryonic keratin filaments promotes pancreatic cancer metastases. bioRxiv. 2022:2022.08.27.504988.

73. Hendriks IA, and Vertegaal AC. A comprehensive compilation of SUMO proteomics. Nat Rev Mol Cell Biol. 2016;17(9):581–95.

74. Partovian C, Zhuang Z, Moodie K, Lin M, Ouchi N, Sessa WC, et al. PKCalpha activates eNOS and increases arterial blood flow in vivo. Circ Res. 2005;97(5):482–7.

75. Kim H, Zamel R, Bai XH, and Liu M. PKC activation induces inflammatory response and cell death in human bronchial epithelial cells. PLoS One. 2013;8(5):e64182.

76. Wei XF, Zhou QG, Hou FF, Liu BY, and Liang M. Advanced oxidation protein products induce mesangial cell perturbation through PKC-dependent activation of NADPH oxidase. Am J Physiol Renal Physiol. 2009;296(2):F427–37.

77. Church DJ, Braconi S, Vallotton MB, and Lang U. Protein kinase C-mediated phospholipase A2 activation, platelet-activating factor generation and prostacyclin release in spontaneously beating rat cardiomyocytes. Biochem J. 1993;290 (Pt 2)(Pt 2):477–82.

78. Fisher GJ, Talwar HS, Baldassare JJ, Henderson PA, and Voorhees JJ. Increased phospholipase C-catalyzed hydrolysis of phosphatidylinositol-4,5-bisphosphate and 1,2-sn-diacylglycerol content in psoriatic involved compared to uninvolved and normal epidermis. J Invest Dermatol. 1990;95(4):428–35.

79. Cataisson C, Pearson AJ, Tsien MZ, Mascia F, Gao JL, Pastore S, et al. CXCR2 ligands and G-CSF mediate PKCalpha-induced intraepidermal inflammation. J Clin Invest. 2006;116(10):2757–66.

80. Rotty JD, and Coulombe PA. A wound-induced keratin inhibits Src activity during keratinocyte migration and tissue repair. J Cell Biol. 2012;197(3):381–9.

81. Wang F, Chen S, Liu HB, Parent CA, and Coulombe PA. Keratin 6 regulates collective keratinocyte migration by altering cell-cell and cell-matrix adhesion. J Cell Biol. 2018;217(12):4314–30.

82. Ku NO, Toivola DM, Strnad P, and Omary MB. Cytoskeletal keratin glycosylation protects epithelial tissue from injury. Nat Cell Biol. 2010;12(9):876–85.

83. Paramio JM, Segrelles C, Ruiz S, and Jorcano JL. Inhibition of protein kinase B (PKB) and PKCzeta mediates keratin K10-induced cell cycle arrest. Mol Cell Biol. 2001;21(21):7449–59.

84. Sahlgren CM, Pallari HM, He T, Chou YH, Goldman RD, and Eriksson JE. A nestin scaffold links Cdk5/p35 signaling to oxidant-induced cell death. EMBO J. 2006;25(20):4808–19.

85. Wang Y, Liao G, Wu Y, Wang R, and Tang DD. The intermediate filament protein nestin serves as a molecular hub for smooth muscle cytoskeletal signaling. Respir Res. 2023;24(1):157.

86. Nestle FO, Di Meglio P, Qin JZ, and Nickoloff BJ. Skin immune sentinels in health and disease. Nat Rev Immunol. 2009;9(10):679–91.

87. Cela EM, Friedrich A, Paz ML, Vanzulli SI, Leoni J, and Gonzalez Maglio DH. Time-course study of different innate immune mediators produced by UV-irradiated skin: comparative effects of short and daily versus a single harmful UV exposure. Immunology. 2015;145(1):82–93.

88. Heckman KL, and Pease LR. Gene splicing and mutagenesis by PCR-driven overlap extension. Nat Protoc. 2007;2(4):924–32.

89. Kremserova S, and Nauseef WM. Isolation of Human Neutrophils from Venous Blood. Methods Mol Biol. 2020;2087:33–42.

90. SenGupta S, Hein LE, Xu Y, Zhang J, Konwerski JR, Li Y, et al. Triple-Negative Breast Cancer Cells Recruit Neutrophils by Secreting TGF-beta and CXCR2 Ligands. Front Immunol. 2021;12:659996.

91. Vieira SM, Lemos HP, Grespan R, Napimoga MH, Dal-Secco D, Freitas A, et al. A crucial role for TNF-alpha in mediating neutrophil influx induced by endogenously generated or exogenous chemokines, KC/CXCL1 and LIX/CXCL5. Br J Pharmacol. 2009;158(3):779–89.

92. Dobin A, Davis CA, Schlesinger F, Drenkow J, Zaleski C, Jha S, et al. STAR: ultrafast universal RNA-seq aligner. Bioinformatics. 2013;29(1):15–21.

93. Law CW, Chen Y, Shi W, and Smyth GK. voom: Precision weights unlock linear model analysis tools for RNA-seq read counts. Genome Biol. 2014;15(2):R29.

94. Choudhary S, and Satija R. Comparison and evaluation of statistical error models for scRNA-seq. Genome Biol. 2022;23(1):27.

