## Supplementary material for "Keratin 17- and PKCα-dependent transient amplification of neutrophil influx after repeated stress to the skin": Xu et al. Suppl Materials ALL

### List of contents

#### Supplemental Figures 1-6:

Supplemental Figure 1. TAR is not related to the skin barrier status and is driven by local signal(s) *in situ.*

Supplemental Figure 2. Additional analyses of the K17-dependent skin response to TPA treatments *in vivo.*

Supplemental Figure 3. *Ex vivo* analyses of K17-dependent neutrophil recruitment.

Supplemental Figure 4. TAR triggered by TPA-UVB dual treatment in mouse skin is partially PKCα-dependent.

Supplemental Figure 5, Effect of TPA treatment on desmosome and protein solubility in A431 keratinocytes.

Supplemental Figure 6. A TAR transcriptional signature occurs in stressed keratinocytes from human inflammatory skin diseases (complement to Figure 8).

#### Supplemental Tables 1-7:

Supplemental Table 1. List of antibodies used in this study

Supplemental Table 2. Oligonucleotide primer sequences used when making K17-K14 chimeric cDNAs and conducting qRT-PCR assays.

Supplemental Table 3. ELISA data for chemokines and cytokines analyzed in conditioned medium (CM) from A431 keratinocytes.

Supplemental Table 4. Genes significantly upregulated at 6h after dual TPA TX in WT and *Krt17*$^{-/-}$ mouse skin.

Supplemental Table 5. Genes significantly downregulated at 6h after dual TPA TX in WT and *Krt17*$^{-/-}$ mouse skin.

Supplemental Table 6. Identification of 254 human gene homologs corresponding to the K17-dependent TAR genes identified in mice.

Supplemental Table 7. Identification of the TAR12 transcriptomic signature.

**Supplemental Figure 1. Xu et al.**

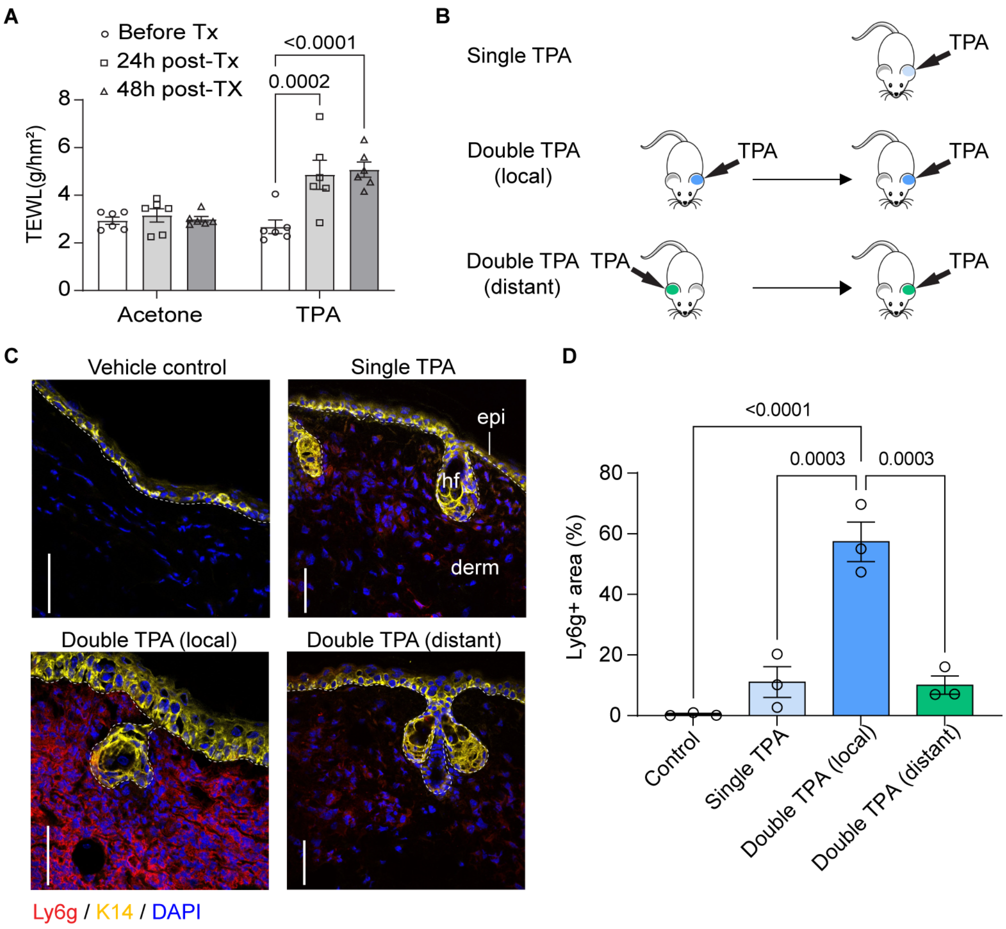

**Supplemental Figure 1** (Xu *et al.*).

**TAR is not related to the skin barrier status and is driven by local signal(s) *in situ.***

**A)** Trans-epidermal water loss measurements (TEWL) of mouse ears treated with a single dose of acetone or TPA. n=6. Two-way ANOVA. **B)** Strategy to test whether TAR is mediated by local or systemic signals. Dual TPA treatment was applied to either the same ear ("local") or contra-lateral ears ("distant"), 24h apart. **C)** Tissue sections from B) were immunostained for Ly6g, K14, and nuclei (DAPI). Scale bars: 50μm. epi, epidermis; derm, dermis; hf, hair follicle. Dashed lines depict the dermo-epidermal interface. **D)** Quantitation of neutrophil fluorescence signal (surface area measurements) of data in C, n=3 mice. Data reported as mean ± SEM. One-way ANOVA.

**Supplemental Figure 2. Xu et al.**

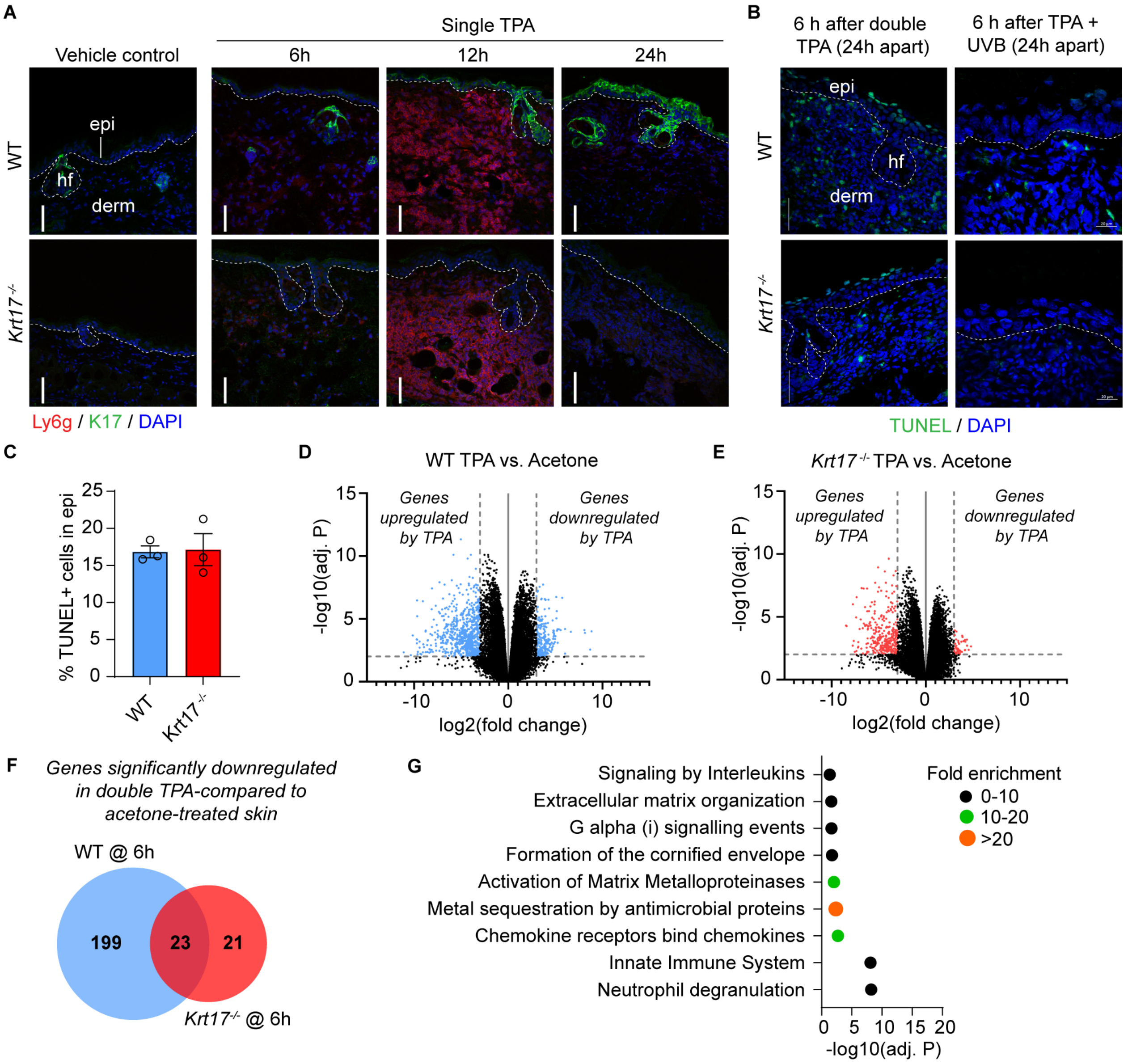

**<u>Supplemental Figure 2</u> (Xu *et al.*).**

**Additional analyses of the K17-dependent skin response to TPA treatments *in vivo.***

**A)** Sections from WT and *Krt17*$^{-/-}$ mouse skin treated with acetone or single TPA were immuno-stained for Ly6g, K14, and nuclei (DAPI). Scale bars: 50μm. epi, epidermis; derm, dermis; hf, hair follicle. Dashed lines depict the dermo-epidermal interface. **B)** TUNEL staining of WT and *Krt17*$^{-/-}$ mouse ear skin at 6h after either double TPA Tx (24h apart) or TPA-UVB combination Tx (24h apart). Scale bar: 50μm**. C)** Percentage of TUNEL positive cells in epidermis after double TPA (data from B). n=3 mice. Data are shown as mean ± SEM. Unpaired t-test. **D-E)** Mouse ear tissues were harvested at 6h after double acetone or dual TPA Tx, 24h apart, and processed for bulk RNAseq analysis. Volcano plots reporting on changes in mRNA transcripts levels in TPA-Tx vs. vehicle-Tx for D) WT skin and E) *Krt17*$^{-/-}$ skin. Genes that are significantly up- or down-regulated by 8-fold or more (adj. P < 0.01) are highlighted in blue (WT) or red (*Krt17*$^{-/-}$).

**F)** Comparison of genes significantly downregulated after dual TPA-Tx vs. to dual acetone-Tx in WT and *Krt17*$^{-/-}$ mouse skin (Cutoffs: FDR-adjusted P < 0.01, fold change > 8). **G)** Panther over-representation test using Reactome pathways (FDR-adjusted P < 0.05) for the 268 genes showing significant upregulated expression in both WT and *Krt17*$^{-/-}$ skin after double TPA Tx (see Figure 2I).

**Supplemental Figure 3. Xu et al.**

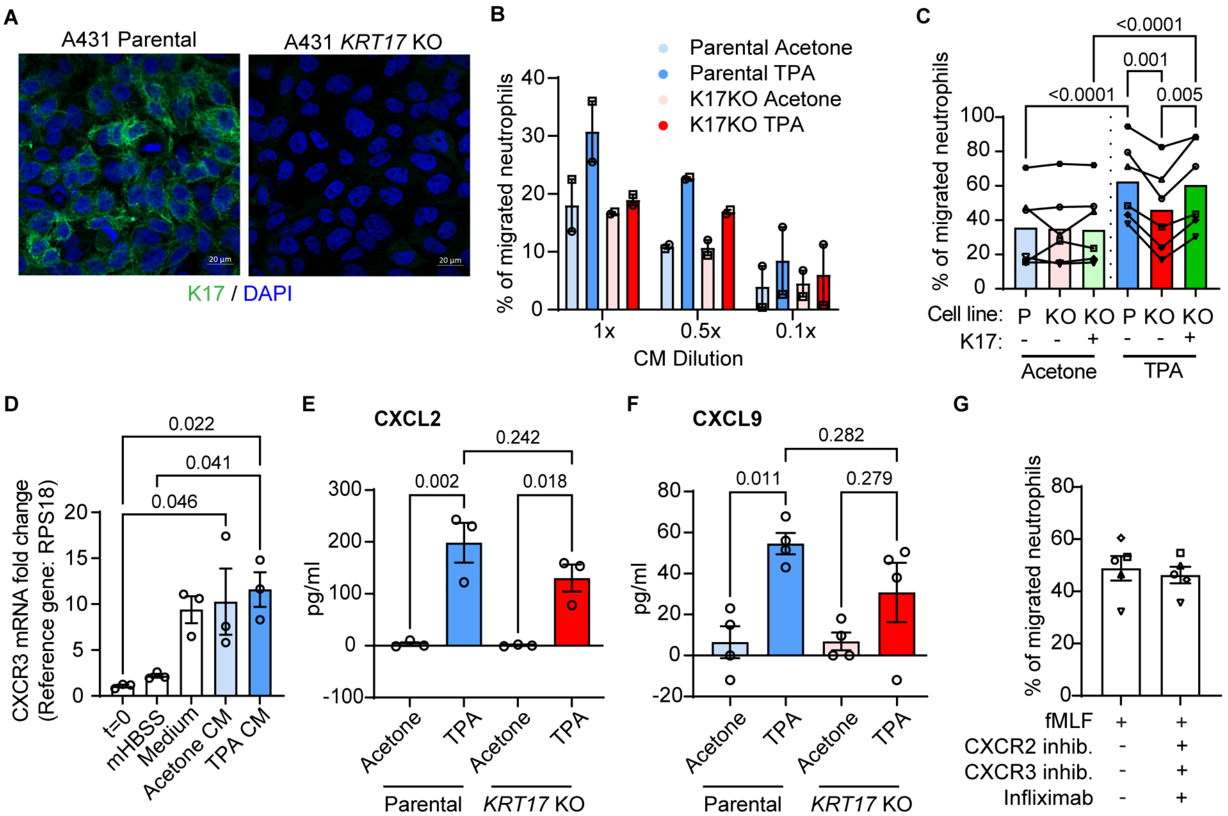

**<u>Supplemental Figure 3</u> (Xu *et al.*).**

***Ex vivo* analyses of K17-dependent neutrophil recruitment.**

**A)** Parental and *KRT17* null A431 keratinocytes stained for K17 and nuclei (DAPI). Scale bars: 20µm. **B)** Migration of human primary neutrophils towards A431 conditioned medium (CM) "titrated" (i.e., diluted) as indicated using control medium (n=2). **C)** Migration of human primary neutrophils towards CM from GFP-transfected parental A431, GFP-transfected *KRT17* null A431, and GFP-K17-transfected *KRT17* null A431 cells. Individual symbols depict data using neutrophils from different donors (n=6). Data reported as mean ± SEM. One-way ANOVA. **D)** Transcript levels of CXCR3 in human primary neutrophils before (t=0) and after treatment with buffer (mHBSS, 2h), culture medium control (2h), CM from acetone-Tx parental A431 (2h), or CM from TPA-Tx parental A431 (2h). n=3. Data reported as mean ± SEM. One-way ANOVA. **E-F)** ELISA measurements for selected chemokine and cytokine levels in A431 CM (reported as pg/ml). n=3 for CXCL2 (2 technical replicates each), and n=4 for CXCL9 (4 technical replicates for 2 measurements, 2 technical replicates for the other 2 measurements). Data reported as mean ± SEM. One-way ANOVA. **G)** Migration of human primary neutrophils towards DMSO (vehicle control) or fMLF, with or without the addition of a CXCR2 antagonist, a CXCR3 antagonist, and infliximab (anti-TNFa). n=5. Data reported as mean ± SEM. Paired t-test.

**Supplemental Figure 4. Xu et al.**

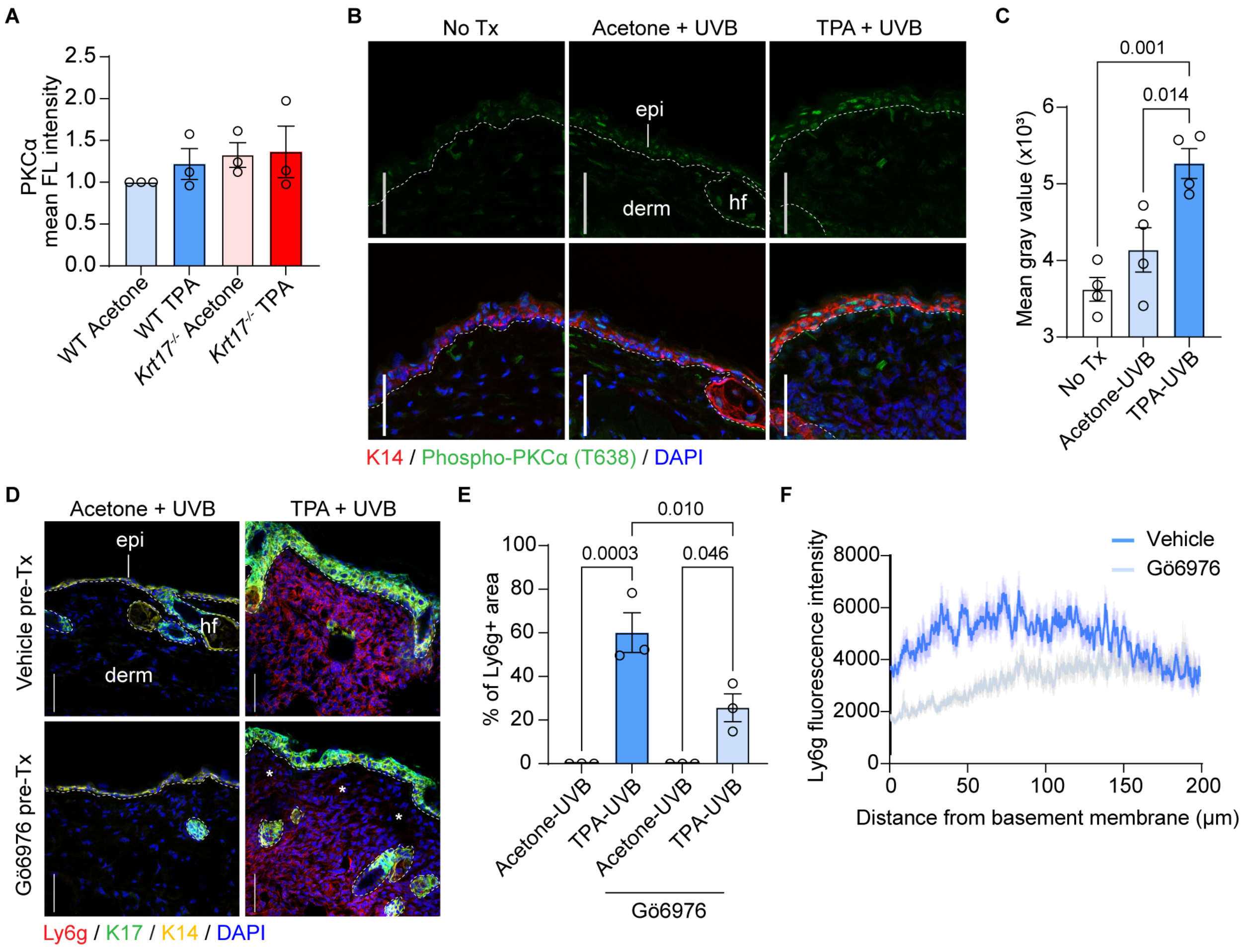

**<u>Supplemental Figure 4</u> (Xu *et al.*).**

**TAR triggered by TPA-UVB dual treatment in mouse skin is partially PKCα-dependent.**

**A)** Quantitation of total PKCα levels in epidermis of WT and *Krt17*$^{-/-}$ mouse skin at 6h after dual TPA Tx, 24h apart. n=3 mice. Data reported as mean ± SEM. One-way ANOVA. **B)** WT and *Krt17*$^{-/-}$ mouse ears were treated with TPA followed by UVB 24h later. Tissues were collected at 6h after UVB and immunostained for phosphor-PKCα (T638), K14, and nuclei (DAPI). Scale bars: 50µm. epi, epidermis; derm, dermis; hf, hair follicle. Dashed lines depict the dermo-epidermal interface. **C)** Quantitation of the phospho-PKCα signal in data from B. n=4 mice. Data are shown as mean ± SEM. One-way ANOVA. **D)** Impact of pre-treatment with the PKCα inhibitor Gö6976 at 30min prior to the 2$^{nd}$ Tx, UVB (24h interval), in WT mouse ear skin. Sections were immunostained for Ly6g, K17, K14, and nuclei (DAPI). Scale bars: 50µm. Asterisks denote the loss of epidermis-proximal pool of neutrophils when pretreated with Gö6976. **E-F)** Quantitation of neutrophil fluorescence signals using E) surface area measurements and by **F)** line vectors extending from the basement membrane to the bottom of the dermis. n=3 mice. Data reported as mean ± SEM. One-way ANOVA.

**Supplemental Figure 5. Xu et al.**

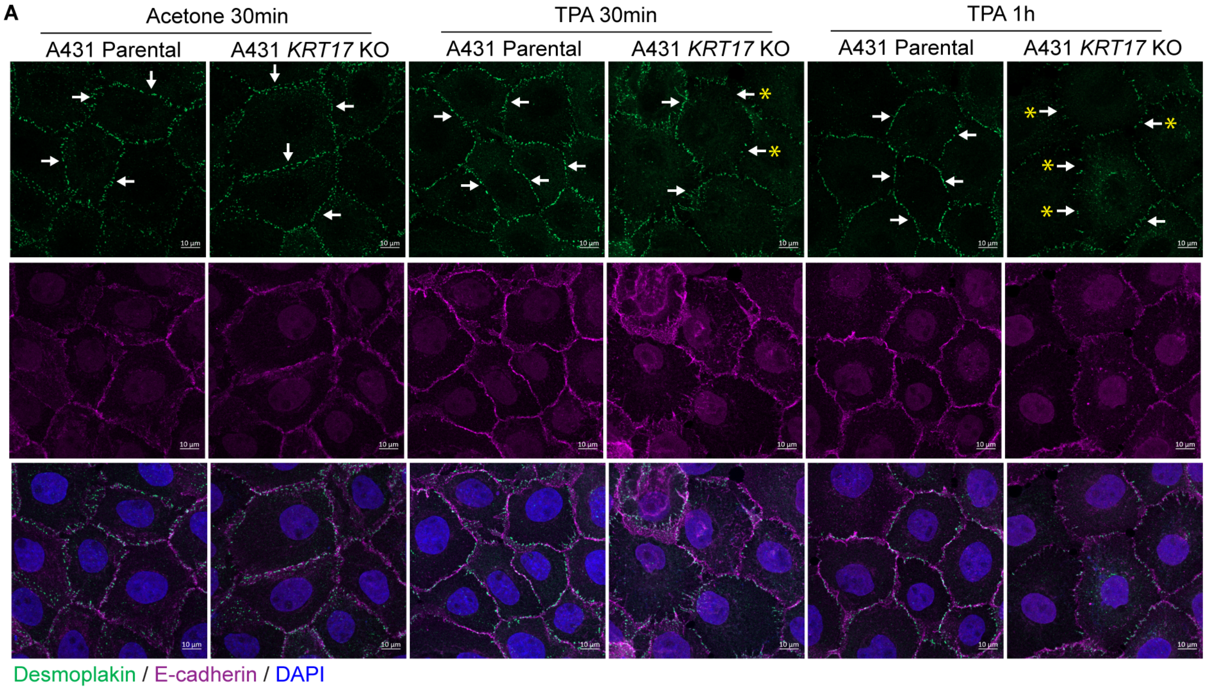


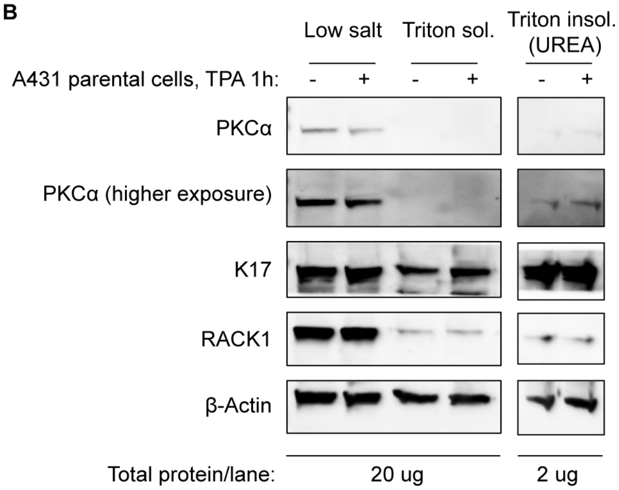

**Supplemental Figure 5** (Xu *et al.*).

**Effect of TPA treatment on desmosome and protein solubility in A431 keratinocytes.**

**A)** Parental and *KRT17* null A431 cells were stained for desmoplakin, e-cadherin, and nuclei (DAPI) at 30 min after acetone-control TX, or at 30 and 60 min after TPA Tx. Scale bars: 10μm. Arrows point to desmoplakin staining at cell-cell borders, and arrows with asterisks denote partial loss of desmoplakin staining at cell-cell borders after TPA Tx. **B)** Western blot analysis of the solubility of K17, PKCα, RACK1 and (control) β-Actin in parental A431 cells at 1h after TPA Tx. 20μg of total protein was loaded in each lane for the low-salt and triton-soluble fractions, and 2μg of total protein was loaded in each lane for the triton-insoluble fractions.

### Supplemental Figure 6. Xu et al.

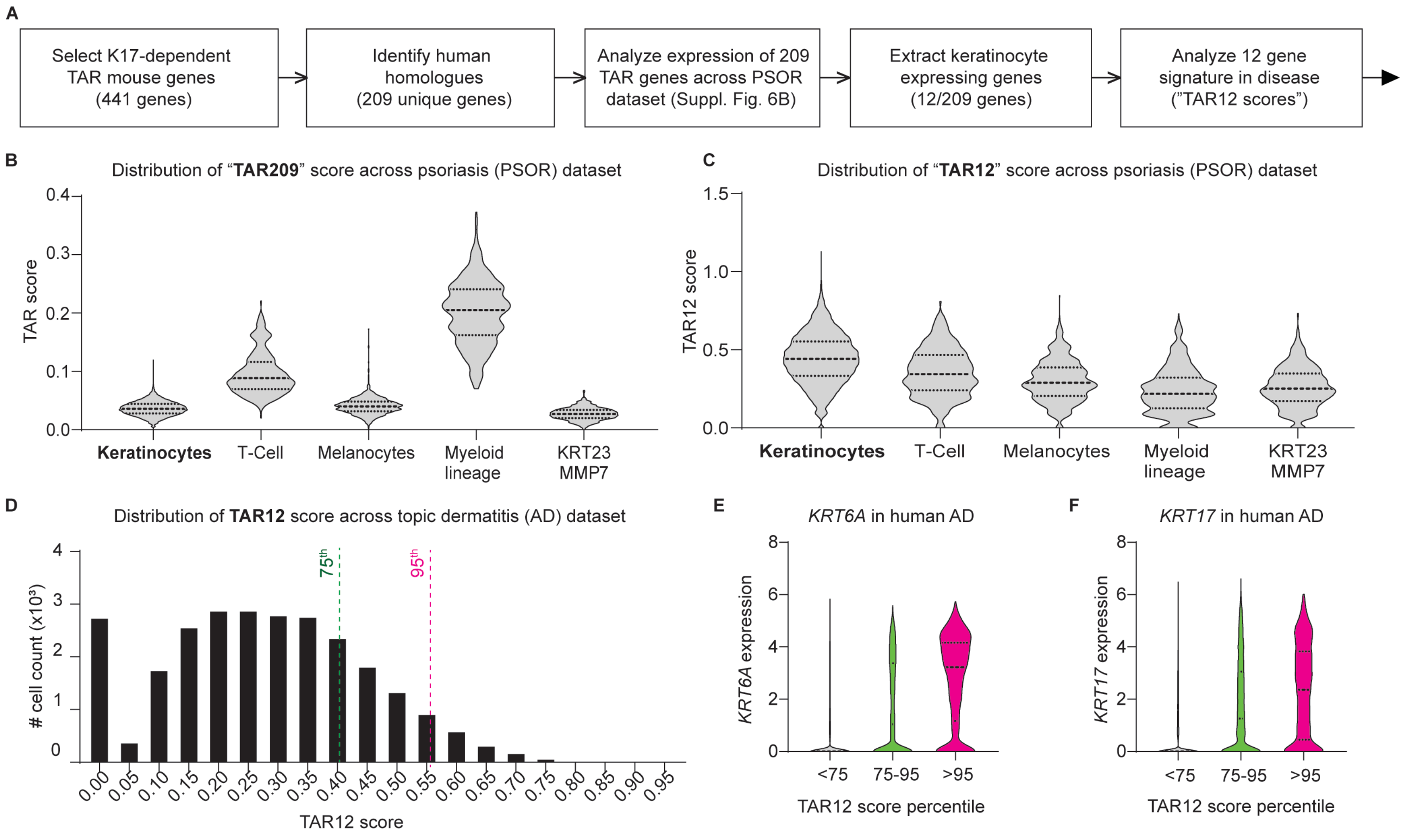

**<u>Supplemental Figure 6</u> (Xu *et al.*).**

**A TAR transcriptional signature occurs in stressed keratinocytes from human inflammatory skin diseases (complement to Figure 8).**

**A)** Strategy for i) devising a TAR-related signature from analyses of RNAseq data in dual TPA-Tx mouse skin and ii) harmonize it with the human transcriptome using an existing dataset from individuals with psoriasis (PSOR) (55) (see Methods). The set of 441 genes significantly upregulated (>8-fold; adj P<0.01) in WT, but not in *Krt17*$^{-/-}$, mouse skin at 6h after dual TPA Tx (see Fig. 2I) were designated as K17-dependent TAR genes. 209 homologs were identified in the PSOR dataset (55), forming the basis for a "TAR209" score. Twelve of these 209 genes are expressed at significant levels in PSOR keratinocytes, as opposed to other cell types (see B and C), forming the basis for the TAR12 composite score. **B)** Expression of the "TAR209" score in specific cell clusters within the PSOR dataset (55). **C)** Expression of the "TAR12" score, which based on the 12 genes showing higher expression level in keratinocytes within the PSOR dataset (*ANGPTL4, IFITM1, IFI16, PHLDA2, KLK1, PDPN, AKR1B10, ACAT2, TUBB6, OAS1, C19orf48, HRH2*), in different cell clusters in this dataset. **D)** Distribution of TAR12 scores in all individual cells from the AD single cell dataset (64). **E, F)** Violin plots reporting on the expression levels of (E) *KRT6A* and (F) *KRT17* in cells showing high, medium, and low TAR12 scores. Cut-offs were arbitrarily set at the 95th percentile for a high TAR score and at the 75th percentile for a medium TAR score based on the distributions shown in D.

**<u>Description of Supplemental Tables 1-7</u> (collated)**

**<u>Supplemental Table 1</u> (Xu *et al.*).**

**List of antibodies used in this study.**

Antibodies utilized in immunofluorescence, proximity ligation assay (PLA), and western blotting.

**<u>Supplemental Table 2</u> (Xu *et al.*).**

**Oligonucleotide primer sequences used when making K17-K14 chimeric cDNAs and conducting qRT-PCR assays.**

Legend: *K14h-K17r-K17t: Chimeric construct consisting of the head domain of K14 and the rod and tail domains of K17; K17h-K14r-K14t: Chimeric construct consisting of the head domain of K17 and the rod and tail domains of K14.

**<u>Supplemental Table 3</u> (Xu *et al.*).**

**ELISA data for chemokines and cytokines analyzed in conditioned medium (CM) from A431 keratinocytes.**

Corresponding sample numbers indicating CM collected from the same experimental batch.

Legend: *Blue: Values below limit of detection; *Red: Values above the highest standards.

**<u>Supplemental Table 4</u> (Xu *et al.*).**

**Genes significantly upregulated at 6h after dual TPA TX in WT and *Krt17*$^{-/-}$ mouse skin.**

Complete list of significantly upregulated genes at 6h after dual TPA treatment of ear skin, 24h apart, in WT and *Krt17*$^{-/-}$ mice. (Cutoff: Adjusted P<0.01, Fold change>8).

**<u>Supplemental Table 5</u> (Xu *et al.*).**

**Genes significantly downregulated at 6h after dual TPA TX in WT and *Krt17*$^{-/-}$ mouse skin.**

Complete list of significantly downregulated genes at 6h after dual TPA treatment of ear skin, 24h apart, in WT and *Krt17*$^{-/-}$ mice. (Cutoff: Adjusted P<0.01, Fold change>8).

**<u>Supplemental Table 6</u> (Xu *et al.*).**

**Identification of 254 human gene homologs corresponding to the K17-dependent TAR genes identified in mice.**

The 441 genes that are upregulated >8-fold (adj. P<0.01) in the bulk RNAseq dataset in WT, but not in *Krt17*$^{-/-}$, mouse skin at 6h after dual TPA Tx (24h apart) were set aside as K17-dependent

TAR genes. Their corresponding human gene homologs (254 genes) were identified and are listed in this table.

**<u>Supplemental Table 7</u> (Xu *et al.*).**

**Identification of the TAR12 transcriptomic signature.**

A total of 209 genes from the 254 TAR-related human homologs (see **Suppl. Table 6**) were found to occur in the PSOR single cell RNAseq dataset (see Cheng et al., Cell Reports 2018; ref. 55). 12 of these genes show preferential expression in keratinocytes (KC) over the myeloid-lineage cells (see **Suppl. Fig. 6C**). These 12 genes form the basis for the “TAR12” composite score.

**Supplemental Table 1. List of antibodies used in this study.**

| Antigen | Host species | Company and catalog number | Dilution |
|---|---|---|---|
| **Immunofluorescence** | | | |
| K14 | chicken | BioLegend #906004 | 1:1000 |
| K17 | rabbit | | 1:1000 |
| Ly6g (Phycoerythrin conjugated) | rat | BioLegend #127607 | 1:100 |
| PKCα | mouse | Santa cruz #8393 | 1:200 |
| Phospho-PKCα (T638) | rabbit | Novus #NBP2-66799 | 1:500 |
| Phospho-PKCα (S657) | rabbit | Abcam #180848 | 1:1000 |
| E-cadherin | rabbit | Cell Signaling Technology, #3195S | 1:1000 |
| Desmoplakin 1& 2 | mouse | Millipore #CBL173 | 1:1000 |
| chicken IgY (Alexa 647 conjugated) | goat | Abcam #150171 | 1:1000 |
| rabbit IgG (Alexa 647 conjugated) | goat | Jackson ImmunoResearch #111-607-003 | 1:400 |
| ms IgG (Alexa 647 conjugated) | goat | Jackson ImmunoResearch #115-605-146 | 1:400 |
| rabbit IgG (Alexa 594 conjugated) | goat | Thermo Fisher Scientific #A11037 | 1:1000 |
| ms IgG (Alexa 594 conjugated) | goat | Fisher Scientific #A11032 | 1:1000 |
| rabbit IgG (Alexa 488 conjugated) | goat | Abcam #150077 | 1:1000 |
| mouse IgG (Alexa 488 conjugated) | goat | Jackson ImmunoResearch #115-545-003 | 1:400 |
| **Proximity ligation assays (PLA)** | | | |
| K17 | rabbit | | 1:1000 |
| Phosho-PKCα (S657) | rabbit | Abcam #180848 | 1:1000 |
| PKCα | mouse | Santa Cruz #8393 | 1:200 |
| RACK1 | mouse | Santa Cruz #17754 | 1:200 |
| **Western blotting** | | | |
| K17 | rabbit | | 1:1000 |
| Histone H3 | rabbit | Cell Signaling Technology #9715 | 1:1000 |
| β-Actin | rabbit | Cell Signaling Technology #4967 | 1:1000 |
| Phosho-PKCα (T638) | rabbit | Novus #NBP2-66799 | 1:1000 |
| PKCα | mouse | Santa cruz #8393 | 1:500 |
| K17 | mouse | Santa cruz #393002 | 1:1000 |
| RACK1 | mouse | Santa cruz #17754 | 1:1000 |
| Phospho-JNK (Thr183/Tyr185) | mouse | Cell signaling Technology #9255 | 1:1000 |
| rabbit IgG (HRP linked) | goat | Cell Signaling Technology #7074P2 | 1:3000 |
| mouse IgG (HRP linked) | goat | Cell Signaling Technology #7076P2 | 1:3000 |

**Supplemental Table 2. Oligonucleotide primer sequences used when making K17-K14 chimeric cDNAs and conducting qRT-PCR assays.**

| Primer Name | Species | FWD/ REV | Sequence (5'-->3') | Restriction site | Notes |
|---|---|---|---|---|---|
| **K14h-K17r-K17t* chimeragenesis** | | | | | |
| 5-XhoI_K14 | Mouse | forward | aaactcgagATGGCCACCTGC | XhoI | Primer A for chimeragenesis and insertion into vector backbone |
| 5-17rodtail_3-14head | Mouse | reverse | GTTCTGCATGGTGGCCTTCTCACTGCCCACCAGGAG | None | Primer B for chimeragenesis |
| 5-14head_3-17rodtail | Mouse | forward | CTCCTGGTGGGCAGTGAGAAGGCCACCATGCAGAAC | None | Primer C for chimeragenesis |
| K17_3-HindIII | Mouse | reverse | aaaaagcttTTAACGGGTGGT | HindIII | Primer D for chimeragenesis and insertion into vector backbone |
| **K17h-K14r-K14t* chimeragenesis** | | | | | |
| 5-XhoI_K17 | Mouse | forward | aaactcgagATGACCACCACC | XhoI | Primer A for chimeragenesis and insertion into vector backbone |
| 5-14rodtail_3-17head | Mouse | reverse | GTTCTGCATGGTCACTTTCTCCCCTCCAGCCAGCAG | None | Primer B for chimeragenesis |
| 5-17head_3-14rodtail | Mouse | forward | CTGCTGGCTGGAGGGGAGAAAGTGACCATGCAGAAC | None | Primer C for chimeragenesis |
| K14_3-HindIII | Mouse | reverse | aaaaagcttTTAGTTCTTGGT | HindIII | Primer D for chimeragenesis and insertion into vector backbone |
| **qRT-PCR** | | | | | |
| CXCR3 FWD | Human | forward | CCACCTAGCTGTAGCAGACAC | | |
| CXCR3 REV | Human | reverse | AGGGCTCCTGCGTAGAAGTT | | |
| RPS18 FWD | Human | forward | GCGGCGGAAAATAGCCTTTG | | |
| RPS18 REV | Human | reverse | GATCACACGTTCCACCTCATC | | |

**Supplemental Table 3. ELISA data for chemokines and cytokines analyzed in conditioned medium (CM) from A431 keratinocytes.**

| Gene | Analyte (pg/ml) | Sample # | Media only | Parental, Acet | Parental, TPA | KRT17 KO, Acet | KRT17 KO, TPA |
|---|---|---|---|---|---|---|---|
| **Multiplexing ELISA conducted by RayBiotech, Inc (Measurements performed in quadruplicates)*** | | | | | | | |
| GRO | CXCL1/2/3 | 1 | 0 | 70251 | 86128 | 72196 | 78677 |
| | | 2 | 778 | 88014 | 97103 | 85034 | 86403 |
| ENA-78 | CXCL5 | 1 | 1 | 0 | 7 | 0 | 2 |
| | | 2 | 0 | 0 | 4 | 0 | 0 |
| GCP-2 | CXCL6 | 1 | 0 | 1 | 1 | 0 | 1 |
| | | 2 | 0 | 0 | 0 | 0 | 0 |
| IL-8 | CXCL8 | 1 | 16 | 121 | 1491 | 163 | 1392 |
| | | 2 | 0 | 103 | 1264 | 210 | 1523 |
| MIG | CXCL9 | 1 | 0 | 15 | 52 | 10 | 47 |
| | | 2 | 0 | 0 | 68 | 0 | 51 |
| I-TAC | CXCL11 | 1 | 9 | 18 | 74 | 4 | 37 |
| | | 2 | 9 | 0 | 63 | 0 | 38 |
| SDF-1a | CXCL12 | 1 | 0 | 0 | 37 | 0 | 6 |
| | | 2 | 0 | 3 | 0 | 19 | 0 |
| MCP-1 | CCL2 | 1 | 4 | 7 | 14 | 3 | 10 |
| | | 2 | 0 | 0 | 1 | 2 | 0 |
| RANTES | CCL5 | 1 | 2 | 417 | 764 | 345 | 597 |
| | | 2 | 1 | 303 | 794 | 348 | 605 |
| MIP-3a | CCL20 | 1 | 2 | 1 | 14 | 4 | 36 |
| | | 2 | 2 | 0 | 45 | 2 | 91 |
| G-CSF | G-CSF | 1 | 0 | 4 | 124 | 25 | 233 |
| | | 2 | 0 | 3 | 197 | 12 | 224 |
| GM-CSF | GM-CSF | 1 | 69 | 63 | 508 | 66 | 866 |
| | | 2 | 44 | 49 | 392 | 62 | 886 |
| TGFb1 | TGFb1 | 1 | 4404 | 4094 | 7514 | 3944 | 9275 |
| | | 2 | 4537 | 4904 | 4037 | 4190 | 4187 |
| TNFa | TNFa | 1 | 49 | 52 | 124 | 43 | 86 |
| | | 2 | 54 | 48 | 153 | 48 | 87 |
| IFNg | IFNg | 1 | 0 | 0 | 21 | 0 | 0 |
| | | 2 | 0 | 0 | 0 | 0 | 0 |
| IL-1a | IL-1a | 1 | 0 | 22 | 61 | 51 | 140 |
| | | 2 | 0 | 0 | 56 | 49 | 133 |
| IL-36a | IL-36a | 1 | 0 | 5 | 6 | 2 | 4 |
| | | 2 | 0 | 0 | 0 | 0 | 0 |
| IL-6 | IL-6 | 1 | 5 | 113 | 4583 | 408 | 3050 |
| | | 2 | 11 | 68 | 3221 | 340 | 2151 |
| IL-20 | IL-20 | 1 | 0 | 0 | 0 | 0 | 19 |
| | | 2 | 0 | 0 | 0 | 0 | 0 |
| **ELISA conducted by UM Immunology Core (Measurements performed in duplicates)** | | | | | | | |
| GROa | CXCL-1 | 3 | 26 | 720 | 26652 | 1124 | 16796 |
| | | 4 | 101 | 860 | 30093 | 1406 | 20764 |
| | | 5 | 0 | 1284 | 22499 | 1275 | 18155 |
| GROb | CXCL-2 | 3 | 0 | 10 | 122 | 2 | 78 |

| | | | | | | | |
|---|---|---|---|---|---|---|---|
| | | 4 | 0 | 0 | 230 | 0 | 154 |
| | | 5 | 1 | 0 | 244 | 0 | 160 |
| MIG | CXCL9 | 3 | 12 | 0 | 68 | 12 | 0 |
| | | 4 | 12 | 35 | 55 | 30 | 50 |

**Supplemental Table 4. Genes significantly upregulated at 6h after dual TPA TX in WT and Krt17-/- mouse skin.**

| WT only (441) | Both genotypes (268) | *Krt17*$^{-/-}$ only (60) |
|---|---|---|
| Gm15511 | C920009B18Rik | A630035G10Rik |
| Gm47043 | ENSMUSG00000120024.1 | Gm23686 |
| Ankrd66 | Csf1r-ps | Myrfl |
| ENSMUSG00000120851.1 | Lilrb4b | Gm15433 |
| Slc2a6 | Nlrp12 | Gm37359 |
| Ccl6 | Ankk1 | Gm26143 |
| ENSMUSG00000121264.1 | Pla2g4d | S100a2 |
| Angptl4 | 1700112H15Rik | Fosl1 |
| Gm37113 | Treml2 | C2cd4b |
| Fcgr3 | Stfa2 | Pla2g4e |
| Gm13400 | Mir511 | Chil3 |
| Icos | ENSMUSG00000120542.1 | Cd5l |
| Siglecf | ENSMUSG00000120622.1 | Amn |
| Igsf6 | Ugt1a10 | Ighv1-64 |
| Sirpb1a | Gm7897 | Hbegf |
| Cyp7b1 | Gad1 | Gm5364 |
| Rbpj-ps3 | Tm4sf19 | Arc |
| Gm6566 | Krt6b | Gm16086 |
| Olfm4 | Gm2245 | Igf2bp1 |
| Gm49428 | Gpx2 | Gm10425 |
| Tigit | Ceacam10 | Fgf3 |
| Gm5650 | Tslp | Gm49035 |
| 4930459C07Rik | Sprr2a3 | Gm17173 |
| Cd8a | Htr7 | Dthd1 |
| ENSMUSG00000121276.1 | Tcf23 | Ptgs2 |
| Mmp8 | Prss27 | Gm30339 |
| Slc7a8 | Gm38335 | Uox |
| Tyrobp | Car4 | Gm29340 |
| Ly6a2 | Krt16 | Gm37567 |
| Gm29483 | Lilrb4a | Clec4a4 |
| Slc11a1 | Mroh4 | Wfdc12 |
| ENSMUSG00000119983.1 | B430306N03Rik | Tpo |
| Gm48786 | Mmp3 | 1700065I16Rik |
| Gm22061 | Gjb6 | Pspn |
| 4931406B18Rik | Slc36a3 | Sprr2j-ps |
| Slc17a6 | Slpi | Egr4 |
| Gm37645 | Tlr13 | ENSMUSG00000120878.1 |
| Omg | Chil4 | Gm47564 |
| Ubash3a | Tfec | Slc5a8 |
| Lin28a | Gm46224 | Fxyd4 |
| Bst1 | Gm2965 | Sprr2i |
| Klk1b27 | Gm38275 | Ccr3 |
| Gm48106 | Ccl7 | Foxd4 |
| Gm37800 | Gm19345 | Syt6 |
| Gm26626 | Csta3 | Snora69 |
| E230032D23Rik | Il4ra | ENSMUSG00002076636.1 |
| Samsn1 | Ccl12 | Klk9 |
| Gm18342 | Nmrk2 | BC035044 |
| Gm15459 | Mmp1b | Gm22771 |

Gm50345
Evi2
Cd6
Gm11714
Coro1a
Pik3ap1
Cxcr4
Cd22
Serpinb1b
Gm38171
Gm47682
Gm5733
Anxa11os
Acat3
Tubb6
Mir146b
Sla
ENSMUSG00000120384.1
Mmp13
Gm28068
Lhx1
Ccr7
Gm12606
Gm19696
Pira1
Sycp2
Gsdme
Gm8116
G530011O06Rik
Ccl4
Plac8
ENSMUSG00000121099.1
5830432E09Rik
Gm12708
1700102H20Rik
Cd101
Gm44165
Pira12
Mrgpra2a
1700019L13Rik
Rgs1
Smox
Gm40645
Spink12
Clec4a2
ENSMUSG00002075978.1
Lrrc25
Batf
Lst1
Ifitm1
5430425K12Rik
Clec4a3
Gm40915
ENSMUSG00000120147.1
Itgb2l
Cd177
Gm49368
Sprr2e
Sprr2a1
2610528A11Rik
ENSMUSG00000120697.1
1700016P03Rik
4930579C12Rik
Gm2682
Csf3
Gpr65
Gzmb
C130050O18Rik
Sprr2a2
Gm38575
1200007C13Rik
Gm5149
Timp1
Klk6
ENSMUSG00000120488.1
Abcb11
Gsdmc
Cd300lb
Gm8307
Gm14010
Gm37168
4930430E12Rik
Msx3
Gm16096
Treml4
Atp6v0d2
Ddah1
Socs3
Clec4e
5830415G21Rik
Has3
Il10
Cass4
Rtn4rl2
Grm6
Mcpt1
Cd14
Gm20610
Lce3c
Gm5414
F7
Sprr2d
Nrg1
Gm35808
Csf3r
F10
Gm13546
Ccl2
Mcpt8
Gm2629
Ccl20
Gm47985
Halr1
Gm44814
Ighv8-10
Lncenc1
Grpr
Areg
Gm47128
Gm13645
Has1

Gm20234
Ms4a4c
Cnr2
Gm13821
Ccl17
Fcgr2b
Il4i1
Slc16a3
Gm4891
ENSMUSG00000120159.1
Gm28809
ENSMUSG00000121091.1
Gm49249
Tmprss11g
Mmp25
Gm49888
Gm45805
C5ar2
Gm1647
Klk14
Gm6201
Oasl2
Fpr1
Ifi209
AU019990
Tmem156
Siglecl1
Ccl8
B3gnt7
Tnfrsf26
Gm16315
Tlr1
Il13
Siglece
1600010M07Rik
A530030E21Rik
Gm44888
Cd33
Cebpe
ENSMUSG00000120937.1
Mgam
ENSMUSG00000120668.1
Gm37975
4833407H14Rik
Kcnh4
Arg2
Gm12250
Th
Gm17230
Pilra
Gm13822
Plek
Gm26576
Mrgpra2b
Ceacam16
Upp1
Gsdmc2
S100a8
Plaur
Fcgr1
Steap1
Ccr5
Slc13a3
Adam8
Gm47719
Stfa1
Cstdc6
AU022793
Dntt
Ms4a4a
Saa1
S100a9
Cdhr1
1700012B09Rik
ENSMUSG00000120067.1
Hk3
Gm6342
Klk13
Olr1
Lce3d
Lpo
Itgam
Clec7a
C78197
C3ar1
Gm6209
Havcr1
Gm568
Cxcr1
Gm37199
Antxrl
Gm13655
Gm25696
Mir155hg
Gm32515
U90926
Cxcr2
Vmn1r210
Spink7
Sell
Gm42602
Btbd17
Nppb
Tarm1
Gsdmc3
ENSMUSG00000121069.1
Gm37131
Abca13
Gm9458
Krt6a

Tnfsf4
Gpr15
Dpys
ENSMUSG00000120744.1
Gm39363
ENSMUSG00000121234.1
Raet1d
Jakmip1
Pik3r5
Ncf2
B3gnt5
Abcc8
Cytip
Csf2rb
Ppp1r3b
Fcgr4
Tnip3
Gm28192
Gm12596
Gm43123
Gm30717
Klra2
ENSMUSG00000120077.1
Tbxas1
Ccr2
Hrh2
G930009F23Rik
P4ha3
Col6a4
Tnfrsf1b
4933412O06Rik
Gm29687
Gm13293
E230025N22Rik
Wfdc17
AI504432
ENSMUSG00000120666.1
Gm44860
Mrc1
4732465J04Rik
Gm44776
D130058E05Rik
Raet1e
Cd300c2
Ccl3
Gm10309
Krt90
Nfe2
Cacna1b
C130026I21Rik
Adgrf1
Adgrg3
Trim30b
ENSMUSG00000120446.1
Gm15413
Cd80
Plekhg4
Gm16556
Gm18445
Pdcd1lg2
Slc7a11
Gml2
Gprc5a
Mir7678
Sh2d5
Lcn2
Saa2
Pgap6
Adamts4
Bpifb1
Fcrlb
Ccl9
Gm8459
Chil1
Gm48065
ENSMUSG00000106319.2
Gm50419
Ankrd1
Ch25h
Pom121l2
Trpm2
Gm19510
Gm16549
Ms4a6d
ENSMUSG00000120676.1
Gm7343
Epgn
Gm47567
Psca
Btf3-ps13
Klk1
Gsdmc4
Mirt1
Alox8
Cd300ld
C5ar1
Sprr2h
Spp1
Gm49928
Gjb2
Lce3e
Aldh1a3
Gm20625
Gm38237
Gpr39
Stat4
Nlrp3
Atp1a3
Hsh2d
A630001O12Rik

ENSMUSG00002076074.1
Hand1
Zbp1
Cd300ld2
Hck
Glipr2
Tal2
Phlda2
Spatc1
E230016K23Rik
Slfn4
Cd300c
ENSMUSG00000120495.1
ENSMUSG00000120328.1
Ms4a6c
Gml
Bcl2a1b
Gm49300
Gm13657
Dok2
ENSMUSG00000121272.1
Gm49339
AB124611
Gm8080
ENSMUSG00000120467.1
Fgr
Hcls1
ENSMUSG00000120333.1
Foxf1
Gm18787
Fcrl1
Rab44
A530032D15Rik
1810073O08Rik
ENSMUSG00000120700.1
Clec4b2
Spef1l
Snx20
Akr1b8
Ly6g
Gapt
Pilrb1
Gm39321
Slc15a3
Morrbid
Gm6377
Gm43661
ENSMUSG00000120511.1
F630028O10Rik
Sros1
Ifi214
Oas1a
Gm43181
B230303A05Rik
Selp
Osm
Ccr1
Gm11597
Mmp12
Cntn2
Saa3
Fgf23
Il18rap
Gad1-ps
Il6
Cyp4f18
Tnfsf14
Hdc
Gm43834
Il1b
Gpr151
Cd300lf
Gm29686
Slfn1
Il19
Gdf3
ENSMUSG00000120223.1
Sprr2g
Htr1b
Bcl3
Bcl2a1a
Gm13068
Stfa3
Msr1
Srgn
Ocstamp
ENSMUSG00000120356.1
Sprr1b
Elfn1
Il24
Ptx3
Cd200r1
Gm21011
Rbp2
Gm39459
Dcstamp
Sirpb1c
Vmn1r198
Mefv
Tnfrsf9
Gm26579
Gm9733
Nt5c1a
Cxcl1
Gm37498
Cstdc5
Rnd1
Sprr2f
Arg1

Myo1f
Mirt2
Gm44659
Slfn10-ps
Ptger2
Aldh1a2
Clec4n
Lilra6
Gm20400
F5
1700011B04Rik
Gal
Smim45
2210406H18Rik
Clec4d
Eomes
Gm43215
Slc28a2
Pira13
Hmga2
Cd53
Gm10521
Gm46355
Gm12407
Slc39a14
Ripor2
Ctse
Cd300a
Ear6
AA467197
ENSMUSG00000120136.1
Gm19705
Gm48761
Vav1
Igfl3
Itgax
H60b
A530058O07Rik
Entpd3
Slamf6
Basp1
Cyth4
Rac2
ENSMUSG00000120115.1
Gm46189
Gm37703
Gm28981
Itgb2
Gm15821
Sfmbt2
Gm43660
Tmem163
Gpr35
Clec5a
ENSMUSG00002075844.1

Gfi1b
ENSMUSG00000120804.1
Vmn2r96
Gm31718
Amer2
Trim30c
Gpr84
Prok2
4933421A08Rik
Spib
Gm44861
Serpine1
Rptoros
Fpr2
ENSMUSG00000121163.1
Gm15956
Il1rl1
Syk
Trem1
Fcrla
Ntrk1
Gvin3
S1pr3
Gm38070
Platr7
Gm47015
Dnmt3l
Apon
Trem3
Tlr6
Themis2
Ly6c2
Mir142b
Pdpn
Muc20
Milr1
Pilrb2
Foxp3
Gm12047
Zfp951
4933432I03Rik
Gm38244
Soat2
Gm5150
ENSMUSG00000120408.1
Fcer1g
Relt
Il27
Gm42793
Lcp2
Mmp9
Runx2os2
Ngp
2810403G07Rik
Gm35147

2610028E06Rik
E230014E18Rik
Cpn1
Rps18-ps2
Trim30d
Lair1
Adm2
Gm15247
ENSMUSG00000121051.1
Mir5104
Gm47148
Gm36287
ENSMUSG00000120964.1
Glipr1
2610318N02Rik
Mcemp1
Gm20100
Gm34589
Dhrs9
Gm30085
A630081D01Rik
A530046M15Rik
Itgb3
Mybpc3
AI839979
Gm18271
Nckap1l
Tnc
Tmem179
1700001K23Rik
Gm8013
Ccrl2
Clec12a
ENSMUSG00000120457.1
Mpeg1
4930445E18Rik
Stx11
Lcp1
Gm34086
Gm26535
Gm37691
Gm34983
Gm10184
Emilin2
Gm34425
Ppbp
Scimp
Gm33370
Dppa3
Mir142hg
Il21r
Gm39556
Slc36a3os
Pira2
Slfn2

Mboat4
Olfr372
Tlr8
Gm38062
Fgf5
Gm29282
Cd84

**Supplemental Table 5. Genes significantly downregulated at 6h after dual TPA TX in WT and Krt17-/- mouse skin.**

| WT only (199) | Both genotypes (23) | *Krt17*$^{-/-}$ only (21) |
|---|---|---|
| Car14 | Tcap | Ubxn10 |
| AI838599 | Nlgn1 | Cntnap2 |
| Scgb1a1 | Foxo6os | Gbp10 |
| Lce1m | ENSMUSG00000120743.1 | Cabp7 |
| Gm16758 | Gng13 | Upk3b |
| Cacng6 | Prr32 | Gm10024 |
| Rpl19-ps3 | Bpifb2 | Mir6236 |
| Skint6 | Smco1 | Gm15564 |
| Lrrtm1 | Map3k7cl | Gm10649 |
| Hsd17b13 | Gm4265 | Mup5 |
| Grid2 | Tmem233 | Gm7001 |
| Plcd4 | Alkal2 | A330032B11Rik |
| 4930481A15Rik | Gm10318 | Cldn22 |
| Klhl38 | Gm34654 | Mrgprg |
| Cmtm5 | Frem3 | Tnfrsf17 |
| Azgp1 | Inmt | Ros1 |
| Igdcc3 | ENSMUSG00000120552.1 | D830013O20Rik |
| 2310016D23Rik | Hamp2 | Syt1 |
| Gpr165 | Gkn3 | Avp |
| Plekhd1 | Lrrc30 | 2310020H05Rik |
| Slitrk1 | Gabra1 | Gm17130 |
| Aoc3 | Gm18589 | |
| Igfn1 | Insyn2b | |
| Gm10100 | | |
| Hhip | | |
| Slc28a1 | | |
| Gsta3 | | |
| Krtap3-3 | | |
| Retn | | |
| Skint3 | | |
| Gm45218 | | |
| Adipoq | | |
| Clic6 | | |
| Mettl11b | | |
| Asb12 | | |
| Ky | | |
| Stmn3 | | |
| Sgo2b | | |
| 9430037G07Rik | | |
| Gm26703 | | |
| Gm14261 | | |

Acsm5
H2-M6-ps
Ibsp
Chaer1
Uckl1os
Gm34280
Klhl31
2610028H24Rik
Nell2
Tmem100
Gm15606
Gpr1
Hba-a2
Gpha2
Kcnh8
Lgi1
Nkain4
Nat8f3
Nnat
Dnah3
Adra2c
Skint9
Rlbp1
Gm47723
4930428N03Rik
Gm28286
Gm6939
Slc7a10
Adra1b
Methig1
Ccdc60
Mettl21e
Gm9885
2210020O09Rik
ENSMUSG00000120563.1
Gm48633
Zfp872
Trp53cor1
Pkhd1l1
Acsm1
Mt4
2900079G21Rik
Vsig4
Rimklb
Rab3b
Lrp2

Gm31305
Gm12224
Pdgfrl
Casq1
Frmpd4
Xkr9
Lrtm1
Gm19554
Lao1
Yipf7
Ces1f
Slc38a3
Gm37269
Doc2b
Bhmt
ENSMUSG00000120815.1
Kcnh6
Gm3822
Trpc4
Ankef1
Nrip2
Myl10
Gm38351
Chat
Asb10
Grik1
ENSMUSG00000120472.1
Gm10272
Gm9918
Stmnd1
Pbld1
Gm29183
Grm7
Rxfp1
C530044C16Rik
Gm26761
A530016L24Rik
Gm12818
Ostn
Klb
Hrk
Adad2
Mrgprb11-ps
2410137M14Rik
Ddit4l
Opcml
Serpina4-ps1
Scd3
Hemgn
Gfra3
Slc15a5

Lrrc38
Sfrp2
Ryr3
4930456G14Rik
Abca6
Rab11fip4os1
Mylk4
Gm2675
Hsd3b6
Gm15723
Gdf10
Sult1a1
Zfp648
Nhlh1
Atcayos
Cthrc1
Ccdc13
Kcnc2
Gm29681
Mrgprb3
1810064F22Rik
Ntng1
Skint4
1700009J07Rik
Otogl
Cfd
Prr33
Tbata
Gm20634
4930567K12Rik
C6
Klhl33
Actc1
Mrln
Cidec
Tspear
Gsta1
Ampd1
ENSMUSG00000120158.1
Mstn
8430422M14Rik
Asb18
Phkg1
Gm10113
Gm50222
Sox6os
Gm28609
6330403K07Rik
Krtap12-1
Gm15756
Krtap3-2
Scn4b
Trhde

Gm15222
Fcer2a
Gm38137
Gm11433
Gm25596
Vmn2r29
Ms4a10
Pck1

**Supplemental Table 6. Identification of 254 human gene homologs corresponding to the K17-dependent TAR genes discovered in mice.**

| TAR Differential Gene | Key (Mouse ID) | Genes Homology (Based on Jackson Lab Homology) | | | | | |
|---|---|---|---|---|---|---|---|
| Ankrd66 | 44487028 | ANKRD66 | | | | | |
| Slc2a6 | 44481660 | SLC2A6 | | | | | |
| Ccl6 | 44476473 | CCL15 | CCL23 | | | | |
| Angptl4 | 44482178 | ANGPTL4 | | | | | |
| Fcgr3 | 44487954 | FCGR2A | | | | | |
| Icos | 44488213 | ICOS | | | | | |
| Siglecf | 44474956 | SIGLEC14 | SIGLEC5 | SIGLEC8 | | | |
| Igsf6 | 44486193 | IGSF6 | | | | | |
| Sirpb1a | 44490335 | SIRPB1 | SIRPG | | | | |
| Cyp7b1 | 44481455 | CYP7B1 | | | | | |
| Olfm4 | 44477616 | OLFM4 | | | | | |
| Tigit | 44490691 | TIGIT | | | | | |
| Cd8a | 44474414 | CD8A | | | | | |
| Mmp8 | 44489073 | MMP8 | | | | | |
| Slc7a8 | 44473175 | SLC7A8 | | | | | |
| Tyrobp | 44475700 | TYROBP | | | | | |
| Slc11a1 | 44472441 | SLC11A1 | | | | | |
| 4931406B18Rik | 44474953 | SIGLEC1 | SIGLECL1 | | | | |
| Slc17a6 | 44473349 | SLC17A6 | | | | | |
| Omg | 44477980 | OMG | | | | | |
| Ubash3a | 44490899 | UBASH3A | | | | | |
| Lin28a | 44475249 | LIN28A | | | | | |
| Bst1 | 44487162 | BST1 | | | | | |
| Klk1b27 | 44488403 | KLK1 | KLK2 | KLK3 | | | |
| Samsn1 | 44474217 | SAMSN1 | | | | | |
| Evi2 | 44487859 | EVI2B | | | | | |
| Cd6 | 44487435 | CD6 | | | | | |
| Coro1a | 44472435 | CORO1A | | | | | |
| Pik3ap1 | 44482382 | PIK3AP1 | | | | | |
| Cxcr4 | 44470930 | CXCR4 | | | | | |
| Cd22 | 44486087 | CD22 | | | | | |
| Serpinb1b | 44473518 | SERPINB1 | | | | | |
| Acat3 | 44470963 | ACAT2 | | | | | |
| Tubb6 | 44483096 | TUBB6 | | | | | |
| Mir146b | 44488792 | MIR146B | | | | | |
| Sla | 44480818 | SLA | | | | | |
| Mmp13 | 44476356 | MMP13 | | | | | |
| Lhx1 | 44471387 | LHX1 | | | | | |
| Ccr7 | 44480972 | CCR7 | | | | | |
| Pira1 | 44488606 | LILRB3 | | | | | |
| Sycp2 | 44481763 | SYCP2 | | | | | |
| Gsdme | 44472853 | GSDME | | | | | |
| Ccl4 | 44476475 | CCL4 | CCL4L1 | CCL4L2 | | | |
| Plac8 | 44477848 | PLAC8 | | | | | |
| Cd101 | 44487417 | CD101 | | | | | |
| Pira12 | 44488602 | LILRA1 | LILRA2 | LILRA3 | LILRA4 | LILRA6 | LILRB3 |
| Mrgpra2a | 44489105 | MRGPRX3 | MRGPRX4 | | | | |
| Rgs1 | 44481380 | RGS1 | | | | | |
| Smox | 44478981 | SMOX | | | | | |

| | | | | | |
|---|---|---|---|---|---|
| Clec4a2 | 44481465 | CLEC4A | | | |
| Lrrc25 | 44488635 | LRRC25 | | | |
| Batf | 44474589 | BATF | | | |
| Lst1 | 44488660 | LST1 | | | |
| Ifitm1 | 44485954 | IFITM1 | | | |
| Clec4a3 | 44481752 | CLEC4A | | | |
| Itgb2l | 44485180 | ITGB2 | | | |
| Cnr2 | 44472892 | CNR2 | | | |
| Ccl17 | 44484745 | CCL17 | | | |
| Fcgr2b | 44487953 | FCGR2A | FCGR2B | FCGR2C | |
| Il4i1 | 44486347 | IL4I1 | | | |
| Slc16a3 | 44478550 | SLC16A3 | | | |
| Tmprss11g | 44490767 | TMPRSS11GP | | | |
| Mmp25 | 44477990 | MMP25 | | | |
| C5ar2 | 44487290 | C5AR2 | | | |
| Klk14 | 44488420 | KLK14 | | | |
| Oasl2 | 44489309 | OASL2P | | | |
| Fpr1 | 44475160 | FPR1 | | | |
| Ifi209 | 44488221 | IFI16 | | | |
| Tmem156 | 44490713 | TMEM156 | | | |
| Siglecl1 | 44490332 | SIGLECL1 | | | |
| Ccl8 | 44474377 | CCL8 | | | |
| B3gnt7 | 44472403 | B3GNT7 | | | |
| Tlr1 | 44472493 | TLR1 | | | |
| Il13 | 44477197 | IL13 | | | |
| Siglece | 44490331 | SIGLEC12 | SIGLEC7 | SIGLEC8 | SIGLEC9 |
| Cd33 | 44474952 | CD33 | SIGLEC6 | | |
| Cebpe | 44471816 | CEBPE | | | |
| Mgam | 44482762 | MGAM | | | |
| Kcnh4 | 44482754 | KCNH4 | | | |
| Arg2 | 44473335 | ARG2 | | | |
| Th | 44471188 | TH | | | |
| Pilra | 44489880 | PILRA | PILRB | | |
| Plek | 44484360 | PLEK | | | |
| Mrgpra2b | 44489107 | MRGPRX3 | MRGPRX4 | | |
| Ceacam16 | 44487463 | CEACAM16 | | | |
| Tnfsf4 | 44490775 | TNFSF4 | | | |
| Gpr15 | 44488076 | GPR15 | | | |
| Dpys | 44475696 | DPYS | | | |
| Raet1d | 44480591 | RAET1G | RAET1L | ULBP2 | |
| Jakmip1 | 44477346 | JAKMIP1 | | | |
| Pik3r5 | 44483351 | PIK3R5 | | | |
| Ncf2 | 44476887 | NCF2 | | | |
| B3gnt5 | 44472313 | B3GNT5 | | | |
| Abcc8 | 44474665 | ABCC8 | | | |
| Cytip | 44482929 | CYTIP | | | |
| Csf2rb | 44476114 | CSF2RB | | | |
| Ppp1r3b | 44483562 | PPP1R3B | | | |
| Fcgr4 | 44481188 | FCGR3A | FCGR3B | | |
| Tnip3 | 44490778 | TNIP3 | | | |
| Tbxas1 | 44475231 | TBXAS1 | | | |
| Ccr2 | 44476706 | CCR2 | | | |
| Hrh2 | 44475726 | HRH2 | | | |
| P4ha3 | 44484167 | P4HA3 | | | |
| Col6a4 | 44480234 | COL6A4P1 | COL6A4P2 | | |

| | | | | | | | |
|---|---|---|---|---|---|---|---|
| Tnfrsf1b | 44485153 | TNFRSF1B | | | | | |
| Mrc1 | 44476821 | MRC1 | | | | | |
| Raet1e | 44480584 | RAET1G | RAET1L | | | | |
| Cd300c2 | 44482454 | CD300C | | | | | |
| Ccl3 | 44476476 | CCL18 | CCL3 | CCL3L1 | CCL3L3 | | |
| Krt90 | 44470832 | KRT90P | | | | | |
| Nfe2 | 44471922 | NFE2 | | | | | |
| Cacna1b | 44476734 | CACNA1B | | | | | |
| Adgrf1 | 44474066 | ADGRF1 | | | | | |
| Adgrg3 | 44482939 | ADGRG3 | | | | | |
| Trim30b | 44490856 | TRIM5 | | | | | |
| Hand1 | 44488151 | HAND1 | | | | | |
| Zbp1 | 44491010 | ZBP1 | | | | | |
| Cd300ld2 | 44482449 | CD300LD | | | | | |
| Hck | 44481792 | HCK | | | | | |
| Glipr2 | 44480193 | GLIPR2 | | | | | |
| Tal2 | 44472412 | TAL2 | | | | | |
| Phlda2 | 44483372 | PHLDA2 | | | | | |
| Spatc1 | 44490499 | SPATC1 | | | | | |
| Slfn4 | 44490393 | SLFN12 | SLFN12L | | | | |
| Cd300c | 44482447 | CD300C | | | | | |
| Ms4a6c | 44477111 | MS4A6A | MS4A6E | | | | |
| Gml | 44488058 | GML | | | | | |
| Bcl2a1b | 44487123 | BCL2A1 | | | | | |
| Dok2 | 44476355 | DOK2 | | | | | |
| AB124611 | 44487218 | C19orf38 | | | | | |
| Fgr | 44472595 | FGR | | | | | |
| Hcls1 | 44475317 | HCLS1 | | | | | |
| Foxf1 | 44480635 | FOXF1 | | | | | |
| Fcrl1 | 44485213 | FCRL1 | | | | | |
| Rab44 | 44474396 | RAB44 | | | | | |
| Snx20 | 44477045 | SNX20 | | | | | |
| Akr1b8 | 44473991 | AKR1B10 | AKR1B15 | | | | |
| Gapt | 44488021 | GAPT | | | | | |
| Pilrb1 | 44489882 | PILRB | | | | | |
| Slc15a3 | 44490344 | SLC15A3 | | | | | |
| Morrbid | 44488964 | MIR4435-2HG | | | | | |
| Ifi214 | 44488220 | IFI16 | | | | | |
| Oas1a | 44489303 | OAS1 | | | | | |
| Selp | 44477858 | SELP | | | | | |
| Myo1f | 44482361 | MYO1F | | | | | |
| Ptger2 | 44474427 | PTGER2 | | | | | |
| Aldh1a2 | 44470708 | ALDH1A2 | | | | | |
| Clec4n | 44481464 | CLEC6A | | | | | |
| Lilra6 | 44488600 | LILRA1 | LILRA2 | LILRA3 | LILRA4 | LILRA6 | LILRB3 |
| F5 | 44472034 | F5 | | | | | |
| Gal | 44478121 | GAL | | | | | |
| Smim45 | 44490428 | SMIM45 | | | | | |
| Clec4d | 44487522 | CLEC4D | | | | | |
| Eomes | 44471436 | EOMES | | | | | |
| Slc28a2 | 44472609 | SLC28A2 | | | | | |
| Pira13 | 44488601 | LILRA1 | LILRA2 | LILRA3 | LILRA4 | LILRA6 | LILRB3 |
| Hmga2 | 44478386 | HMGA2 | | | | | |
| Cd53 | 44486576 | CD53 | | | | | |
| Slc39a14 | 44486660 | SLC39A14 | | | | | |

| | | | | |
|---|---|---|---|---|
| Ripor2 | 44475168 | RIPOR2 | | |
| Ctse | 44487604 | CTSE | | |
| Cd300a | 44482453 | CD300A | | |
| Ear6 | 44490175 | RNASE2 | RNASE3 | |
| AA467197 | 44486675 | C15orf48 | | |
| Vav1 | 44490941 | VAV1 | | |
| Igfl3 | 44488237 | IGFL2 | IGFL3 | IGFL4 |
| Itgax | 44482035 | ITGAX | | |
| Entpd3 | 44476075 | ENTPD3 | | |
| Slamf6 | 44486058 | SLAMF6 | | |
| Basp1 | 44477652 | BASP1 | | |
| Cyth4 | 44476753 | CYTH4 | | |
| Rac2 | 44473964 | RAC2 | | |
| Itgb2 | 44485181 | ITGB2 | | |
| Sfmbt2 | 44474634 | SFMBT2 | | |
| Tmem163 | 44472629 | TMEM163 | | |
| Gpr35 | 44477908 | GPR35 | | |
| Clec5a | 44487525 | CLEC5A | | |
| Gfi1b | 44482396 | GFI1B | | |
| Amer2 | 44482311 | AMER2 | | |
| Trim30c | 44477812 | TRIM5 | | |
| Gpr84 | 44484275 | GPR84 | | |
| Prok2 | 44477196 | PROK2 | | |
| Spib | 44490510 | SPIB | | |
| Serpine1 | 44481492 | SERPINE1 | | |
| Fpr2 | 44475163 | FPR2 | | |
| Il1rl1 | 44474828 | IL1RL1 | | |
| Syk | 44472895 | SYK | | |
| Trem1 | 44490838 | TREM1 | | |
| Fcrla | 44486768 | FCRLA | | |
| Ntrk1 | 44470709 | NTRK1 | | |
| S1pr3 | 44475327 | S1PR3 | | |
| Dnmt3l | 44471287 | DNMT3L | | |
| Apon | 44487050 | APONP | | |
| Tlr6 | 44472492 | TLR6 | | |
| Themis2 | 44485183 | THEMIS2 | | |
| Pdpn | 44489856 | PDPN | | |
| Muc20 | 44489178 | MUC20 | | |
| Milr1 | 44488734 | MILR1 | | |
| Pilrb2 | 44489881 | PILRA | PILRB | |
| Foxp3 | 44475008 | FOXP3 | | |
| Soat2 | 44481190 | SOAT2 | | |
| Fcer1g | 44475701 | FCER1G | | |
| Relt | 44483632 | RELT | | |
| Il27 | 44488289 | IL27 | | |
| Lcp2 | 44481713 | LCP2 | | |
| Mmp9 | 44475238 | MMP9 | | |
| Cpn1 | 44474879 | CPN1 | | |
| Trim30d | 44477822 | TRIM5 | | |
| Lair1 | 44488555 | LAIR1 | LAIR2 | |
| Adm2 | 44475299 | ADM2 | | |
| Glipr1 | 44477414 | GLIPR1 | | |
| Mcemp1 | 44488713 | MCEMP1 | | |
| Dhrs9 | 44474904 | DHRS9 | | |
| Itgb3 | 44476047 | ITGB3 | | |

| | | | | | | | |
|---|---|---|---|---|---|---|---|
| Mybpc3 | 44481278 | MYBPC3 | | | | | |
| Nckap1l | 44486477 | NCKAP1L | | | | | |
| Tnc | 44471085 | TNC | | | | | |
| Tmem179 | 44480025 | TMEM179 | | | | | |
| Ccrl2 | 44487415 | CCRL2 | | | | | |
| Clec12a | 44487514 | CLEC12A | | | | | |
| Mpeg1 | 44477893 | MPEG1 | | | | | |
| Stx11 | 44479452 | STX11 | | | | | |
| Lcp1 | 44471178 | LCP1 | | | | | |
| Emilin2 | 44476249 | EMILIN2 | | | | | |
| Ppbp | 44486150 | PPBP | | | | | |
| Scimp | 44490267 | SCIMP | | | | | |
| Dppa3 | 44487766 | DPPA3 | | | | | |
| Il21r | 44476107 | IL21R | | | | | |
| Pira2 | 44488599 | LILRA1 | LILRA2 | LILRA3 | LILRA4 | LILRA6 | LILRB3 |
| Slfn2 | 44490395 | SLFN12 | SLFN12L | | | | |
| Mboat4 | 44476712 | MBOAT4 | | | | | |
| Tlr8 | 44472489 | TLR8 | | | | | |
| Fgf5 | 44473526 | FGF5 | | | | | |
| Cd84 | 44486057 | CD84 | | | | | |

**Supplemental Table 7. Identification of TAR12 transcriptomic signature.**

| 209 genes occur in PSOR dataset | Average expression | | | | | KC>Myeloid AND KC>0.1? |
|---|---|---|---|---|---|---|
| | Melanocytes | Keratinocytes | KRT23_MMP7 | T-Cells | Myeloid Lineage | |
| ANGPTL4 | 0.063656 | 0.290166 | 0.152302 | 0.193288 | 0.056211 | yes |
| IFITM1 | 0.306694 | 0.150862 | 0.042319 | 0.189995 | 0.048363 | yes |
| IFI16 | 1.299224 | 1.184575 | 0.747518 | 1.371006 | 0.945852 | yes |
| PHLDA2 | 0.540415 | 1.060991 | 0.849200 | 0.669865 | 0.653782 | yes |
| KLK1 | 0.012366 | 0.102685 | 0.089409 | 0.028413 | 0.016692 | yes |
| PDPN | 0.020386 | 0.109907 | 0.033672 | 0.082095 | 0.028818 | yes |
| AKR1B10 | 0.075856 | 0.554196 | 0.146836 | 0.296131 | 0.110013 | yes |
| ACAT2 | 0.236885 | 0.292024 | 0.231847 | 0.313133 | 0.206577 | yes |
| TUBB6 | 0.344928 | 0.549359 | 0.354967 | 0.324203 | 0.378425 | yes |
| OAS1 | 0.302142 | 0.320890 | 0.134855 | 0.246144 | 0.083924 | yes |
| HRH2 | 0.184349 | 0.172061 | 0.023017 | 0.108659 | 0.077164 | yes |
| C19orf48 | 0.218125 | 0.524632 | 0.319385 | 0.404773 | 0.198777 | yes |
| SLC2A6 | 0.040241 | 0.012448 | 0.011916 | 0.003312 | 0.026179 | no |
| BATF | 0.073984 | 0.002628 | 0.003187 | 0.687662 | 0.094743 | no |
| TNFRSF1B | 0.001580 | 0.002080 | 0.001494 | 0.167092 | 0.197921 | no |
| IGFL2 | 0.002961 | 0.020905 | 0.025000 | 0.013939 | 0.006703 | no |
| LCP1 | 0.020358 | 0.005961 | 0.009037 | 0.757721 | 1.901677 | no |
| LST1 | 0.015065 | 0.007208 | 0.004081 | 0.089121 | 2.123160 | no |
| MRC1 | 0.001334 | 0.000109 | 0.000000 | 0.000000 | 0.141362 | no |
| IGFL3 | 0.003311 | 0.012716 | 0.194152 | 0.014603 | 0.002408 | no |
| EMILIN2 | 0.032736 | 0.002750 | 0.001338 | 0.015836 | 0.233203 | no |
| FCGR2A | 0.049883 | 0.002685 | 0.002576 | 0.001109 | 0.658526 | no |
| CD300C | 0.000000 | 0.000108 | 0.000000 | 0.001299 | 0.103100 | no |
| IGFL4 | 0.001716 | 0.009274 | 0.134423 | 0.002636 | 0.002512 | no |
| SCIMP | 0.001546 | 0.000885 | 0.001035 | 0.000000 | 0.007684 | no |
| ICOS | 0.001565 | 0.000456 | 0.000433 | 0.152430 | 0.001931 | no |
| ITGB2 | 0.009081 | 0.002493 | 0.086338 | 0.250265 | 0.853788 | no |
| CCL18 | 0.603732 | 0.001615 | 0.000899 | 0.003883 | 0.002868 | no |
| ITGAX | 0.001745 | 0.000104 | 0.000691 | 0.001428 | 0.094058 | no |
| IL21R | 0.001059 | 0.000527 | 0.001106 | 0.080616 | 0.053668 | no |
| SIGLEC14 | 0.000245 | 0.000000 | 0.000000 | 0.000000 | 0.010666 | no |
| CNR2 | 0.000000 | 0.000518 | 0.000000 | 0.003503 | 0.002124 | no |
| CCL3 | 0.175055 | 0.004647 | 0.002520 | 0.425071 | 0.326740 | no |
| ENTPD3 | 0.005572 | 0.019361 | 0.066583 | 0.016726 | 0.004260 | no |
| TLR8 | 0.000211 | 0.000057 | 0.000000 | 0.000000 | 0.046120 | no |
| SIGLEC5 | 0.000000 | 0.000020 | 0.000000 | 0.000000 | 0.005168 | no |
| CCL17 | 0.006970 | 0.002095 | 0.000000 | 0.003247 | 0.255465 | no |
| CCL3L3 | 0.001880 | 0.001888 | 0.001476 | 0.060868 | 0.143591 | no |
| SLAMF6 | 0.000000 | 0.000252 | 0.000000 | 0.031554 | 0.001458 | no |
| CD84 | 0.001160 | 0.000766 | 0.003112 | 0.163293 | 0.191475 | no |
| IGSF6 | 0.004885 | 0.001805 | 0.001766 | 0.003393 | 1.108868 | no |
| FCGR2B | 0.041634 | 0.001491 | 0.001176 | 0.000000 | 0.723027 | no |
| NFE2 | 0.000431 | 0.001007 | 0.000707 | 0.002124 | 0.000000 | no |
| BASP1 | 0.098910 | 0.107886 | 0.036414 | 0.120950 | 0.780326 | no |
| SIRPB1 | 0.000000 | 0.000608 | 0.000000 | 0.010182 | 0.015906 | no |
| IL4I1 | 0.011452 | 0.002586 | 0.004828 | 0.055622 | 0.568947 | no |
| CACNA1B | 0.000534 | 0.000115 | 0.000688 | 0.000958 | 0.000552 | no |
| CYTH4 | 0.001942 | 0.000749 | 0.001508 | 0.160194 | 0.128167 | no |

| | | | | | | |
|---|---|---|---|---|---|---|
| SIRPG | 0.001857 | 0.000448 | 0.001931 | 0.168537 | 0.000950 | no |
| SLC16A3 | 0.005276 | 0.002916 | 0.005018 | 0.023985 | 0.353499 | no |
| ADGRF1 | 0.005690 | 0.034269 | 0.003210 | 0.025687 | 0.006114 | no |
| RAC2 | 0.013337 | 0.046582 | 0.013017 | 0.921680 | 0.237275 | no |
| CYP7B1 | 0.026038 | 0.075246 | 0.099774 | 0.042930 | 0.012355 | no |
| MMP25 | 0.002490 | 0.001409 | 0.000000 | 0.005907 | 0.023731 | no |
| ADGRG3 | 0.001988 | 0.000714 | 0.000747 | 0.000000 | 0.000000 | no |
| SFMBT2 | 0.019987 | 0.001330 | 0.001736 | 0.065258 | 0.074803 | no |
| OLFM4 | 0.000000 | 0.000133 | 0.000000 | 0.000000 | 0.000000 | no |
| C5AR2 | 0.000739 | 0.001464 | 0.007497 | 0.000000 | 0.018364 | no |
| TRIM5 | 0.055603 | 0.048084 | 0.043004 | 0.058966 | 0.039760 | no |
| TMEM163 | 0.000000 | 0.000487 | 0.000000 | 0.000000 | 0.010256 | no |
| TIGIT | 0.000270 | 0.000508 | 0.001152 | 0.239227 | 0.003076 | no |
| FPR1 | 0.000720 | 0.000366 | 0.000713 | 0.000000 | 0.421170 | no |
| HAND1 | 0.002103 | 0.002054 | 0.002698 | 0.005954 | 0.000000 | no |
| GPR35 | 0.000665 | 0.000532 | 0.001359 | 0.004728 | 0.077411 | no |
| CD8A | 0.000876 | 0.001032 | 0.000996 | 0.404190 | 0.003112 | no |
| ZBP1 | 0.000000 | 0.001437 | 0.000000 | 0.033055 | 0.011784 | no |
| CLEC5A | 0.000884 | 0.000843 | 0.001345 | 0.003345 | 0.589383 | no |
| SLC7A8 | 0.312888 | 0.071154 | 0.025809 | 0.073953 | 0.017059 | no |
| TMEM156 | 0.000551 | 0.001150 | 0.000000 | 0.033747 | 0.041788 | no |
| HCK | 0.002725 | 0.006704 | 0.000000 | 0.003207 | 0.574200 | no |
| AMER2 | 0.131502 | 0.000118 | 0.001582 | 0.000000 | 0.001941 | no |
| TYROBP | 0.017831 | 0.008927 | 0.006992 | 0.040060 | 2.772929 | no |
| CCL8 | 0.001748 | 0.001328 | 0.000000 | 0.003125 | 0.017937 | no |
| GLIPR2 | 0.338385 | 0.003995 | 0.002397 | 0.180737 | 0.579193 | no |
| GPR84 | 0.001771 | 0.003286 | 0.001947 | 0.006341 | 0.162203 | no |
| SLC11A1 | 0.007153 | 0.008676 | 0.008373 | 0.008488 | 0.021141 | no |
| B3GNT7 | 0.005166 | 0.001540 | 0.003158 | 0.000000 | 0.047053 | no |
| TAL2 | 0.000000 | 0.000623 | 0.000000 | 0.000000 | 0.000558 | no |
| PROK2 | 0.000000 | 0.000099 | 0.000000 | 0.014779 | 0.001071 | no |
| SIGLEC1 | 0.000000 | 0.000061 | 0.000000 | 0.000000 | 0.008659 | no |
| TLR1 | 0.009949 | 0.003072 | 0.019024 | 0.008306 | 0.211884 | no |
| SPIB | 0.001202 | 0.002242 | 0.000493 | 0.001229 | 0.108690 | no |
| OMG | 0.010420 | 0.000134 | 0.000000 | 0.000000 | 0.001111 | no |
| IL13 | 0.000000 | 0.000343 | 0.000931 | 0.007673 | 0.000000 | no |
| SLFN12 | 0.037914 | 0.023007 | 0.027186 | 0.060506 | 0.060887 | no |
| SERPINE1 | 0.004675 | 0.026473 | 0.010304 | 0.022885 | 0.018585 | no |
| UBASH3A | 0.000678 | 0.000194 | 0.000000 | 0.080240 | 0.000000 | no |
| SIGLEC12 | 0.000000 | 0.000000 | 0.000000 | 0.000000 | 0.005634 | no |
| SLFN12L | 0.003297 | 0.002956 | 0.000723 | 0.072015 | 0.001854 | no |
| FPR2 | 0.000000 | 0.000066 | 0.000000 | 0.003311 | 0.021444 | no |
| LIN28A | 0.000000 | 0.000179 | 0.000000 | 0.000000 | 0.000000 | no |
| SIGLEC7 | 0.000000 | 0.000035 | 0.000000 | 0.000000 | 0.023830 | no |
| MS4A6A | 0.004371 | 0.004488 | 0.001162 | 0.001703 | 1.095284 | no |
| IL1RL1 | 0.000000 | 0.000231 | 0.000000 | 0.000000 | 0.001052 | no |
| BST1 | 0.176225 | 0.010492 | 0.003986 | 0.005007 | 0.031413 | no |
| SIGLEC9 | 0.000000 | 0.000056 | 0.000000 | 0.000000 | 0.066296 | no |
| GML | 0.000000 | 0.000167 | 0.000000 | 0.000000 | 0.000000 | no |
| SYK | 0.012461 | 0.076524 | 0.026827 | 0.055063 | 0.201715 | no |
| CD33 | 0.001546 | 0.000636 | 0.001122 | 0.000000 | 0.188271 | no |
| BCL2A1 | 0.522255 | 0.002219 | 0.001422 | 0.018984 | 0.545483 | no |
| TREM1 | 0.000000 | 0.001068 | 0.000000 | 0.001009 | 0.057799 | no |
| SAMSN1 | 0.004110 | 0.001362 | 0.000904 | 0.457050 | 0.452712 | no |

| | | | | | | |
|---|---|---|---|---|---|---|
| CEBPE | 0.000355 | 0.000332 | 0.000000 | 0.003332 | 0.003045 | no |
| DOK2 | 0.001529 | 0.001384 | 0.000692 | 0.237676 | 0.280423 | no |
| FCRLA | 0.400099 | 0.001191 | 0.000721 | 0.004523 | 0.006620 | no |
| EVI2B | 0.021058 | 0.001279 | 0.000628 | 0.276242 | 0.679486 | no |
| MGAM | 0.000000 | 0.000048 | 0.000000 | 0.000000 | 0.005890 | no |
| FGR | 0.002064 | 0.001605 | 0.000689 | 0.001387 | 0.204597 | no |
| S1PR3 | 0.004194 | 0.024597 | 0.083284 | 0.010667 | 0.003790 | no |
| CD6 | 0.001337 | 0.001003 | 0.001269 | 0.140282 | 0.017249 | no |
| KCNH4 | 0.000000 | 0.000155 | 0.000000 | 0.000000 | 0.000000 | no |
| HCLS1 | 0.005923 | 0.002216 | 0.001529 | 0.397953 | 0.705350 | no |
| TLR6 | 0.010945 | 0.003112 | 0.018735 | 0.007319 | 0.074016 | no |
| CORO1A | 0.015775 | 0.010156 | 0.015182 | 1.350055 | 1.206103 | no |
| ARG2 | 0.033249 | 0.092611 | 0.029812 | 0.077486 | 0.016712 | no |
| FOXF1 | 0.000000 | 0.000189 | 0.000000 | 0.000000 | 0.000000 | no |
| THEMIS2 | 0.115832 | 0.003020 | 0.006257 | 0.026628 | 0.188027 | no |
| PIK3AP1 | 0.000000 | 0.000207 | 0.000000 | 0.016742 | 0.046454 | no |
| TH | 0.001550 | 0.002193 | 0.000000 | 0.001324 | 0.000475 | no |
| RAB44 | 0.000000 | 0.002715 | 0.000000 | 0.000000 | 0.000391 | no |
| CXCR4 | 0.029059 | 0.097745 | 0.035172 | 0.726390 | 0.565115 | no |
| PILRA | 0.001628 | 0.004571 | 0.000646 | 0.004422 | 0.210075 | no |
| SNX20 | 0.002014 | 0.000602 | 0.000432 | 0.069921 | 0.272470 | no |
| MUC20 | 0.004078 | 0.008373 | 0.013028 | 0.017748 | 0.003037 | no |
| CD22 | 0.000000 | 0.000903 | 0.000000 | 0.002380 | 0.007235 | no |
| PILRB | 0.062860 | 0.091689 | 0.051701 | 0.087487 | 0.036956 | no |
| MILR1 | 0.001498 | 0.004931 | 0.001032 | 0.000000 | 0.085908 | no |
| SERPINB1 | 0.581097 | 0.315406 | 0.416017 | 0.400441 | 1.049928 | no |
| PLEK | 0.001810 | 0.001466 | 0.000000 | 0.005112 | 0.880668 | no |
| AKR1B15 | 0.003448 | 0.011156 | 0.010971 | 0.007985 | 0.000728 | no |
| FOXP3 | 0.000000 | 0.000724 | 0.000519 | 0.050092 | 0.000000 | no |
| MRGPRX3 | 0.000000 | 0.000458 | 0.000000 | 0.000000 | 0.000000 | no |
| GAPT | 0.000282 | 0.000218 | 0.000945 | 0.000000 | 0.197065 | no |
| SOAT2 | 0.005988 | 0.000596 | 0.000846 | 0.000000 | 0.000596 | no |
| CEACAM16 | 0.000000 | 0.000119 | 0.000000 | 0.000000 | 0.000000 | no |
| SLC15A3 | 0.005357 | 0.010631 | 0.004796 | 0.006257 | 0.052938 | no |
| FCER1G | 0.017129 | 0.007918 | 0.008623 | 0.030208 | 2.080826 | no |
| SLA | 0.003669 | 0.001885 | 0.002953 | 0.548100 | 0.445880 | no |
| TNFSF4 | 0.000000 | 0.000307 | 0.000892 | 0.036837 | 0.018855 | no |
| MIR4435-2HG | 0.018972 | 0.010709 | 0.133392 | 0.418377 | 0.196322 | no |
| RELT | 0.019007 | 0.024459 | 0.021172 | 0.028955 | 0.091318 | no |
| MMP13 | 0.000000 | 0.000071 | 0.020099 | 0.000000 | 0.001102 | no |
| GPR15 | 0.000000 | 0.000117 | 0.000000 | 0.002913 | 0.000000 | no |
| IL27 | 0.000000 | 0.000035 | 0.000000 | 0.000000 | 0.024988 | no |
| LHX1 | 0.000000 | 0.000392 | 0.000756 | 0.000000 | 0.000000 | no |
| RAET1G | 0.006475 | 0.060763 | 0.050267 | 0.030552 | 0.011297 | no |
| SELP | 0.000000 | 0.000043 | 0.000000 | 0.004449 | 0.001468 | no |
| LCP2 | 0.004412 | 0.001410 | 0.002857 | 0.167168 | 0.392781 | no |
| CCR7 | 0.000865 | 0.001099 | 0.000000 | 0.056447 | 0.278374 | no |
| RAET1L | 0.002337 | 0.016885 | 0.012260 | 0.007587 | 0.001524 | no |
| MYO1F | 0.015229 | 0.030520 | 0.025654 | 0.202385 | 0.235939 | no |
| MMP9 | 0.003151 | 0.002270 | 0.001246 | 0.001529 | 0.477654 | no |
| LILRB3 | 0.008669 | 0.017818 | 0.005553 | 0.020164 | 0.114682 | no |
| ULBP2 | 0.009202 | 0.034753 | 0.030987 | 0.027547 | 0.010615 | no |
| PTGER2 | 0.000000 | 0.000781 | 0.035291 | 0.008853 | 0.007119 | no |
| LAIR1 | 0.004366 | 0.002721 | 0.002650 | 0.022654 | 0.076990 | no |

| | | | | | | |
|---|---|---|---|---|---|---|
| SYCP2 | 0.011309 | 0.008238 | 0.041738 | 0.015313 | 0.007980 | no |
| JAKMIP1 | 0.000000 | 0.000080 | 0.000000 | 0.058136 | 0.001440 | no |
| ALDH1A2 | 0.000000 | 0.001977 | 0.003968 | 0.002662 | 0.000000 | no |
| LAIR2 | 0.004533 | 0.019314 | 0.002988 | 0.165871 | 0.008703 | no |
| CCL4 | 0.006166 | 0.005053 | 0.007154 | 0.528034 | 0.185821 | no |
| PIK3R5 | 0.001245 | 0.000157 | 0.000868 | 0.018655 | 0.067236 | no |
| CLEC6A | 0.000000 | 0.000000 | 0.000000 | 0.000000 | 0.004768 | no |
| ADM2 | 0.000713 | 0.004364 | 0.001354 | 0.009019 | 0.000000 | no |
| CCL4L2 | 0.002139 | 0.001209 | 0.000000 | 0.053799 | 0.161926 | no |
| NCF2 | 0.004697 | 0.020169 | 0.013788 | 0.013978 | 0.387988 | no |
| GAL | 0.006360 | 0.019543 | 0.008155 | 0.007126 | 0.003337 | no |
| GLIPR1 | 0.004962 | 0.007207 | 0.042249 | 0.234417 | 0.862489 | no |
| PLAC8 | 0.003288 | 0.000608 | 0.004869 | 0.011196 | 0.022201 | no |
| B3GNT5 | 0.039494 | 0.196646 | 0.090337 | 0.117327 | 0.303677 | no |
| CLEC4D | 0.000000 | 0.000051 | 0.000000 | 0.000000 | 0.014958 | no |
| MCEMP1 | 0.000000 | 0.000092 | 0.000000 | 0.000000 | 0.004921 | no |
| CD101 | 0.000627 | 0.004524 | 0.001777 | 0.020128 | 0.026149 | no |
| CYTIP | 0.005688 | 0.001715 | 0.001946 | 0.436558 | 0.621518 | no |
| EOMES | 0.002774 | 0.000183 | 0.000000 | 0.007849 | 0.000190 | no |
| DHRS9 | 0.000435 | 0.004272 | 0.000000 | 0.001861 | 0.102617 | no |
| LILRA1 | 0.000000 | 0.000060 | 0.000000 | 0.000000 | 0.014634 | no |
| CSF2RB | 0.000245 | 0.000475 | 0.000000 | 0.003677 | 0.261348 | no |
| SLC28A2 | 0.001710 | 0.001893 | 0.000633 | 0.004586 | 0.001223 | no |
| ITGB3 | 0.002439 | 0.000016 | 0.000000 | 0.000000 | 0.000000 | no |
| LILRA2 | 0.000000 | 0.000398 | 0.000000 | 0.000000 | 0.052607 | no |
| PPP1R3B | 0.022075 | 0.019125 | 0.046256 | 0.013694 | 0.024339 | no |
| CD53 | 0.006537 | 0.002613 | 0.002679 | 0.644066 | 0.892719 | no |
| MYBPC3 | 0.000000 | 0.000149 | 0.000000 | 0.000000 | 0.006798 | no |
| LILRA4 | 0.000000 | 0.000098 | 0.000000 | 0.000000 | 0.018367 | no |
| FCGR3A | 0.001280 | 0.000674 | 0.000000 | 0.008080 | 0.245835 | no |
| SLC39A14 | 0.074119 | 0.026459 | 0.001706 | 0.028039 | 0.015955 | no |
| NCKAP1L | 0.002696 | 0.001364 | 0.000549 | 0.144049 | 0.306880 | no |
| LILRA6 | 0.000919 | 0.000146 | 0.000000 | 0.000000 | 0.017799 | no |
| TNIP3 | 0.011462 | 0.080398 | 0.012038 | 0.182075 | 0.041583 | no |
| CTSE | 0.000000 | 0.000115 | 0.000000 | 0.000000 | 0.000814 | no |
| TNC | 0.009931 | 0.043351 | 0.146647 | 0.038245 | 0.011285 | no |
| RGS1 | 0.021753 | 0.010317 | 0.009665 | 0.855657 | 1.552819 | no |
| TBXAS1 | 0.019858 | 0.023314 | 0.003334 | 0.044975 | 0.079523 | no |
| CD300A | 0.000880 | 0.001729 | 0.003570 | 0.020393 | 0.261524 | no |
| CCRL2 | 0.000743 | 0.000138 | 0.000000 | 0.009163 | 0.067476 | no |
| SMOX | 0.071231 | 0.023287 | 0.022607 | 0.026187 | 0.015341 | no |
| CCR2 | 0.000000 | 0.000809 | 0.000000 | 0.010010 | 0.061981 | no |
| RNASE2 | 0.000000 | 0.000000 | 0.000000 | 0.000000 | 0.013860 | no |
| CLEC12A | 0.001602 | 0.000048 | 0.000000 | 0.008527 | 0.011682 | no |
| CLEC4A | 0.002592 | 0.002826 | 0.003904 | 0.001292 | 0.491352 | no |
| VAV1 | 0.000991 | 0.000703 | 0.001411 | 0.059543 | 0.080119 | no |
| MPEG1 | 0.000000 | 0.000324 | 0.000651 | 0.000000 | 0.193452 | no |
| LRRC25 | 0.000000 | 0.000355 | 0.000000 | 0.000000 | 0.043470 | no |
| P4HA3 | 0.000000 | 0.001471 | 0.001925 | 0.003252 | 0.001307 | no |
| STX11 | 0.117859 | 0.053419 | 0.088516 | 0.076279 | 0.173052 | no |
| F5 | 0.000000 | 0.000259 | 0.000629 | 0.039484 | 0.001393 | no |
| HMGA2 | 0.000000 | 0.002005 | 0.000648 | 0.001178 | 0.000000 | no |
| C19orf38 | 0.001444 | 0.001215 | 0.000000 | 0.002418 | 0.093953 | no |
